# Cortical localization and dynamics of elementary mathematical concepts

**DOI:** 10.64898/2026.04.30.721827

**Authors:** Samuel Debray, Alireza Karami, Daniela Valério, Maxime Cauté, Christophe Pallier, Stanislas Dehaene

**Author notes:** Corresponding Author: Samuel Debray, Alireza Karami, Stanislas Dehaene. These authors contributed equally to this work (Samuel Debray, Alireza Karami).

## Abstract

How the brain encodes abstract concepts remains poorly understood. Current theories propose that, in brains and computers alike, word meanings are represented by vectors of neural activation whose similarities reflect semantic relationships. Here, we tested whether this hypothesis also applies to abstract concepts of elementary mathematics. We collected behavioral, 7 Tesla functional MRI and magneto-encephalography (MEG) data and used representational similarity analysis to ask where, when and how fifteen concepts of integers, fractions, and geometric shapes are encoded in the adult brain. Behavioral similarity ratings revealed a rich conceptual structure characterized by both categorical distinctions (numbers vs shapes, integers vs fractions), a numerical distance effect for integers, and systematic correspondences between items involving the same number (e.g. three, third, triangle). Functional MRI identified a bilateral cortical network whose neural encodings of concepts correlated with their semantic similarity, overlapping with classic math-responsive regions and encompassing IPS and ITG as well as dorsolateral prefrontal cortex (dlPFC). A double dissociation was observed, with a preference for arithmetic in the right anterior intraparietal sulcus (IPS), and for geometry in left inferior temporal gyrus (ITG) and bilateral posterior IPS. MEG revealed that a semantic neural code common to written words and symbols is activated by about 230 ms, again primarily distinguishing integers, fractions and geometry concepts. Together, these findings suggest that mathematical concepts are organized in the brain along both categorical and numerical dimensions, with overlapping but partially distinct sites supporting arithmetic and geometry domains.

## Introduction

A central goal of cognitive neuroscience is to understand how concepts are represented in the human brain. In both cognitive science and artificial neural networks, an increasingly accepted theory is that word meanings can be represented geometrically as distributed embeddings in high-dimensional representational spaces (Landauer and Dumais 1997; Mikolov et al. 2013): the meaning of each word is represented by a vector, and the distance between those vectors reflect the semantic similarity of the corresponding concepts. The goal of the present work is to test whether this hypothesis remains true for abstract words, and specifically within the semantic domain of elementary mathematical concepts.

Over the past decade, a growing body of research has begun to map the neural architecture underlying conceptual word knowledge, revealing distributed and systematic representations across cortical regions and their timing (e.g. Borghesani et al. 2019; Borghesani and Piazza 2017; Goldstein et al. 2022, 2024; Tong et al. 2022). Using naturalistic language paradigms, Huth et al. (2016) provided a comprehensive cortical map of semantic representations by having participants listen to hours of narrative stories while undergoing functional magnetic resonance imaging (fMRI). They showed that distinct cortical regions are selectively responsive to broad categories of concepts, from social and visual domains to abstract knowledge, including numbers. This work and related ones established a powerful framework for linking human concepts to distributed neural activation patterns (Caucheteux, Gramfort, and King 2022; Deniz et al. 2019; Pasquiou et al. 2023; Pereira et al. 2018; Popham et al. 2021; Saha et al. 2025; Tong et al. 2022; Vargas and Just 2022).

Several studies further probed the geometry of cortical semantic spaces. Nelli et al. (2023) demonstrated that new items can be flexibly embedded within semantic spaces: after participants learned to order previously unknown objects along an arbitrary scale, the neural representations in posterior parietal and dorsomedial prefrontal cortices reflected this learned ordering. Moreover, when two independently trained scales were later merged through comparative learning, the neural representations of the two object groups collapsed onto a unified dimension, illustrating the brain’s ability to restructure representational space according to newly acquired relations. In a related set of studies, Viganò et al. (2021; 2021; 2020) identified neural substrates supporting navigation within a two-dimensional semantic space. Their results revealed that both distance and direction (both relative and absolute) in an abstract conceptual space recruit the dorsolateral prefrontal cortex (dlPFC) and the right entorhinal cortex (EHC), thus linking mechanisms of spatial navigation and grid-cell coding to semantic cognition. Together, these findings support the hypothesis that the brain encodes conceptual relations using structured semantic maps that allow generalization and flexible inference (Bellmund et al. 2018; Gärdenfors 2000).

While these studies provide broad insights into how a few major dimensions of meaning are represented across the cortex, other studies have focused on the analysis of specific linguistic or conceptual domains. Domain-specific investigations have shown that distinct semantic categories such as abstract words (Kaiser, Jacobs, and Cichy 2022), objects (Almeida et al. 2023; Bonner and Epstein 2021), or physics concepts (Mason and Just 2016; Mason, Schumacher, and Just 2021) can recruit specialized cortical regions. Modality-independent conceptual representations have also been characterized. For example, Dirani & Pylkkänen (2024) found using MEG that conceptual representations common to pictures and words contain both semantic and visual features. Likewise, Bezsudnova et al. (2024) showed that category information in MEG data could be decoded from pictures and words between ∼150–430 ms, whereas Giari et al. (2020) reported early (∼105 ms) conceptual information for pictures and later (∼230–335 ms) for words. These studies suggest that conceptual category structure can be rapidly accessed across modalities. Using fMRI, Popham et al. (2021) found that visual and linguistic semantic representations are aligned along the border of visual cortex, with the same semantic categories represented in adjacent visual and language maps, consistent with the idea that semantic structure can be preserved across different input formats. Here, our goal is to extend this line of research to the domain of mathematics, focusing on the cortical representation of elementary mathematical concepts, and asking whether they exhibit systematic relationships among related arithmetic and geometric concepts.

### Previous research on the representation of abstract mathematical concepts

Despite extensive research on numerical cognition (for a review see Piazza and Eger 2016), the neural representation of mathematical concepts as a broader semantic class remains poorly understood. A large body of work focused on the representation of concrete numerosities and, using fMRI in humans and electrophysiology in non-human primates, consistently identified the intraparietal sulcus (IPS) as a key hub for basic number sense (for a review see Nieder and Dehaene 2009). This representation is thought to play a foundational role in the human-specific expansion of mathematical concepts, yet how education reorganizes it remains debated (Dehaene et al. 2022; Dehaene, Sablé-Meyer, and Ciccione 2025). Using human fMRI, Piazza et al. (2007) showed that the human intraparietal sulcus carries a notation-independent magnitude code common to numerosities and number symbols supporting an abstract number representation in the parietal cortex (see also Eger et al. 2009). Additional activations in dorsolateral prefrontal cortex (PFC) have also been reported (e.g. Arsalidou and Taylor 2011; Simon et al. 2004), supporting a fronto-parietal network for number processing. A major signature of this network, supported by both behavioral (Moyer and Landauer 1967) and neuroimaging evidence (Dehaene et al. 2003; Göbel, Walsh, and Rushworth 2001; Karami, Castaldi, Eger, and Piazza 2025; Nieder and Dehaene 2009; Pinel et al. 2001), is the distance effect: numbers that are close in magnitude lead to similar behavioral and neural responses. This distance effect is consistent with the notion of a mental number line, an internal spatial map on which numbers are ordered. Moreover, human single-neurons tuned to specific numbers have been observed in the mesial temporal lobe (Kutter et al. 2018, 2023, 2024), and whole-brain imaging has revealed spatially organized topographic maps of preferred numerosity at multiple cortical sites using population receptive field modeling (Cai, Hofstetter, and Dumoulin 2023; Harvey et al. 2013, 2015; Harvey and Dumoulin 2017). Using multivariate representational similarity analyses (RSA; Kriegeskorte 2008), Karami et al. (2025) also identified a network of regions exhibiting numerosity-related distance effects, with a spatial distribution that overlapped the ventral and parietal numerotopic maps previously reported by Harvey and Dumoulin (Harvey and Dumoulin 2017).

Beyond adult neuroimaging studies, developmental studies indicate that numerical magnitude can become automatically accessible at a very early age. Van Rinsveld & Schiltz (2025) used a frequency-tagged EEG paradigm to show that children (ages 5–10) automatically extract numerical magnitude information from Arabic digits. Using non-symbolic numerosities, Gennari et al. (2023) found that sleeping 3-month-old infants form a decodable, supra-modal number representation (∼400 ms after stimulus) that generalizes across auditory and visual inputs. These findings imply that abstract number concepts may be spontaneously encoded well before formal education, before they get connected to symbols for mathematical concepts through education.

What is peculiar to the mathematical domain, however, is that, through education, novel and increasingly abstract concepts continue to get added to this semantic field. Even the representation of exact integers gets transformed when acquiring a formal counting system (Pica et al. 2004). Furthermore, with formal schooling, new concepts of numbers are acquired, including zero, negative numbers, decimals and fractions, whose semantic representations remain much less charted. Just like integers, the processing of fractions elicits distance effects: adults are faster to compare or judge the equivalence of two symbolic fractions when they are further apart in numerical value, even when controlling for the distance between numerators and between denominators (Binzak and Hubbard 2020; Gabriel, Szucs, and Content 2013; Hurst and Cordes 2016; Schneider and Siegler 2010; Toomarian, Meng, and Hubbard 2019; but see also Bhatia et al. 2020; Bonato et al. 2007). FMRI studies further support this observation: human fronto-parietal cortex shows neural habituation to the shared magnitude of equivalent fractions, even in the absence of a task, and this effect is modulated by the numerical distance of the deviant fraction to the habituated one (Jacob and Nieder 2009). During a fraction comparison task, activation in the IPS is similarly modulated by numerical distance (Ischebeck, Schocke, and Delazer 2009). Such a distance effect, observed across modalities and even with non-symbolic fractions (Lewis, Matthews, and Hubbard 2016; Matthews and Chesney 2015), suggests that fractions may be integrated with integers on a single mental number line and within a similar cortical circuit. However, the large reliance on flawed integer-based strategies by both children (Braithwaite, Pyke, and Siegler 2017; Cauté et al. 2026;

Clarke and Roche 2009; Ni and Zhou 2005; Rinne, Ye, and Jordan 2017) and adults (Bonato et al. 2007) suggests that fractions are not easy to understand and integrate with previous integer knowledge (Siegler, Thompson, and Schneider 2011). Additionally, although both symbolic and non-symbolic fractions elicit cerebral activations close to that of integers, these do not fully overlap (DeWolf et al. 2016). All in all, while educated adults seem to encode fractions on a mental number line, the automaticity of access to this representation as well as its relation to that of whole numbers remains uncertain.

Beyond arithmetic, Amalric and Dehaene (2016, 2018, 2019) examined fMRI responses to statements from a wide range of non-mathematical and mathematical domains, ranging from arithmetic to algebra, geometry, topology and analysis. The work was initially performed in professional mathematicians and math students, and recently extended to adolescents and young adults with standard schooling (Moreno et al. 2025). Their results indicate the existence of a highly consistent math-responsive network systematically including intraparietal sulci (IPS), inferior temporal gyri (ITG) and dorsolateral prefrontal cortices (dlPFC), and more activated by math statements in any domain than by control non-math statements of similar length and difficulty (e.g. in history, geography or other general cultural knowledge). Beyond this common network, their results pointed to a single robust categorical dissociation: compared to the rest of mathematics, geometry problems elicit additional neural activation in bilateral posterior IPS and left posterior inferior temporal regions. More specific studies of geometrical concepts were recently performed by Sablé-Meyer et al. (2025), who found that viewing geometric shapes selectively engaged math-responsive regions, particularly in IT and IPS. Ciccione and Dehaene (2026) also demonstrated that processing mathematical graphs (scatterplots) recruits both right anterior IPS and right posterior IT/anterior lateral occipital (aLOC) cortices. These findings suggest that geometry, like arithmetic, may be supported by distributed cortical circuits that overlap with the canonical math-responsive network, but may also exhibit partial specificity for geometric concepts.

To systematically investigate the mental representations of math concepts, Debray and Dehaene (2025) recently collected behavioral familiarity and similarity ratings for the 1000 most frequent words of mathematics. About 43% of the explainable variance in human similarity ratings was captured by vector embeddings in a distributional model of semantics, GloVe (Pennington, Socher, and Manning 2014). Interestingly, those GloVe embeddings exhibited vector arithmetic regularities, such as “three” - “triangle” ≈ “four” - “square”, or “three” - “third” ≈ “four” - “fourth”, as previously shown for other linguistic categories (Mikolov et al. 2013), suggesting that the human mental representation of math concepts may encode cross-category numerical-based correspondences. These findings fit with a recent proposal for the development of mathematical concepts, the hypothesis of a shared language of thought (LoT) for arithmetic and geometry (Dehaene et al. 2025). This model proposes that mathematical development starts with a small number of conceptual primitives, to which new concepts can be added by conceptual combinations of previous ones in a language akin to a programing language similar to Logo (Amalric et al. 2017; Dehaene et al. 2022; Sablé-Meyer et al. 2022, 2025). The concept of “square”, for instance, can be encoded by a simple mental program “repeat 4 times (trace a line and turn by a right angle)”. Within such a framework, the syntactic expression associated with the geometric concept “square” thus includes the primitives “4”, “line” and “right angle”. This view predicts that the neural representations of these items should reflect their syntactic expressions in the LoT. As a consequence, concepts related to the same primitive (e.g. “four”, “fourth” and “square”) should all share a coding axis in cortical representational space.

### Goals of the present study

Here, we used 7 Tesla fMRI and millisecond-resolution MEG to evaluate the hypothesis that the neural representations of math concepts form relational structures of this kind, and to probe where, when and how they are represented in the adult human brain. While the vast majority of previous imaging studies examined math concepts at the categorical level (e.g. arithmetic vs analysis, geometry or topology), we took advantage of high-field fMRI to measure and analyze brain activity to individual concepts, thus opening up the possibility of analyzing them as semantic vectors in a high-dimensional space. We investigated the neural representations and the associated temporal signature of fifteen individual concepts from three distinct mathematical categories (integers, fractions, and geometric shapes). We probed whether basic items drawn from these categories elicit spatially distinct responses in the math-responsive network. This allowed us to ask whether neural embeddings of mathematical concepts reflect both categorical distinctions and cross-category relations based on shared numbers: for instance, whether “three,” “third,” and “triangle” elicit similar neural response patterns due to their common relation to the number 3. Using behavioral similarity ratings, high-field 7 Tesla fMRI and millisecond-resolution MEG, we addressed the following questions:

1. How do participants rate the conceptual similarity between elementary math concepts from different domains?
2. Do distinct regions preferentially encode different mathematical domains (e.g., numbers vs geometric shapes)?
3. Which brain regions encode the conceptual relations between math words, and at what time are they activated? We used RSA to ask in which brain regions (fMRI), and at which point in time (MEG) the similarity between brain activation patterns matches participants’ behavioral semantic similarity ratings. We also probed the presence, in both fMRI and MEG, of hypothesis-driven semantic similarity patterns, explicitly capturing categorical structure (integers, fractions or geometric shapes) and within-category numerical distance, consistent with Shepard & Chipman (1970) notion of second-order isomorphism.
4. Do concepts that belong to distinct domains, yet relate to the same number, such as “three” and “triangle” share partially similar neural activations, as might be predicted by the vector arithmetic hypothesis?

A methodological difficulty is that mathematical words drawn from different math domains inevitably differ in other linguistic properties, including length and frequency. In particular, in our stimulus set, number words were, on average, shorter than geometric shape words (e.g. compare “one”, “two”, “three”, “four” with “pentagon”). To mitigate those problems, we focused on the simplest math words (e.g. “square” rather than “quadrilateral”) and used both univariate multiple regression analyses and multivariate RSA to separate semantic effects from other length- or frequency-based influences.

## Result

### Behavioral measure of conceptual similarity

During fMRI, participants rated the semantic similarity between all pairs of the 15 mathematical concepts chosen. To evaluate behavioral data quality for each participant, we calculated the proportion of ordered item pairs that received the same similarity rating across the two runs (i.e., inter-run consistency). Across participants, mean consistency was 69.8% (σ = 9.7%). Furthermore, when detecting two consecutive identical items, for which participants were asked to respond with a specific “identical” button, accuracy was high overall (μ = 85.7%, σ = 9.3%) and all participants performed above 65%.

We then computed the behavioral representational dissimilarity matrix (RDM) by computing, for each ordered pair of items, its mean dissimilarity rating across participants, on a scale from 1 (unrelated items) to 0 (identical items), with intermediate levels 0.666 and 0.333. As shown on Figure 1E, the resulting behavioral RDM was nearly symmetric, as evaluated by the correlation of its upper and lower triangles (Pearson’s *r*(103) = .98, *p* < .001), indicating that ratings were nearly independent of item order.

**Figure 1.**
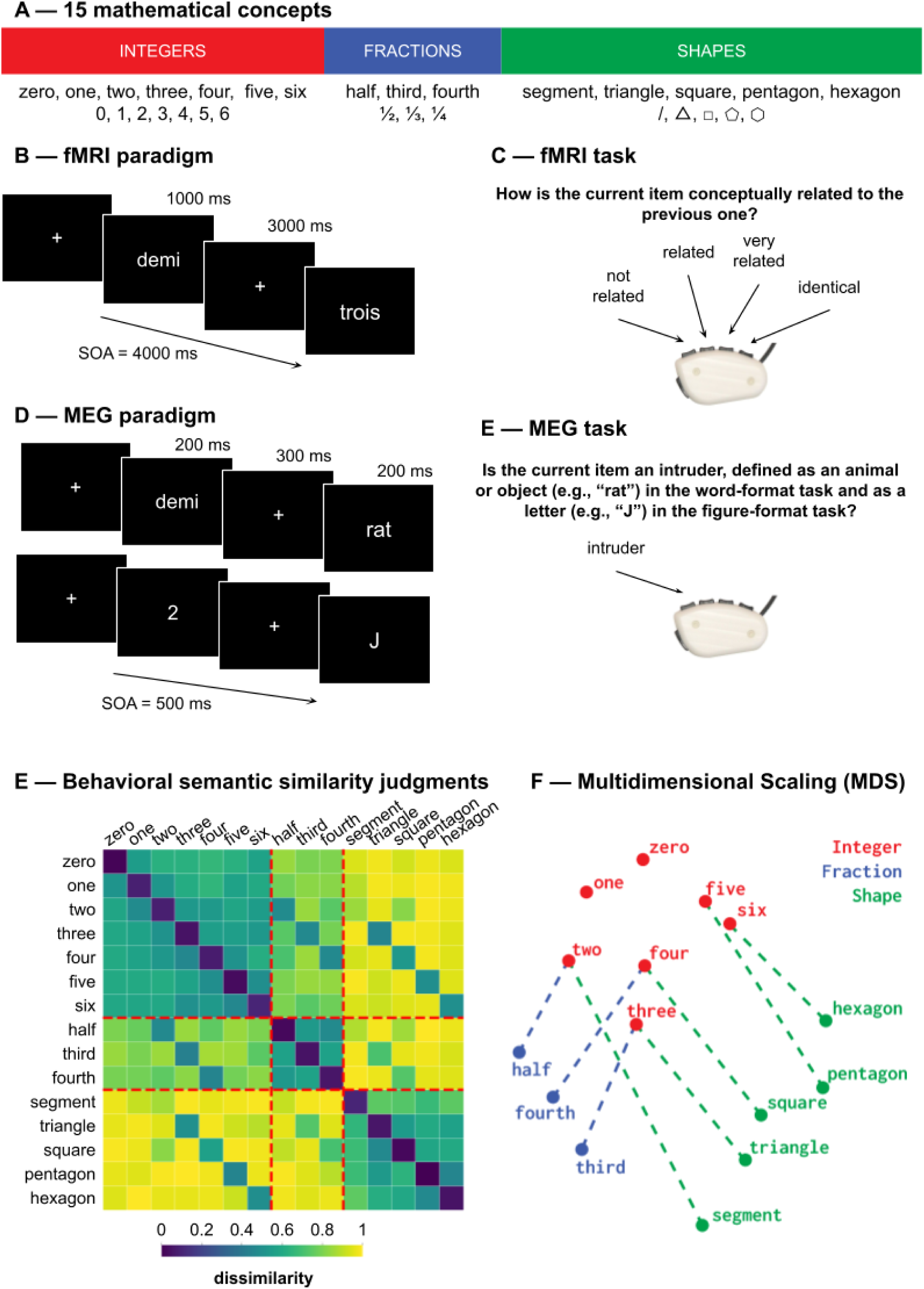
Behavioral semantic dissimilarity ratings. (A) Average dissimilarity matrix acquired during fMRI acquisitions across all participants (N = 18). (B) 2-dimensional metric multidimensional scaling (MDS) of this dissimilarity matrix, indicating systematic groupings of integers, fractions and shapes, as well as vector correspondences between them (dashed lines).

The matrix showed a semantic categorical organization. Mean dissimilarity within integers, fractions or shapes was 0.58 (σ = 0.05), whereas mean dissimilarity across those categories was 0.86 (σ = 0.14). A two-sample t-test confirmed a robust effect of category membership, *t*(208) = −14.71, *p* < .001. Also, items associated with the same number (e.g., “three” and “triangle”, “three” and “third”) were rated less dissimilar than items associated with different numbers. Mean dissimilarity was lower for same-number pairs (μ = 0.55, σ = 0.10) than for different-number pairs (μ = 0.80, σ = 0.17), *t*(208) = −6.11, *p* < .001. This effect can be seen as small cross-category diagonals on Figure 1E (but note how the word “segment” failed to strongly relate to either “two” or “half”). We then aimed to model the behavioral RDM more comprehensively, by fitting the full model reported in the RSA methods section (see also Figure S1). The model explained a substantial proportion of variance, *R²* = .95, *F*(14, 210) = 282.0, *p* < .001. Significant effects emerged for category predictors (integer-integer pairs: β = 0.42, *t*(210) = 16.85, *p* < .001; fraction-fraction pairs: β = 0.39, *t*(210) = 9.73, *p* < .001; integer-fraction or fraction-integer pairs: β = 0.12, *t*(210) = 3.33, p = .001; shape-shape pairs – excluding “segment”: β = 0.43, *t*(210) = 12.42, *p* < .001) and for numerical-based correspondence across categories (integers and fractions associated with the same number: β = 0.35, *t*(210) = 8.51, *p* < .001; integers and shapes associated with the same number: β = 0.44, *t*(210) = 20.86, *p* < .001; fractions and shapes associated with the same number: β = 0.22, *t*(210) = 7.41, *p* < .001). A small numerical distance effect was observed for integers (β = 0.04, *t*(210) = 2.17, *p* = .03), but not for other categories. Finally, we also observed effects of the “segment” category (β = 0.08, *t*(210) = 6.27, *p* = .03).

To obtain a visualization of the underlying semantic space, we applied metric multidimensional scaling (MDS) on the behavioral RDM (Figure 1F). A 2-dimensional representation revealed three separate clusters corresponding to integers, fractions, and shapes, as well as parallel alignments of items associated with the same number (numerical-based correspondence). Consistent with the observed distance effect, integers were arranged according to a “C” curve, similar to what is usually found in the literature (Karami, Castaldi, Eger, and Piazza 2025; Karami, Castaldi, Eger, Hebart, et al. 2025; Nelli et al. 2023).

Together, these behavioral analyses confirm that participants’ similarity ratings capture both categorical structure and associations of items across categories, giving hope that we might find similar effects in the brain.

### Univariate fMRI contrasts between mathematical categories

#### Math versus non-math contrast

As a first sanity check, we confirmed that, using a classical sentence localizer task, we could retrieve the expected parieto-temporo-frontal math-responsive network (Amalric and Dehaene 2016, 2019). Consistent with prior work the contrast math > non-math (more specifically, arithmetic statements + geometric statements > general knowledge + contextual knowledge) was highly similar to those reported by (Amalric and Dehaene 2016) (Figure 2A), with the most significant activation lying in the bilateral intraparietal sulci (IPS) and adjacent inferior parietal regions, bilateral inferior temporal gyri (ITG), and a large swath of bilateral dorsolateral prefrontal cortices. The table of peak coordinates for this contrast is provided as supplementary material S2.

**Figure 2.**
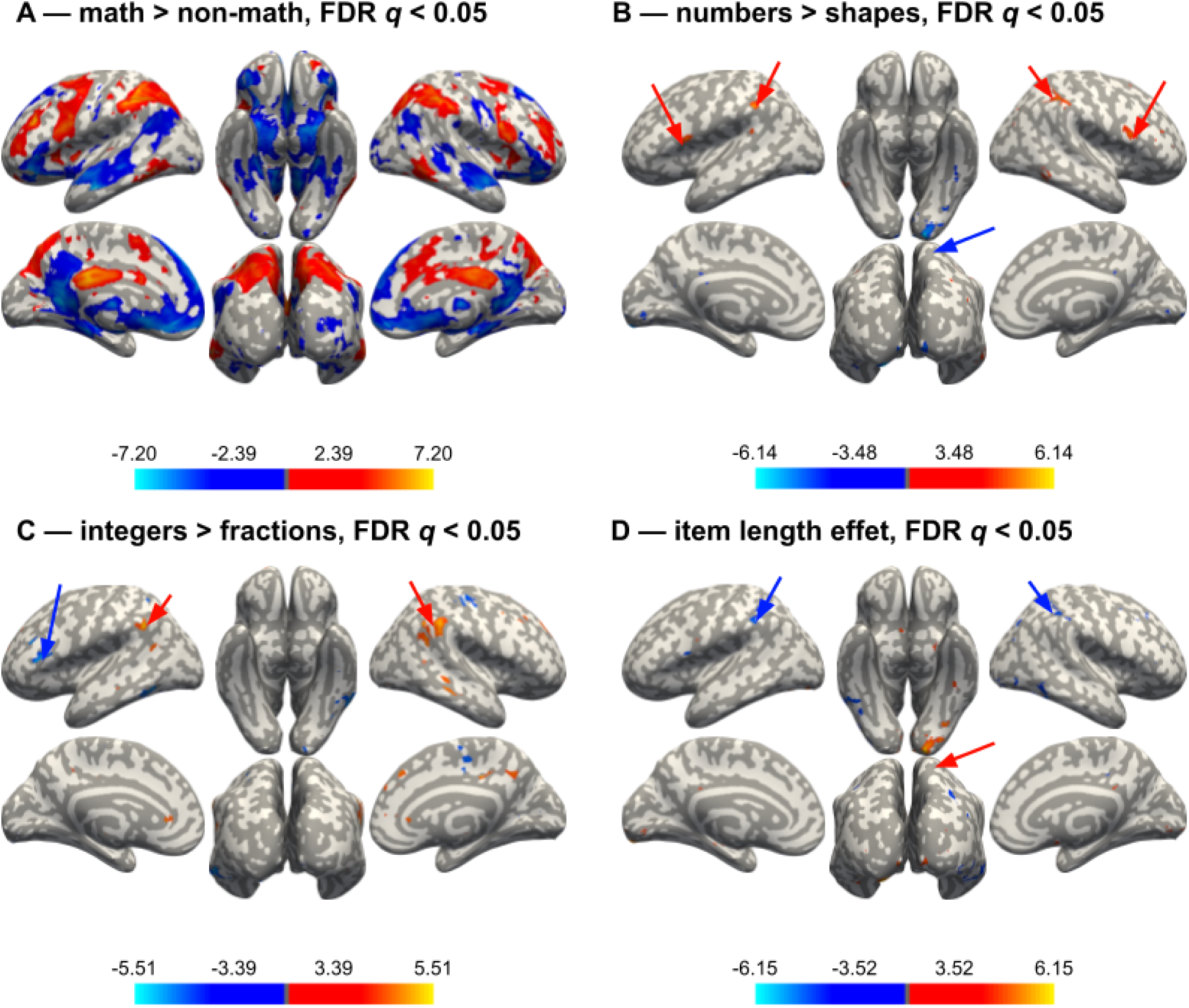
Univariate analyses. (A) Mathematical > non-mathematical statements in the localizer task. Red shows the areas more activated by math sentences, and blue those more activated by non-math sentences. (B-D) Contrasts from a mass-univariate mixed-effect linear model predicting word betas of the first-level GLMs with fixed effects of category (integers, fractions or shapes), word length, and a random intercept by participant. (B) Number words > shape words. Red shows the areas more activated by numbers (integers or fractions) than by shapes, and blue the converse. (C) Integers > fractions. Red = greater activation for integers than for fractions, blue = the converse. (D) Effect of item length (number of letters). Red = greater activation for longer words, blue = greater activation for shorter words. All maps were FDR-corrected for multiple comparisons (q < .05).

#### Numbers versus shapes

We next examined whether specific regions were preferentially recruited by single words referring to numerical versus geometric concepts. In a univariate mixed-effect GLM with category and word length as regressors, the contrast numbers > shapes (i.e. integer + fraction items > shape items) revealed stronger activation in the bilateral posterior supramarginal gyri, bilateral dorsolateral prefrontal cortices, left postcentral gyrus and right inferior frontal gyrus. Conversely, the contrast shapes > numbers showed increased activity in the left LOC, left middle temporal gyrus, right postcentral gyrus, left angular gyrus and left precuneus. The contrast map is shown on Figure 2B and a table of peak coordinates for this contrast is provided as supplementary material S3.

Remember that category and word length are partially confounded: number words contained significantly fewer letters than shape words (Welch’s *t*(6.33) = −4.17, *p* = .005). The mixed-effect model was meant to mitigate this issue, and we verified that the parametric effects of word length was indeed partially disjointed from the number-shape contrast reported above. The length regressor, shown on Figure 2D, shows stronger activation for longer items in bilateral occipital poles, left postcentral gyrus and left anterior cingulate gyrus; and stronger activation for shorter items in bilateral inferior temporal gyrus, bilateral superior LOC and bilateral IPS. Visual inspection confirms that the spatial maps of item length and shapes > numbers were indeed partly different, suggesting that the mixed linear model offered a first step towards disentangling the effects of math category and item length. The table of peak coordinates for this contrast is provided as supplementary material S4.

#### Integers versus fractions

We then tested for potential specialization within the number domain by comparing integers and fractions (Figure 2C). Unlike the previous contrast, integers and fractions did not differ significantly in number of letters (Welch’s *t*(7.80) = −1.13, *p* = .29). The integers > fractions contrast revealed stronger activation for integers in right supramarginal gyrus, left angular gyrus, right cingulate gyrus, bilateral frontal pole, bilateral middle frontal gyrus, right posterior middle temporal gyrus and bilateral precuneus; and stronger activation for fractions in the left ITG, left inferior frontal gyrus, right precentral gyrus, left IPS, left superior temporal gyrus and left cingulate gyrus. The table of peak coordinates for this contrast is provided as supplementary material S5. The integer-preferring regions largely overlap with components of the default mode network (DMN). One possible interpretation is that fractions, being conceptually harder to process than integers, lead to reduced activity in DMN regions. However, since response time (as a proxy for difficulty) was regressed out in the GLMs, these differences cannot be straightforwardly explained by difficulty alone.

#### Double dissociation between numbers and shapes in subject-specific ROIs

To work around the issue of the confound between word length and math categories, as a first approach at the univariate level, we implemented a two-step approach: we first used the localizer task (in which sentences were matched in length) to identified voxels preferring arithmetic or geometry statements for each participant, and then analyzed how these regions also showed a preference for individual math words of the same domain in the main task. Specifically, subject-specific ROIs were defined individually from the independent localizer contrast (arithmetic > geometry and geometry > arithmetic), constrained to those anatomical parcels from the HCP-MMP1.0 atlas (Glasser et al. 2016) that were located along the IPS and ITG (see Methods and supplementary file S6). For each participant, mean beta values were extracted for all items of the main task and tested for an effect of category (integers, fractions, shapes) using a likelihood ratio test between two linear mixed effects model predicting mean beta values: one with fixed effects of category and length and a random intercept by participant, and one with only a fixed effect of length and a random intercept by participant. Only regions showing a significant χ^2^ for model comparison after FDR correction (*q* < .05) and positive overall activation were retained. See Methods section for more details.

After FDR correction, mean betas in 10 regions were better explained by a model taking category into account than by a model only using length as a predictor. These regions are shown on Figure 3 and described in Table 1, in addition, the beta plots in each region are provided as supplementary material S7.

**Figure 3.**
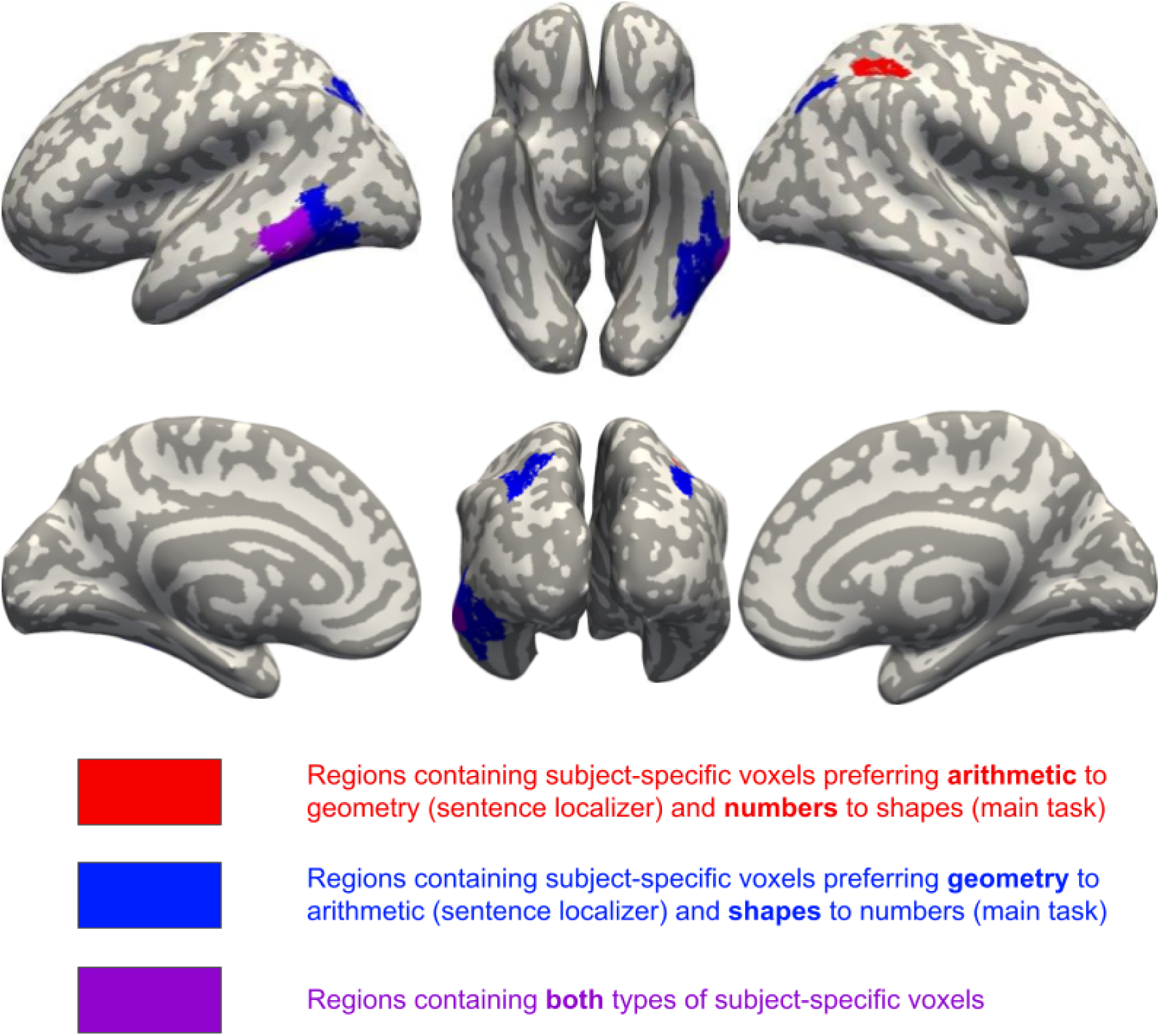
**Double dissociation between number words and shape words in the main task**, within subject-specific voxels defined using the independent sentence localizer (see Table 1 for statistics).

**Table 1.**
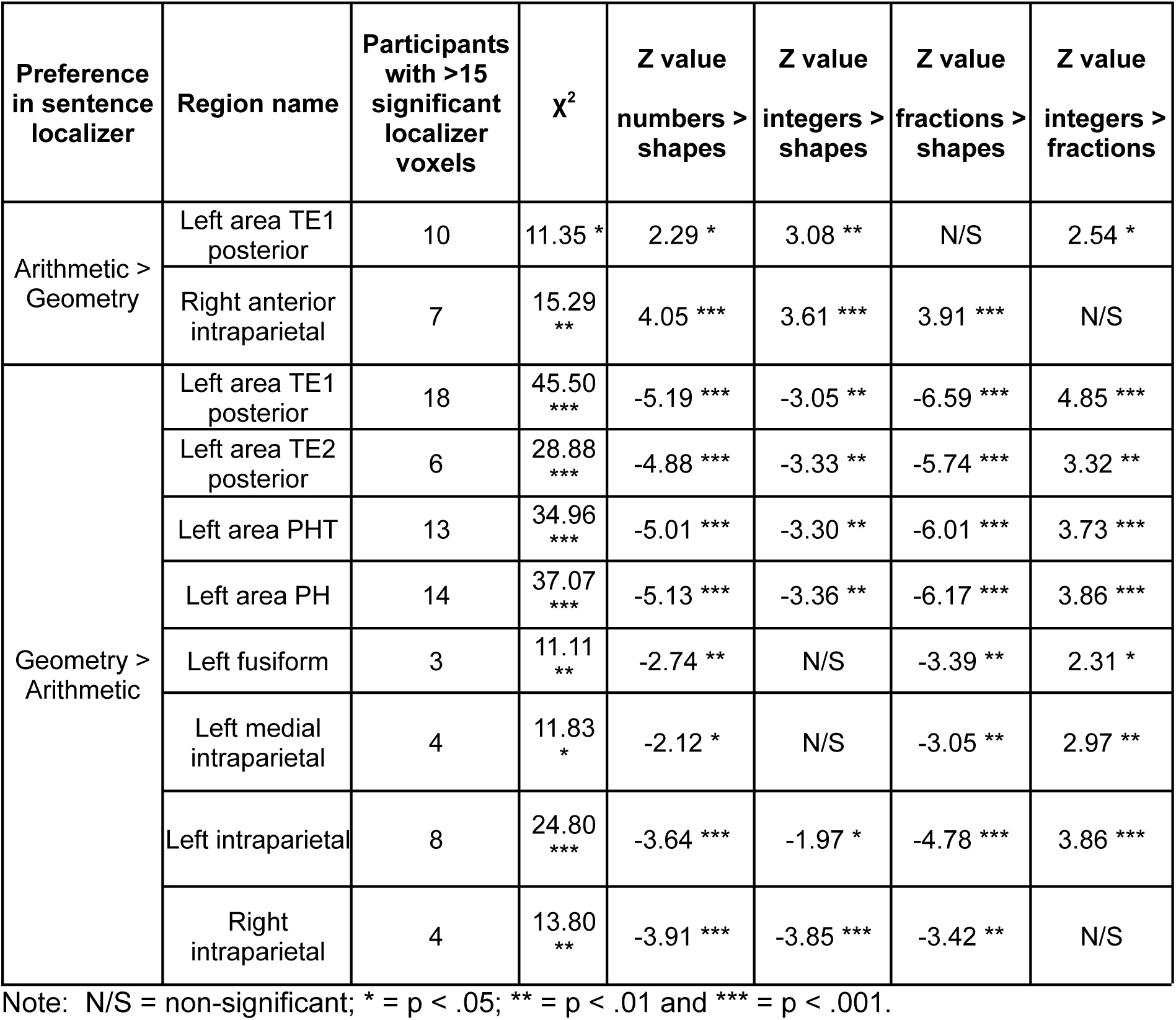
Brain regions from the HCP-MMP1.0 atlas and containing subject-specific voxels showing preferring arithmetic (or geometry) in the sentence localizer and numbers (or shapes) in the main task (see Figure 3 for a visualization of the regions).

In all regions, the direction of preference identified in the localizer was replicated in the main task, with no instances of reversed selectivity. A clear hemispheric asymmetry was observed: shape-preferring regions were bilateral, whereas number-preferring regions were confined to the right hemisphere. Within the right IPS, an anterior-posterior gradient was apparent, with number-related regions located anteriorly and shape-related regions posteriorly, as visible on Figure 3. Interestingly, one region, left area TE1, contained both voxels preferring numbers and voxels preferring shapes. This suggests that this region pulls together neural populations that play different roles in math cognition.

Together, these findings demonstrate that 7T fMRI can resolve adjacent cortical territories selectively engaged in arithmetic and geometric processing. However, it is still possible that, for single words, this dissociation arose due to differences in word length that would overlap with but be unrelated to the preference for arithmetic vs geometry sentences. We therefore turned to a second method, representational similarity analysis, to better separate the effects due to word length and to conceptual similarity.

### fMRI encoding of conceptual similarity

#### Whole-brain searchlight

A first searchlight RSA was used to identify, across the whole brain, regions that responded to conceptual similarity for elementary mathematics, and regions that cared about visual similarity between words. Thus, the regressors included behavioral semantic similarity ratings (see Figure 1A) and the logarithm of the difference in item length. Semantic similarity was a significant predictor of neural RDM in a distributed fronto-parieto-temporal network, shown on Figure 4A, comprising bilateral intraparietal sulci (IPS), the left supramarginal gyrus, and bilateral middle/inferior temporal gyri (MTG/ITG). Additional clusters were found in dorsolateral prefrontal regions, including the left precentral gyrus, the left subcallosal cortex, and the right precentral gyrus. This network, associated with behavioral similarity ratings, closely resembled the math-responsive network identified with the localizer for the math > non-math contrast (see Figure 2A, 21.53% of the voxels activated here were also positively activated in the math > non-math contrast), suggesting that math-responsive regions encode conceptual similarities between math words.

**Figure 4.**
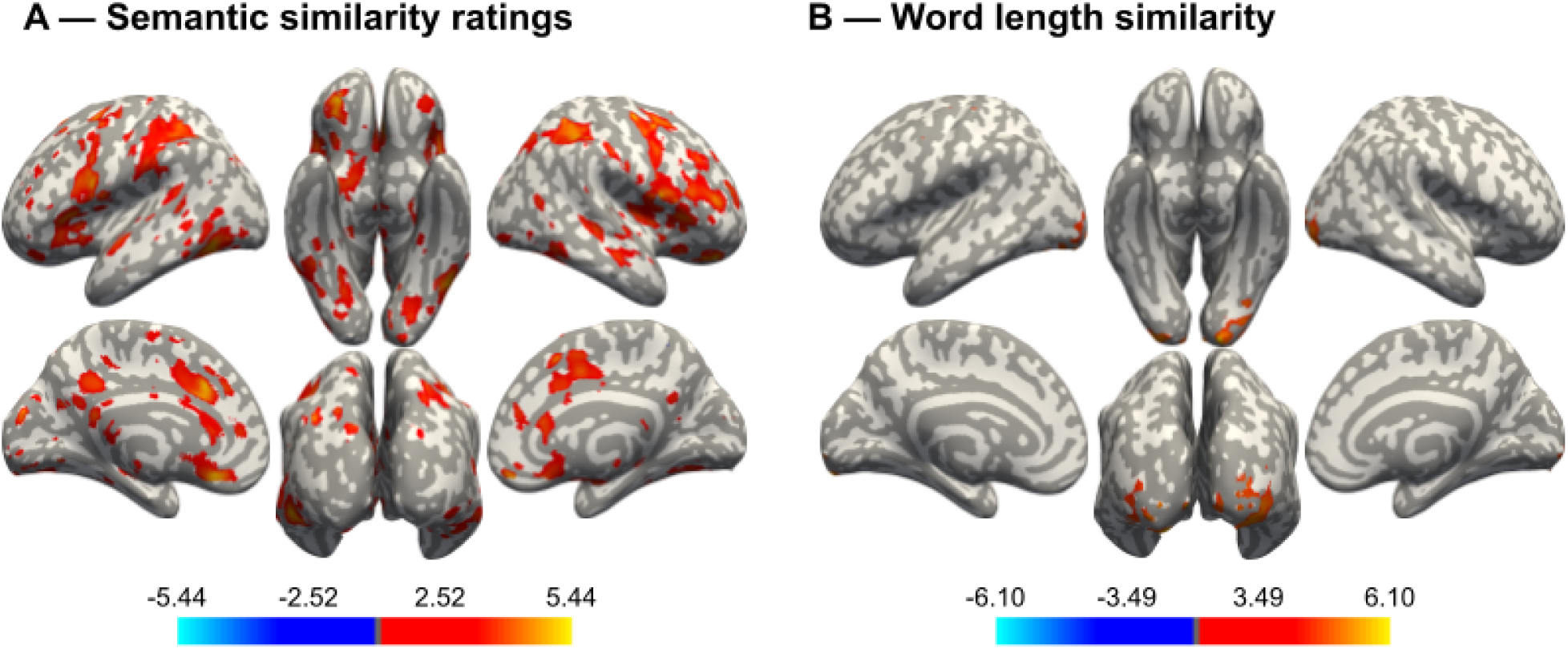
Searchlight RSA maps obtained using the behavioral model. (A) Effect of the semantic similarity ratings predictor and (B) effects of the word length predictor.

The log difference in item length was only associated with activation in bilateral occipital cortex (see Figure 4B), which is consistent with the expected sensitivity of visual regions to low-level visual properties and provide a useful validation of the searchlight approach. Tables giving the coordinates of the significant clusters and their peaks are provided for both the behavioral RDM and the item length predictors in the supplementary material S8 and S9 respectively.

As mentioned in the Methods section, this analysis was also run with an additional predictor for word frequency and yielded very similar results.

While these findings identify the neural basis of participant’s behavioral similarity ratings, the searchlight approach did not provide sufficient statistical power to test finer-grained effects, as the other two RDM models (categorical and full) did not reach FDR-corrected significance at the whole-brain level. We therefore turned to ROI-based RSA to further refine the characterization of this network.

#### ROI-based Representational Similarity Analysis

We focused on six a priori fMRI regions previously implicated in mathematical processing (Amalric and Dehaene 2016; see Methods): bilateral IPS, ITG, and dlPFC. The corresponding average RDMs, extracted from subject-specific voxels isolated from the localizer contrast (math > non-math), are shown on Figure 5.

**Figure 5.**
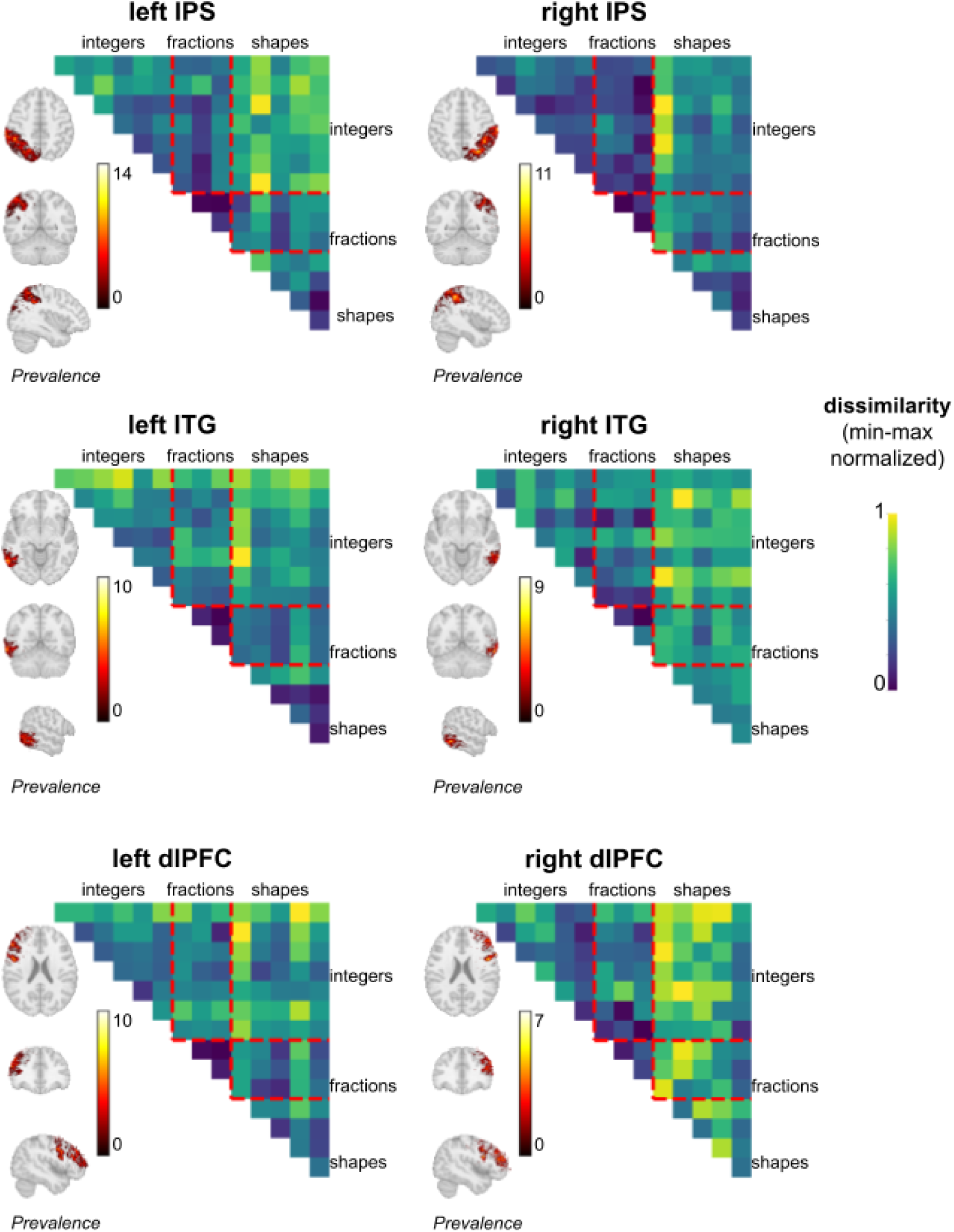
Average representational dissimilarity matrices in 6 functional ROIs: bilateral IPS, ITG and dlPFC. Within each ROIs, we only analyzed subject-specific voxels from the math > non-math sentence localizer. Prevalence indicates, for each voxel, the number of participants with a significant effect in the sentence localizer. For each ROI, three orthogonal slices were chosen to best represent the prevalence maps. Abbreviations: int = integers; frac = fractions.

The category model, which included predictors for length similarity, numbers, and shapes revealed significant similarities in the encoding of numbers in the right IPS (*t*(17) = 3.02, *p* = .008), right ITG (*t*(17) = 2.68, *p* = .02), and right dlPFC, (*t*(17) = 2.35, *p* = .03), and significant similarities in the encoding of shapes (excluding ”segment”, see Methods) in bilateral IPS (left: *t*(17) = 2.51, *p* = .02; right: *t*(17) = 2.65, *p* = .02) and left ITG (*t*(17) = 4.72, *p* < .001).

The finer-grained, full model, which differentiated between integers and fractions and included within- and across-categories predictors, revealed additional effects. We observed significant similarities in the encoding of fractions in the right IPS (*t*(17) = 2.29, *p* = .04) and the right dlPFC (*t*(17) = 2.26, *p* = .04), and significant similarities in the encoding of shapes in the left ITG (*t*(17) = 2.41, *p* = .03) and right ITG (*t*(17) = 2.80, *p* = .01), in full agreement with the univariate results previously reported. We also observed significant similarities in the encoding of matched integers and fractions (e.g., “three” and “third”) in the right IPS (*t*(17) = 2.33, *p* = .03), and in the encoding of matched fractions and shapes (e.g., “third” and “triangle”) in the left ITG (*t*(17) = 2.55, *p* = .02), consistent with the dissociation reported in the univariate analyses. Within categories, we observed a numerical distance effect on the similarity of encodings for shapes in the right ITG (*t*(17) = 2.13, *p* = .048), meaning that shapes whose numbers of edges were numerically close (e.g. triangle and square, i.e. 3 vs 4) were rated as more similar than others (e.g., triangle and pentagon, 3 vs 5).

For a finer-grained analysis, we also examined all math-responsive regions defined by intersecting the HCP-MMP1.0 atlas (Glasser et al. 2016) with each participant’s localizer map for math > non-math sentences. These exploratory analyses only revealed small effects of additional predictors in two isolated regions. We observed in the left frontal eye fields a categorical effect for integer-fraction pairs (*t*(12) = 2.48, *p* = .03) as well as a semantic distance effect between integers and fractions (*t*(12) = 2.41, *p* = .03). We also observed in the right anterior area 32’ (anterior cingulate) a numerical correspondence between integers and shapes (*t*(14) = 2.39, *p* = .03). Given the large number of ROIs tested, these small effects should be interpreted with caution. Here again, adding a predictor for item frequency in the multiple regression did not affect the results.

Overall, fMRI results suggest that math-responsive regions (bilateral IPS, ITG and dlPFC) house partially distinct encodings of arithmetic and geometric concepts.

### Temporal dynamics of access to semantic representations

In addition to fMRI, we ran a second MEG experiment in which the same 15 items were presented at a fast pace (200 ms per stimulus, with a 500-ms SOA) while participants performed an orthogonal intruder detection task. Given that the pace was 5 times faster than in fMRI, the experiment offered enough trials to probe two distinct presentation formats for the same concepts: written words (as in fMRI) and pictorial figures. This design served two purposes. First, by presenting each concept both as a word and as a figure we could test cross-notation generalization: if similarity structure generalizes across formats, this provides stronger evidence that the observed effects reflect abstract conceptual coding rather than modality-specific visual features. Second, by using an orthogonal intruder detection task and rapid serial presentation, we minimized task-related strategies, thereby probing automatic access to the semantic content of abstract mathematical concepts even when their content is task-irrelevant.

#### Time-resolved RSA within format

To characterize the temporal evolution of visual and semantic representations, we performed time-resolved multiple-regression representational similarity analysis (RSA) on the MEG sensor-space data (see Methods). For each time point (−50 to 450 ms relative to stimulus onset), the vectorized neural representational dissimilarity matrix (RDM) was regressed onto visual-feature, lexical, and semantic model RDMs (see Supplementary videos S12 and S13 for dynamic visualizations of the RDMs; see Supplementary videos S15 and S16 for the corresponding MDS representations). Group-level inference was conducted using one-sample *t*-tests against zero, with threshold-free cluster enhancement (TFCE) permutation correction (see Methods).

Visual-feature models produced robust early effects. For word length, significant TFCE-corrected clusters emerged beginning at ∼66 ms and peaked at ∼104 ms. Similarly, the CORnet-S first-layer embedding showed significant effects beginning at ∼73 ms, with a peak at ∼120 ms (Figure 6). These early effects are consistent with a rapid encoding of low-level visual properties and were strongest for stimuli presented in figure format. Notably, visual information remained detectable throughout the epoch, although its magnitude gradually decreased over time.

**Figure 6.**
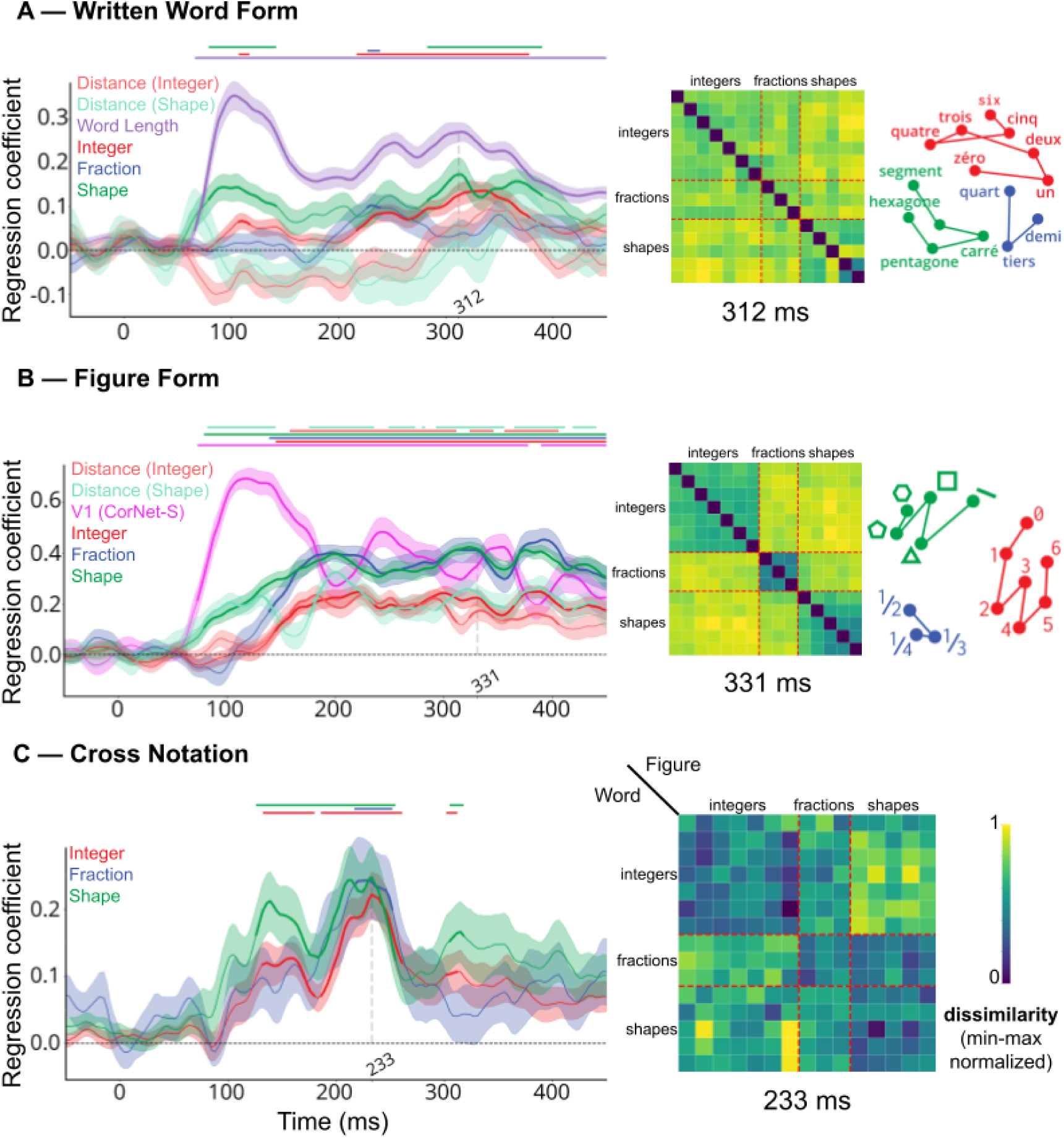
Time-resolved representational similarity results: (A) Time courses of regression coefficients from the representational similarity analysis (RSA) for the written-word-form representation. Predictors: word length, Integer, Fraction, Shape, and distance effects for Integer and Shape. (B) Time courses of regression coefficients from RSA for the figure-form representation. Predictors: embeddings extracted from the V1 layer of Cornet-S (Kubilius et al. 2019), Integer, Fraction, Shape, and distance effects for Integer and Shape. (C) Time courses of regression coefficients from RSA for Integer, Fraction, and Shape in the cross-notation analysis. A sample representational dissimilarity matrix (RDM) at a representative time point and its corresponding multidimensional scaling (MDS) embedding are shown. For a dynamic visualization, see the Supplementary videos S12 and S13 (RDM) and S15 and S16 (MDS). Horizontal coloured dots indicate time points at which effects significantly exceeded zero (p < 0.05, TFCE-corrected). See Figure S4 for the full stimulus set in both written-word and figure formats.

In contrast, the lexical frequency RDM did not yield any significant TFCE-corrected clusters at any time point. Thus, we found no evidence that language-use frequency modulated the representational geometry of the MEG responses.

Most crucially, semantic predictors significantly modulated the neural representational structure. In the figure format, category effects peaked at ∼224 ms (integers), ∼327 ms (shapes), and ∼393 ms (fractions). In the word format, peaks were observed at ∼230 ms (fractions), ∼312 ms (shapes), and ∼337 ms (integers). Regarding fine-grained semantic structure, the within-category distance RDM did not yield significant effects for the written format at any time point, but reliable within-category distance effects were detected in the figure format, specifically for integers and geometric shapes (Figure 6). Together, these results suggest that access to the specific meaning of symbols is more robustly expressed in the MEG signal from visually instantiated numerical and geometric forms than from their written word counterparts, suggesting that digits and shapes promote more automatic semantic access. However, from such within-format analyses alone, any inferences to semantic representations remain fragile because visual features covaried with semantic similarity (see Figure S1). We therefore turned to the more crucial analyses, asking whether and when a cross-format representational similarity effect would emerge.

#### Timing of access to format-invariant conceptual representations

To isolate format-invariant representations and probe whether semantic category information generalized across surface forms, we applied the same regression approach to the cross-notation representational dissimilarity matrices (RDMs), matrices constructed from pairwise correlation between conditions presented in different formats (figure vs. written word) (see Methods; see also Supplementary videos S14 for the temporal evolution of the cross-notation RDM). As in the fMRI analysis, we first fitted a category model including predictors for numbers and shapes. This model revealed robust category-level structure that generalized across formats. Specifically, for numbers, we observed significant intervals (TFCE-corrected) from 132–246 ms and 299–315 ms. For shapes, significant intervals were observed from 126–249 ms and 305–318 ms.

We then fitted a finer-grained, full model that differentiated between integers and fractions. Specifically, our regression model included three category predictors corresponding to integers, fractions, and shapes, entered simultaneously to estimate each category’s unique contribution to cross-format correspondence. The analysis revealed significant category-level structure that generalized across notation: a TFCE-corrected effect emerged in the mid-latency window (approximately 150–250 ms), indicating that the MEG signal carries abstract conceptual information shared between figure and written formats. This result supports the existence of format-invariant conceptual representations during mid-latency stages of processing (Figure 6).

Finally, we examined the contribution of behavioral semantic similarity ratings. Semantic similarity significantly predicted the neural RDM, with significant intervals (TFCE-corrected) at 132–158 ms and 189–252 ms, and a peak effect at 233 ms after stimulus onset.

#### Spatiotemporal mapping of conceptual representations via MEG–fMRI fusion

To relate the temporal dynamics observed in MEG to spatially localized representations identified with fMRI, we performed RSA-based MEG-fMRI fusion. We used a multiple-regression approach in which, for each MEG time point, the group-averaged fMRI RDMs from early visual cortex (V1–V3) and the math network (including parietal, inferior temporal gyrus, and frontal regions) were entered simultaneously as predictors of the MEG RDM. This approach allowed us to dissociate the unique contributions of early visual and higher-level mathematical representations over time.

The analysis revealed a clear temporal dissociation between these two regions. The V1–V3 predictor showed an early and sustained effect, becoming significant at ∼66 ms and remaining significant throughout the entire epoch (66–450 ms), with a peak at ∼101 ms. In contrast, the math network predictor exhibited temporally delayed effects, with a robust and sustained period from ∼217 to 353 ms, peaking at ∼277 ms (Figure 7).

**Figure 7.**
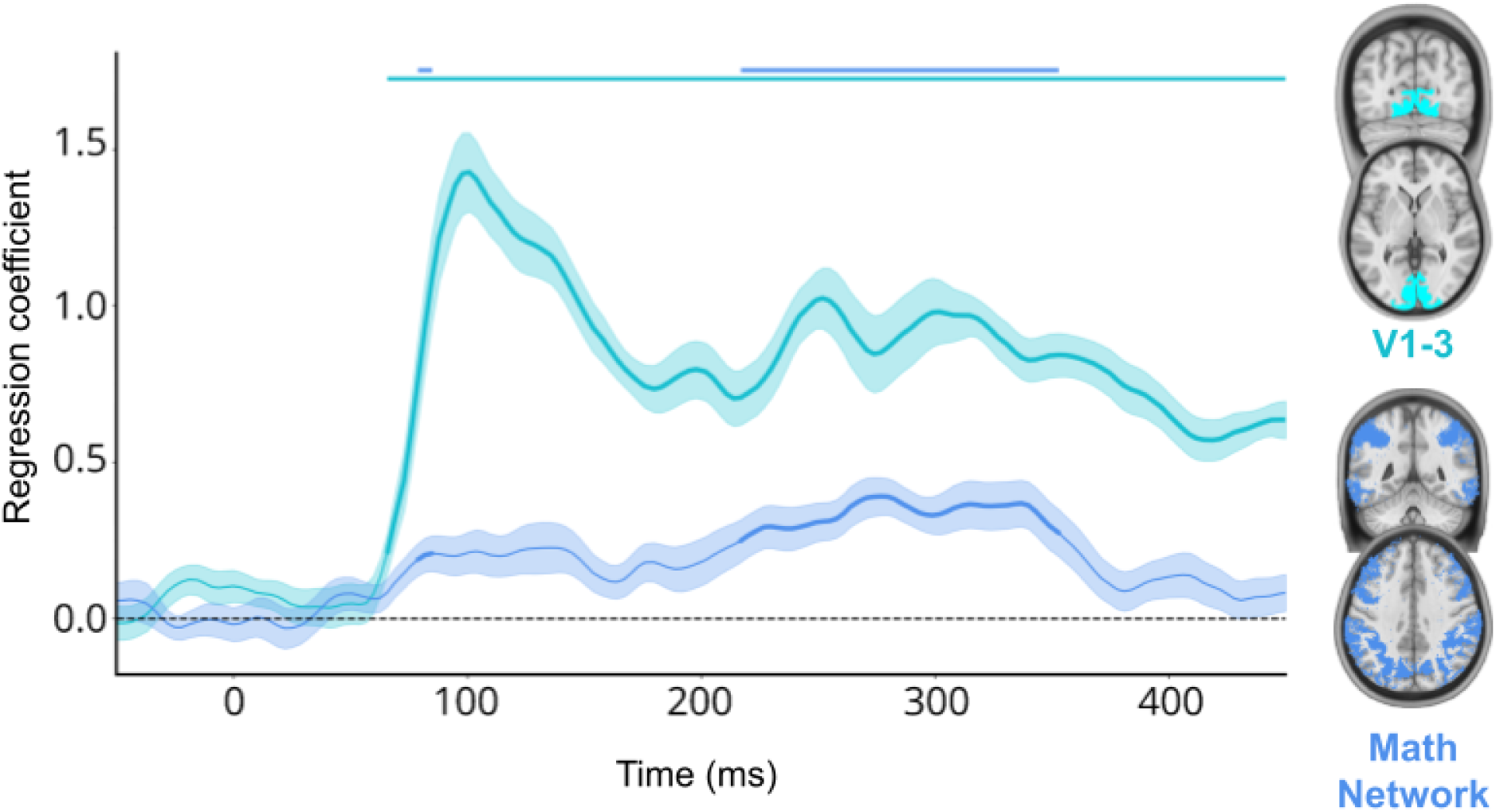
MEG-fMRI fusion results: Time courses of regression coefficients from representational similarity analysis (RSA) for written-word representations across V1–V3 and the math network (including parietal, ITG, and frontal regions) derived from the fMRI study. Horizontal coloured dots indicate time points at which effects significantly exceeded zero (p < 0.05, TFCE-corrected).

To further characterize the contributions of individual components of the math network, we conducted additional regressions in which V1–V3 was entered together with each subregion separately (frontal, parietal, and ITG; see Figure S2). These analyses revealed that frontal, parietal, and ITG regions all showed significant effects in the later time window (frontal: 242–353 ms; parietal: 217–346 ms; ITG: 220–362 ms), with peak responses at ∼337 ms (frontal and ITG) and ∼280 ms (parietal).

Together, these results indicate a temporal progression from early visual representations to higher-level conceptual representations. While V1–V3 dominate early processing, the math network (including parietal, temporal, and frontal regions) contributes at later time points. This pattern supports a posterior-to-anterior cascade in which visual and conceptual representations overlap in time but reach their maximal correspondence at distinct latencies.

## Discussion

The present study investigated how elementary mathematical concepts are organized in the brain, combining behavioral similarity judgments with high-resolution 7 Tesla fMRI, millisecond-resolution MEG, and representational similarity analyses. At the behavioral level, participants’ similarity ratings revealed a clear categorical structure, distinguishing numbers from shapes, arithmetic from geometry, and systematic numerical-based correspondences across categories, such that items sharing the same numerical value (e.g., “three” and “triangle” or “two” and “half”) were judged as more similar to one another. These results indicate that participants spontaneously accessed both categorical distinctions and cross-category numerical relations when comparing elementary mathematical concepts. We then searched for the cortical localization and timing of these effects. Subject-specific ROIs revealed a double dissociation between arithmetic-preferring regions, located primarily in the right anterior intraparietal sulcus, and geometry-preferring regions, encompassing left inferior temporal gyrus and bilateral posterior intraparietal sulcus. This dissociation was further confirmed using ROI-based RSA in math-responsive areas. In addition, whole-brain searchlight RSA identified a distributed fronto-parieto-temporal network in which local multivoxel patterns mirrored the behavioral similarity structure, with numerical-based correspondences across categories effects again located in left ITG and right IPS. MEG added a temporal dimension to these findings by demonstrating categorical and cross-notation semantic effects by about 250 ms, peaking at ∼300 ms in math-related areas.

In terms of localization, these findings closely align with previous work on the neural bases of arithmetic and geometry. Dissociations between these domains are increasingly well documented. A large body of research has linked the intraparietal sulcus to number symbols and arithmetic (Castaldi et al. 2019; Eger et al. 2009; Karami, Castaldi, Eger, and Piazza 2025; Piazza et al. 2004). In parallel, growing evidence indicates that inferior and posterior temporal regions, particularly in the left hemisphere at the border between the ITG and the anterior LOC, respond to geometry and shape-related information (Amalric and Dehaene 2016; Ciccione and Dehaene 2026; Moreno et al. 2025; Sablé-Meyer et al. 2025). These regions have also been implicated in mental imagery (Liu et al. 2025; Liu, Spagna, and Bartolomeo 2022), which is compatible with their present involvement in processing geometric concepts and shapes initially presented as written words. The present results thus fit within an emerging picture in which arithmetic and geometry rely on overlapping but partially distinct neural substrates within the broader math-responsive network, with right IPS playing a dominant role for numerical concepts and left ITG/aLOC for geometric concepts.

In addition to posterior temporal regions, our results point to a specific role of bilateral posterior parietal cortex in geometric processing. These regions have previously been associated with numerical-spatial associations, including links between arithmetic operations and spatial attention (Knops et al. 2009) as well as attentional orientation along the mental number line (Dehaene et al. 2003). Although arithmetic itself is known to recruit spatial representations such as a mental number line (Dehaene, Bossini, and Giraux 1993; Eccher et al. 2025; Mathieu et al. 2016), it is plausible that geometry places even greater demands on these spatial-attentional mechanisms. From this perspective, the stronger involvement of posterior parietal regions in geometry than in arithmetic is coherent with their known functional properties.

Within arithmetic, our results suggest distinct neural processing of the integers “two”, “three” and “four”, compared to the corresponding fractions “(one) half”, “(one) third” and “(one) fourth”. We observed increased activity during fraction processing in several brain regions, matching DeWolf et al.’s (2016) report of partially distinct networks for processing integers and decimals on the one hand, and fractions on the other hand. In particular, DeWolf et al.’s fraction network partially overlaps ours in the left ITG and the left IPS (note that they also reported increased activity for fractions in the right IPS, which we did not observe here). Unlike them, however, we also observed stronger activation during integer processing in several regions, which partially overlapped with the Default Mode Network. Of note, the right MTG, which shows reduced activity for fractions in our data, was previously observed to be less strongly activated by fractions in 5th graders compared to 2nd graders (Park et al. 2025). In any case, the overlap in the pattern of results suggests that distinctions between numerical subdomains are robust across methodological variations and can even be detected with the relatively low spatial resolution of MEG.

An important question is whether the present results were influenced by task design. Our fMRI experiment, inspired by Nelli et al. (2023), required participants to rate the semantic similarity between consecutive items. We opted for this task because it ensures an active engagement and compliance, in contrast to paradigms relying on passive presentation (Mason and Just 2016; Mason et al. 2021). Semantic similarity judgments probably amplified the depth of semantic processing relative to surface-level categorization or number detection (as indicated by the behavioral results on Figure 1E; see also Shepard, Kilpatric, and Cunningham 1975). Furthermore, performing similarity judgments serially requires constantly holding the current item in working memory until the next one appears and can be compared, thus enforcing an active representation of the current target for the entire duration of the inter-item interval and likely amplifying the corresponding neural response. However, this design may also have led, transiently at least, to an overlap of the activations evoked by the previous and current items. Such overlap could blur neural representations and reduce the sensitivity to fine-grained effects, such as numerical distance or correspondence effects. Although all possible item transitions were presented to limit systematic biases, the potential persistence of previous item representations remains a concern. Future fMRI studies may benefit from task designs that more strongly decouple successive stimuli to minimize carryover effects. Note, however, that this potential problem was minimized in the MEG experiment, where participants performed a simple intruder detection task which required minimal processing of the math targets – and yet we found clear evidence for semantic access to categorical information (for both words and figures) and even to fine-grained semantic information (a semantic distance effect for digits). It is likely that the higher temporal resolution of MEG allowed to isolate a brief period of semantic activation which, in BOLD fMRI, would likely be diluted in time and therefore harder to detect. In brief, our design attempted to select an optimal task for each brain-imaging modality.

Several other factors may have contributed to the discrepancy between the richness of the observed behavioral similarity matrix and the small cross-category effects found in fMRI data. The spatial resolution of fMRI may be insufficient to detect fine-grained numerical coding, particularly within integers. It is also possible that neural representations supporting numerical distance are highly distributed and thus more readily expressed at the behavioral level than within localized ROI-based analyses, especially when participants are instructed to consider concepts as mathematical entities rather than just quantities to be compared. Indeed, during the same fMRI session, in distinct runs, the participants also completed a numerical comparison task with symbolic digits and fractions instead of written words (results reported in Valerio et al. 2026). During that task, the activations in math-responsive regions exhibited a clear organization along a mental number line (shared between integers and fractions), with robust distance effects and topographic voxel tuning to specific numerical values reminiscent of the numerosity maps observed by Harvey & Dumoulin (2017). Thus, the choice of task may warp attention and affect whether or not fine-grained semantic effects emerge, as previously reported (e.g. Castaldi et al. 2019; Piazza et al. 2004). Since the present paradigm emphasized conceptual comparison along any semantic dimension, rather than focused attention on magnitude processing, it may have reduced the salience of numerical magnitude codes in neural signals.

The format of the stimuli may also have influenced the results. In fMRI, we deliberately used written word stimuli, while most previous studies in numerical or shape cognition used symbolic digits or non-symbolic perceptual stimuli (Göbel et al. 2001; Kutter et al. 2018; Piazza et al. 2004; Sablé-Meyer et al. 2025). Although there is some evidence in the literature that symbolic and non-symbolic numbers converge onto shared neural substrate (Cai et al. 2023; Eger et al. 2003; Naccache 2001; Piazza et al. 2007), consistent with the triple-code model of number coding (Dehaene 1992; Dehaene and Cohen 1995), it remains possible that non-symbolic or pictorial stimuli would have evoked stronger or more distinct representations. Thus, to complement the fMRI similarity-judgment paradigm and to more directly probe automatic and format-invariant conceptual codes, we ran a complementary MEG experiment with the same 15 items presented much more rapidly. The speed afforded by MEG allowed us to acquire data from more trials, which we used to present the same concepts in two surface formats (written words and figures). This allowed us to test cross-notation generalization: if representational structure is truly conceptual (not merely visual or lexical), similarity relations should generalize across formats. The MEG data provided complementary evidence to the fMRI results. Time-resolved RSA revealed format-invariant category structure in a mid-latency window (peaking around 230 ms, figure 6C), indicating that abstract category information is automatically available in the MEG signal and generalizes across words and figures. Moreover, within-category numerical distance effects were absent for the written-word format, but were reliably observed starting around 150 ms when stimuli were shown in the figure format for both integers and geometric shapes (Figure 6B), suggesting that this format did promote a greater automaticity of access to meaning. Importantly, with non-symbolic displays of numerosity, evidence from time-resolved electrophysiology indicates that a distance effect can occur very early on the order of ∼100 ms (Karami, Castaldi, Eger, Hebart, et al. 2025; Park et al. 2015). For example Karami et al. (2025), using MEG and explicitly controlling for non-numeric visual confounds, found numerosity distance effects emerging very early (reported around ∼115 ms) independently of other non-numeric stimulus features. The present results suggest that digits (at least the frequent ones used here) elicit a distance effect for integers only slightly later in time.

Our MEG results are also consistent with prior time-resolved work on cross-format semantic coding outside the math domain. Dirani & Pylkkänen (2023) reported that category information (tools vs. animals) becomes decodable across picture naming and word reading at ∼150 ms and persists to ∼450 ms, indicating rapid emergence of modality-independent semantic structure. Bezsudnova et al. (2024) similarly found that a classifier trained on images could decode word stimuli (and vice versa) between ∼150–430 ms. Notably, Dirani & Pylkkänen (2024) also showed that these shared conceptual representations correlate with both semantic and visual feature models. Giari et al. (2020) likewise suggested that picture-induced semantic content can emerge as early as ∼105 ms, whereas word-induced conceptual signals appear by ∼230–335 ms. Together, these studies support the idea that pictorial stimuli access semantic content earlier and that mid-latency (≈150–300 ms) format-invariant codes are a common signature of semantic access, a pattern that matches our observation of cross-notation category structure around 150–250 ms and of stronger within-category distance effects for figure presentations.

The absence of a frequency effect may relate to the high degree of item repetition in the present experiment. Recent behavioral work by Corps and Meyer (2025) demonstrated that repeated production of the same reduced set of words can eliminate the word-frequency effect, with effects persisting across sessions. Although our task did not require overt production, repeated exposure to the same 15 written stimuli may have reduced frequency-based differences in lexical accessibility, potentially attenuating frequency-related representational structure in the MEG signal. We note, however, that this interpretation remains speculative and would require direct manipulation of repetition to confirm.

An additional question raised by the MEG–fMRI fusion results concerns the near-simultaneous peak (∼277 ms) observed across the math network, including parietal, inferior temporal (ITG), and frontal regions. Rather than reflecting independent, sequential processing in these areas, this temporal convergence may indicate a late, large-scale integration and information broadcasting process. Global Workspace Theory predicts that once a stimulus is sufficiently processed (around 200–300 ms) it triggers a sudden, widespread activation (“ignition”) in a fronto-parietal network (Dehaene, Changeux, and Naccache 2011; Sergent and Dehaene 2004). The ∼277 ms peak shared by all these regions could reflect such a broadcasting of conceptual category information. This interpretation is consistent with prior EEG/MEG oddball studies showing simultaneous parietal–frontal sources (REFS), and with GWT models that place the “ignition” of conscious content right around the 300 ms latency (Cohen et al. 2024; Dehaene et al. 2011; Sergent and Dehaene 2004). However, this is not the only interpretation. If multiple regions have highly similar representational dissimilarity matrices (RDMs), their fusion signal will be inherently ambiguous (Cichy and Oliva 2020). In our data, the parietal, ITG, and frontal fMRI RDMs were very highly correlated (see Figure S2), so the common ∼277 ms peak may simply reflect their overlapping coding rather than the activity of a single region. To resolve this ambiguity, future studies could use richer stimulus sets that decorrelate the RDMs across ROIs, as well as more precise intracranial recording techniques.

The presence of numerical correspondence effects in behavior, and to a lesser extent in neural data, lends support to the proposal of a shared language of thought linking arithmetic and geometry (Dehaene et al. 2025). However, the neural signatures of these correspondences were weaker and less systematic than their behavioral counterparts and must therefore be interpreted in the context of the task. Given that robust numerical distance effects were observed in the same participants under conditions explicitly requiring magnitude processing (Valerio et al. 2026), the extent to which a shared LoT is recruited may depend strongly on the cognitive operations elicited by the task. Our results therefore point toward a representational architecture that can express LoT-like correspondences but does so selectively depending on task demands.

More broadly, this work illustrates the possibility of measuring neural responses to individual words and using representational similarity analysis to separate conceptual structure from confounding factors such as word length. Recent intracranial work has shown that individual human neurons can encode stimuli along interpretable abstract dimensions (Courellis et al. 2024; Franch et al. 2025; Karkowski et al. 2025), suggesting that conceptual spaces may be vector-like at the neuronal level. With fMRI, whose spatial resolution and signal-to-noise ratio are coarser, multivariate methods provide an appropriate window onto representational geometry. The present work shows that RSA uncover a semantic similarity structure from non-invasive measurements of cortical activation patterns. Previous fMRI attempts at eliciting vector arithmetic (e.g. Ferrante et al. 2025) relied on decoding and therefore could not clearly distinguish compositionality inherent in neural codes from compositionality introduced by the decoder. RSA avoids this ambiguity by analyzing representational structure directly in the measured activation patterns.

Despite this methodological advantage, our goal of demonstrating robust vector-like relationships across mathematical domains was only fully achieved at the behavioral level. Participants’ similarity ratings clearly reflected numerical-based correspondences across categories, as well as a small distance effect within integers. In contrast, RSA revealed only limited numerical correspondence effects, each confined to a single ROI, and no distance effect within integers. Distance effects were observed within fractions and shapes in the right inferior temporal gyrus, but these findings should be interpreted cautiously given the small number of fractions tested. Importantly, in the MEG data we observed a reliable numerical distance effect for integers and a distance effect for geometric shapes specifically when stimuli were presented in the figure format, indicating that fine-grained within-category magnitude relations may be more robustly expressed for perceptual depictions than for written words.

However, our results should not be interpreted as a fundamental failure of vector-based semantics. The idea that concepts are organized in vector-like spaces has gained considerable support (Piantadosi et al. 2024). For example, Wu et al. (2022) showed that adding and subtracting fMRI activation patterns for related words could recover analogical relations. Similarly, intracranial studies have reported converging evidence at the neural level. Franch et al. found that hippocampal neurons fire in patterns reflecting contextual word meaning in a way that mirrors distances in word-embedding space, and Zhu et al. (2026) observed “parallelogram” geometry in single-neuron populations across multiple semantic analogies, meaning that for certain analogies, the activity differences between concept pairs align to form a parallelogram in representational space. The fact that our combined fMRI/MEG study of mathematical concepts (e.g., “three,” “triangle,” “four,” “square”) revealed no evidence for such vector arithmetic may simply reflect the difficulty of accessing such fine-scale information on neural vectors. In neural-network terms, these analogies might lie in finer dimensions. Deep-network theory (Saxe, McClelland, and Ganguli 2019) shows that networks learn representations gradually: early in training they capture only the the primary axes of variation of the data, and only later integrate the weaker ones. Thus, the broadest conceptual distinctions emerge first, and finer distinctions (such as those needed to form precise analogy vectors) appear later. Consistent with this theory, our data showed basic categorical distinctions in all imaging modalities, but only small evidence of finer-grained distinction such as the numerical distance effect. If experience with math concepts plays a role in the strength with which semantic distinctions are encoded in neural dimensions, future work might test expert mathematicians to see if their neural representations exhibit more robust evidence of semantic structures.

Another factor may be the sheer dimensionality of conceptual codes. Recent work finds that cortical sensory representations are “scale-free” and extremely high-dimensional: neural population codes spread information across thousands of latent dimensions (Gauthaman, Ménard, and Bonner 2025). Conventional RSA and low-dimensional analyses, however, are sensitive only to the few largest dimensions and miss this “high-dimensional” structure (Gauthaman et al. 2025). If mathematical concepts likewise occupy a vast latent space, then our RSA might have captured only the tip of the iceberg of the true geometry. In that case, vector-arithmetic relations could exist in lower-variance dimensions beyond our current reach. Future studies using richer recording techniques or spectrum-based methods could reveal whether a fuller, high-dimensional conceptual space supports the predictor vector-semantic structures.

Finally, several limitations and directions for future work should be noted. The regions identified here need not exclusively encode conceptual representations; they may also contribute to the cognitive operations involved in mentally manipulating mathematical concepts, for instance the posterior parietal activation to geometric shapes may reflect attentional shifts, at or near area (Simon et al. 2004). Further work will be required to isolate conceptual representations more precisely, for instance by using cross-modal convergence designs in which the same concepts are presented as words and as pictures, as in Popham et al. (2021). In addition, the present study focused on a small set of fifteen basic concepts. Future work could expand this stimulus set to cover mathematics more broadly, building on our recent database of 1000 mathematical concepts (Debray and Dehaene 2025). Leveraging the observed arithmetic–geometry dissociation, it may become possible to construct large-scale neural embeddings of mathematical knowledge, analogous to linguistic word embeddings such as GloVe (Pennington et al. 2014). Comparing such neural embeddings to computational models could provide a powerful framework for understanding how the human brain represents the abstract concepts of mathematics.

## Materials and Methods

### fMRI Experiment

#### Tasks

Pre-fMRI questionnaire

Before the fMRI acquisition, participants were handed a booklet containing the list of items they would see during the task. For each item, participants were asked to list as many features as they could think of (e.g. “two is an even number”). Participants were instructed to spend roughly one minute per item. This questionnaire (inspired by Mason and Just 2016; Mason et al. 2021) aimed at forcing participants to think about the mathematical properties of the concepts they would see during the task, inducing them to generate richer mental representations for these concepts.

Language and math localizer

In order to identify the math-responsive network in each participant and to define math related functional regions-of-interest (ROIs), fMRI began with a localizer task in which participants judged the truth value of statements about mathematical and non-mathematical topics. Participants were presented with 20 statements (10 true and 10 false) from each of these four categories: general knowledge (e.g. “The human body is made up of billions of cells.”), contextual knowledge (e.g. “In Beijing, pollution is a major issue.”), arithmetic statements (e.g. “The square root of thirty-six is greater than five.”) and geometric statements (e.g. “The diameter of a circle is equal to twice its radius.”). The full list of stimuli is given in supplementary file S10. The statements were displayed word by word on the screen, with each word being displayed for 350ms as white text on a black background (mean statement duration 3277ms). At the end of a statement, a fixation cross was displayed until the beginning of the next statement. The stimulus-onset-asynchrony (SOA) between statements was jittered according to the formula SOA = 3500ms + 300×J where J follows a Poisson distribution with parameter 15 (so that the expected value of the SOA as a random variable is 8s), with the constraint that mean SOA was actually exactly 8s. Before the next statement, the fixation cross turned red for 350ms to warn participants. Half of the participants were instructed to press the leftmost button when they thought the statement was true and the rightmost button when they thought it was false, and the other half received the opposite instructions. Participants did not receive any feedback during the task.

The run started with a 2s display of a fixation cross to focus participants’ attention and ended with another display of a fixation cross, whose duration was set to fix the total length of the run to 11mn, allowing the BOLD signal to return to baseline. See Figure S3 (supplementary material S11) for a visual summary of this task’s design.

Similarity rating task

In the main fMRI task, the stimuli were fifteen math words: six integers (0 to 5; “zéro”, “un”, “deux”, “trois”, “quatre”, “cinq” in French), three fractions (the first three unit fractions, expressed as a single word; “demi”, “tiers”, “quart”), and five geometric shapes (polygons from two to six vertices; “segment”, “triangle, “carré”, “pentagone”, “hexagone”). Note that, while the English words “third” and “fourth” are ambiguous (the same words are used for ordinals and fractions), no such ambiguity affects the corresponding French words “tiers” and “quart”, which differ from the ordinals “troisième” and “quatrième”.

These stimuli were presented serially to the participants, who were instructed to rate the conceptual similarity of each concept with the preceding one (see Figure 6). This task was directly inspired by Nelli et al. (2023) who showed that it allows to measure the activation to each target and reconstruct their mental organization. Using a 4-button response pad held with both hands, participants were instructed to press the leftmost button (with their left middle finger) when they thought the current items was “unrelated” to the previous one, the second button from the left (with their left forefinger) for “somewhat related”, the second button from the right (with their right forefinger) for “very related”, and the rightmost button (with their right middle finger) when they were “identical” (see Figure 6B).

Participants took two runs of 16 min each. During each run, they saw every possible transition between items (including from an item to itself), resulting in a total of 15×15 = 225 different transitions. Each run was divided into five blocks of 46 items (i.e. 45 similarity ratings) each, separated by a rest period of 10s. The items were presented in white on a black background, displayed for 1000ms, followed by a fixation cross (SOA = 4s, jittered between 2s and 4s). A black screen with a white fixation cross was presented for 3s at the beginning and for 10s at the end of each run.

Participants were informed that their response was recorded when the fixation cross turned green. The fixation cross turned red 500 ms before the onset of the next stimulus to warn participants of the upcoming trial. To increase participants’ motivation, a screen with their percentage of correct answers appeared every 46 stimuli. Before the session, we emphasized that the feedback was only provided for information purposes and that there were no right or wrong answers. The feedback was created by asking another group of participants to rate the relation between concepts on a scale from 0 (unrelated concepts) to 7 (identical concepts). The average ratings from this control group were then divided into quartiles, corresponding to a 0-3 scale. In fMRI, the feedback shown to each participant corresponded to the percentage of trials in which their responses matched the quartile classification derived from the control group. This feedback was displayed for 3s and was followed by a resting period of 7s, during which a white fixation cross was displayed on screen. This fixation cross turned red 1s before the task resumed, and the last item presented before the feedback was once again presented after the resting period, to remind participants of it.

#### Participants

Twenty-two healthy adults (gender: 13 females, 9 males; age: range = 19-39 y.o., μ = 26.5 y.o., σ = 5.6 y.o.) participated in the experiment. Out of these initial participants, four were excluded (respectively: panic attack; temporal-lobe cyst; low accuracy [37%] in the main task; lack of task engagement and motor and visual activations). All results reported in this work were obtained from the remaining eighteen participants (gender: 9 females, 9 males; age: range = 20-39 y.o., μ = 27.1 y.o., σ = 5.9 y.o.). All were native French speakers who had studied math until at least high-school graduation (baccalauréat). They were all right-handed, had normal or corrected-to normal vision and had no history of neurological or psychiatric disorders. Participants provided written informed consent for the fMRI study and received monetary compensation. The study was approved by the ethics committee (Comité de Protection des Personnes Sud-Ouest et Outre-mer III; references: CEA 100 055, ID RCB 2020-A01713-36) and was conducted in accordance with the Declaration of Helsinki.

#### MRI parameters

MRI data were acquired using a 7T MAGNETOM whole-body MR scanner (Siemens Healthineers, Erlangen, Germany) with a 32-channel head coil (Nova Medical, Wilmington, USA) at the NeuroSpin Center. Structural MRI data were collected using T1-weighted rapid gradient echo (MPRAGE) sequence (repetition time (TR) = 5000 ms, echo time (TE) = 2.51 ms, resolution = 0.65 mm isotropic, field of view (FoV) = 208⨯208 mm2, flip angle = 5/3, bandwidth (BW) = 250 Hz/px, echo spacing = 7 ms). Functional MRI (fMRI) data were acquired using a T2*-weighted gradient echo planar imaging (EPI) sequence (TR = 2000 ms, TE = 21 ms, voxel size = 1.2 mm isotropic, multiband acceleration factor = 2, FoV = 192⨯192 mm2, flip angle=75, BW=1488 Hz/px, partial Fourier = 6/8, echo spacing = 0.78 ms, number of slices = 70 (no gap)). To correct for EPI distortion, a five-volume functional run with the same parameters except for the opposite phase encoding direction (posterior to anterior) was acquired immediately before each task run. Manual interactive shimming of the B0 field was performed for all participants. The system voltage was set at 250 V for all sessions, and the fat suppression was decreased to ensure that the specific absorption rate (SAR) did not surpass 62% across all functional runs.

#### MRI data preprocessing

Preprocessing was performed using fMRIPrep 22.0.2 (Esteban et al. 2019; Markiewicz et al. 2026) (RRID:SCR_016216), which is based on Nipype 1.8.5 (Esteban et al. 2025; Gorgolewski et al. 2011) (RRID:SCR_002502).

Preprocessing of B0 inhomogeneity mappings

For each participant, a B0-nonuniformity map (or fieldmap) was estimated based on two (or more) echo-planar imaging (EPI) references with topup (Andersson, Skare, and Ashburner 2003) (FSL 6.0.5.1:57b01774).

Anatomical data preprocessing

T1-weighted (T1w) images were corrected for intensity non-uniformity (INU) with N4BiasFieldCorrection (Tustison et al. 2010), distributed with ANTs 2.3.3 (Avants et al. 2008) (RRID:SCR_004757), and used as T1w-references throughout the workflow for each participant. The T1w-references were then skull-stripped with a Nipype implementation of the antsBrainExtraction.sh workflow (from ANTs), using OASIS30ANTs as target template. Brain tissue segmentations of cerebrospinal fluid (CSF), white-matter (WM) and gray-matter (GM) were performed on the brain-extracted T1w using fast (FSL 6.0.5.1:57b01774, RRID:SCR_002823) (Zhang, Brady, and Smith 2001). Brain surfaces were reconstructed using recon-all (FreeSurfer 7.2.0, RRID:SCR_001847) (Dale, Fischl, and Sereno 1999), and the brain masks estimated previously were refined with a custom variation of the method to reconcile ANTs-derived and FreeSurfer-derived segmentations of the cortical gray-matter of Mindboggle (RRID:SCR_002438) (Klein et al. 2017). Volume-based spatial normalization to one standard space (MNI152NLin2009cAsym) was performed through nonlinear registration with antsRegistration (ANTs 2.3.3), using brain-extracted versions of both T1w references and the T1w templates. The following template was selected for spatial normalization: ICBM 152 Nonlinear Asymmetrical template version 2009c (Fonov et al. 2009) (RRID:SCR_008796; TemplateFlow ID: MNI152NLin2009cAsym).

Functional data preprocessing

For each of the five BOLD runs found per subject (across all tasks and sessions), the following preprocessing was performed. First, a reference volume and its skull-stripped version were defined using a single-band reference (SBRef). Head-motion parameters with respect to the BOLD reference (transformation matrices, and six corresponding rotation and translation parameters) are estimated before any spatiotemporal filtering using mcflirt (FSL 6.0.5.1:57b01774) (Jenkinson et al. 2002). The estimated fieldmap was then aligned with rigid-registration to the target EPI (echo-planar imaging) reference run. The field coefficients were mapped on to the reference EPI using the transform. BOLD runs were slice-time corrected to 0.975 s (0.5 of slice acquisition range 0 s-1.95 s) using 3dTshift from AFNI (Cox and Hyde 1997) (RRID:SCR_005927). The BOLD reference was then co-registered to the T1w reference using bbregister (FreeSurfer) which implements boundary-based registration (Greve and Fischl 2009). Co-registration was configured with nine degrees of freedom to account for distortions remaining in the BOLD reference. First, a reference volume and its skull-stripped version were generated using a custom methodology of fMRIPrep. Several confounding time-series were calculated based on the preprocessed BOLD: framewise displacement (FD), DVARS and three region-wise global signals. FD was computed using two formulations following Power, defined as the absolute sum of relative motions (Power et al. 2014), and Jenkinson, defined as the relative root mean square displacement between affines (Jenkinson et al. 2002). FD and DVARS were calculated for each functional run using their implementations in Nipype, following the definitions in Power et al. (2014). The three global signals are extracted within the CSF, the WM, and the whole-brain masks. Additionally, a set of physiological regressors were extracted to allow for component-based noise correction (CompCor, Behzadi et al. 2007). Principal components are estimated after high-pass filtering the preprocessed BOLD time-series (using a discrete cosine filter with 128 s cut-off) for the two CompCor variants: temporal (tCompCor) and anatomical (aCompCor). tCompCor components are then calculated from the top 2% variable voxels within the brain mask. For aCompCor, three probabilistic masks (CSF, WM and combined CSF+WM) are generated in anatomical space. The implementation differs from that of Behzadi et al. (2007) in that instead of eroding the masks by 2 pixels on BOLD space, a mask of pixels that likely contain a volume fraction of GM is subtracted from the aCompCor masks. This mask is obtained by dilating a GM mask extracted from the FreeSurfer’s aseg segmentation, and it ensures components are not extracted from voxels containing a minimal fraction of GM. Finally, these masks are resampled into BOLD space and binarized by thresholding at 0.99 (as in the original implementation). Components are also calculated separately within the WM and CSF masks. For each CompCor decomposition, the k components with the largest singular values are retained, such that the retained components’ time series are sufficient to explain 50% of variance across the nuisance mask (CSF, WM, combined, or temporal). The remaining components are dropped from consideration. The head-motion estimates calculated in the correction step were also placed within the corresponding confounds file. The confound time series derived from head motion estimates and global signals were expanded with the inclusion of temporal derivatives and quadratic terms for each (Satterthwaite et al. 2013). Frames that exceeded a threshold of 0.5 mm FD or 1.5 standardized DVARS were annotated as motion outliers. Additional nuisance time series are calculated by means of principal components analysis of the signal found within a thin band (crown) of voxels around the edge of the brain, as proposed by Patriat, Reynolds, and Birn (2017). The BOLD time series were resampled into standard space, generating a preprocessed BOLD run in MNI152NLin2009cAsym space. First, a reference volume and its skull-stripped version were generated using a custom methodology of fMRIPrep. All resamplings can be performed with a single interpolation step by composing all the pertinent transformations (i.e. head-motion transform matrices, susceptibility distortion correction when available, and co-registrations to anatomical and output spaces). Gridded (volumetric) resamplings were performed using antsApplyTransforms (ANTs), configured with Lanczos interpolation to minimize the smoothing effects of other kernels (Lanczos 1964). Non-gridded (surface) resamplings were performed using mri_vol2surf (FreeSurfer).

#### Statistical analyses

Unless otherwise stated, all analyses were run in Python 3.11. We used `nilearn` and `nibabel` for fMRI data processing; `ants` for conversions between native and MNI spaces; `scipy`, `statsmodels` and `scikit-learn` for statistical analyses; and `numpy`, `pandas`, `matplotlib` and `seaborn` for further data handling and visualization. All our codes and statistical maps are available online at https://osf.io/vtnu2 (code) and https://identifiers.org/neurovault.collection:22716 (statistical maps).

Univariate analyses

We implemented a two-level analysis. For each participant, the time series for each run were modeled by the convolution of several regressors with SPM’s canonical hemodynamic response function (HRF). The regressors included the onsets of the 15 items; each of the four button presses; a parametric modulator for response time; the onsets of the feedback periods (kernel = duration of the whole period); the first item of each block (because it could not be compared to a previous item); the six head motion parameters; the CSF and white matter signal, 10 anatomical CompCor (computed by fMRIprep); 20 cosine drifts; and a constant. We then defined specific contrasts, either comparing the activation elicited by any two subsets of items, or evaluating the impact of a continuous variable such as item length

For the second-level group analysis, individual contrast images for each of the experimental conditions relative to rest were smoothed with an isotropic Gaussian filter of 6 mm FWHM and entered into a second-level model: using afni `3dLME2` function, the beta values in each voxel analysed using a mixed linear model (independently for each voxel) with fixed effects of category (integers, fractions or shapes) and word length (centered around 0) and a random intercept by participant. This was done as an attempt to control for the effects of word length in the contrasts of interest.

All brain activations were corrected for multiple comparisons using a FDR threshold of *q* < .05.

Arithmetic versus geometry ROIs

Subject-specific ROIs were defined using a combination of anatomical parcels and functional localizer contrasts. For each participant, all HCP-MMP1.0 atlas regions (Glasser et al. 2016) along the bilateral IPS and ITG were considered (48 regions in total, see supplementary material S6). Within each region, for each participant, the voxels showing significant activation at *p* < .001 (uncorrected) for the arithmetic statements > geometric statements contrast in the sentence localizer were selected, and similarly for the geometric statements > arithmetic statements, producing a total of 96 possible ROIs per participant (48 arithmetic-preferring and 48 geometry-preferring). Individual ROIs containing less than 15 voxels were discarded, meaning that all ROIs were not present in all participants. In subsequent analyses, ROIs that were only present in a single participant were also discarded, yielding a total of 54 regions (to maximize sensitivity afforded by 7 Tesla fMRI, regions present in two or more participants were retained). IPS and ITG were retained as a priori regions of interest as they are the only regions in which Amalric and Dehaene (2016) found a difference between geometry and other math domains.

Mean beta values for each item in the main task were extracted for each ROI. All ROIs in which the mean beta across items was greater than zero were then tested individually: to assess the effect of category (integer, fraction or shape) on the mean beta values, two mixed linear models with mean beta value as dependent variable were fitted, one with a fixed effect of category and a random intercept by participant, and one with fixed effects of category and word length and a random intercept by participant. To assess the presence of a category effect while controlling for word length, the two models were compared using a likelihood ratio test, and results were corrected for multiple comparisons across the 54 regions in which the mean beta across items was non-negative, using false discovery rate (FDR) correction. For regions showing a significant effect of category while controlling for word length, we also report the *Z*-score and p-values for the contrasts integers + fractions > shapes and integers > shapes computed from the beta ∼ category + length + (1|participant) model.

Representational Similarity Analysis (RSA)

We performed two representational similarity analyses (RSA) (Kriegeskorte 2008). In ROI-based RSA, we directly tested the effects of the various properties of the items (the detailed models are described below) in six subject-specific regions of interest. In searchlight RSA, we instead considered a sliding searchlight sphere centered around each voxel (radius = 5 voxels), allowing us to probe the effects of our predictors across the whole brain. The analyses were implemented using a custom version of the classes and methods provided by Python’s rsatoolbox.

For the ROI-based RSA, the ROIs were defined using the functional localizer, with a contrast aimed at isolating brain areas preferentially engaged during mathematical reasoning. Specifically, we computed the contrast arithmetic statements + geometric statements > general knowledge + contextual knowledge, thresholded at *p* < .001 uncorrected. To control for potential confounding variables, we included the same regressors of non-interest as those used in the similarity task: six motion parameters, CSF and white matter signal, ten anatomical CompCorr, twenty cosine drifts and a constant. For each individual participant, we then generated subject-specific mathematical ROIs by intersecting the thresholded individual statistical map of the math

> non-math contrast from the localizer (*p* < .001 uncorrected) with anatomically defined masks for six a priori regions based on previous studies (Amalric and Dehaene 2016, 2018, 2019): left and right IPS, ITG, and dorsolateral prefrontal cortices (dlPFC). Since the thresholding did not ensure equal number of voxels in each region across participants, we report the statistics about ROI sizes in Table 2.

**Table 2.** Statistics about the number of voxels in each subject-specific math ROI.

| Region | Mean | STD | Min | 25% CI | Median | 75% CI | Max |
| --- | --- | --- | --- | --- | --- | --- | --- |
| Left IPS | 2898 | 1271 | 482 | 2078 | 2644 | 3337.25 | 5897 |
| Right IPS | 2346 | 1175 | 107 | 1657 | 2229 | 3110.5 | 4495 |
| Left ITG | 508 | 277 | 63 | 293.75 | 521.5 | 624.5 | 1173 |
| Right ITG | 311 | 180 | 33 | 138 | 285.5 | 463.5 | 635 |
| Left dIPFC | 1520 | 778 | 131 | 1032 | 1441 | 1942 | 3008 |
| Right dIPFC | 1050 | 594 | 60 | 587 | 938 | 1331.5 | 2203 |

For each participant, we started by estimating the beta coefficients of each voxel’s response to a given item. For each subset of voxels (ROI or searchlight sphere), we combined these coefficients into an item-specific vector of spatial activations within the region. We then compared the resulting 15 vectors – one per item – pairwise using correlation distance (1-Pearson’s *r*), yielding a 15×15 representational dissimilarity matrix (RDM).

Three successive models were fit to the neural RDMs (see predictor matrices on Figure S1):

● **Subjective similarity model:** The behavioral model examined which areas showed neural RDMs that matched the behavioral semantic similarity ratings (averaged across subjects, as collected during fMRI acquisition). To separate them from visual similarity, we also included a regressor for the absolute difference in word length on a logarithmic scale, i.e. ∣log(len(i1))−log(len(i2))∣ where len(i1) and len(i2) are the lengths of items i1 and i2, respectively.
● **Category model:** We next investigated which areas encoded the two main categories of concepts: numbers versus shapes. Thus, in addition to the word length similarity predictor as above, one regressor predicted a greater similarity between targets when both were numbers (fractions or integers), and another when both were shapes (except segment). Note that we decided to assign the item “segment” to its own category and exclude it from the other geometric shapes. This decision stemmed from the observation that (1) a segment is not a 2D shape, and (2) “segment” was consistently distinguished from the other shapes in our data. A likely explanation is that most shapes in our set are perceived as two-dimensional (for example, a triangle has an area), whereas a segment is inherently one-dimensional. Consequently, the two endpoints defining a segment are less visually salient and are not corners, unlike the three corners of a triangle or the four of a square. This difference may also weaken the intuitive link between “segment” and number 2, compared with the stronger correspondence between “triangle” and 3. Therefore, we treated “segment” as a distinct case in our analyses.
● **Full model:** The full model aimed at testing specific dimensions of semantic similarity. It differed from the simple model in two ways (see Figure S1). First, it encoded item categories more precisely: the binary predictor for numbers, which did not distinguish between fractions and integers, was broken down into three binary predictors which respectively responded to integer-integer pairs, integer-fraction or fraction-integer pairs, and fraction-fraction pairs. Second, the model aimed at encoding conceptual similarity both within and across categories, based on the number associated with each item (e.g., “three”, “triangle” or “third” are all associated with 3, “four”, “square” or “fourth” are all associated with 4). Within each category, a continuous predictor tested for a distance effect, encoded the logarithm of the distance between numbers (log(1+|num(i1)-num(i2)|) for items i1 and i2); an additional predictor was also included for a semantic distance effect between integers and fractions. Across categories, binary predictors simply encoded whether the two items were associated with the same number, with one predictor for each pair of categories (integer-fraction, integer-shape, and fraction-shape). Other predictors from the simple model were left untouched.

For all three models, we initially included an additional predictor of frequency (|logfpm(i1)-logfpm(i2)| where logfpm denotes log frequency per million in the math corpus from Debray & Dehaene (Debray and Dehaene 2025). However, as expected from Zipf’s law (Piantadosi 2014; Zipf 2013), length and frequency were closely related (across our 15 words, the correlation between log frequency per million and log length was *r*(13) = −0.61, *p* = .02). We therefore retained only the word length predictor (but the results were very similar when we kept both).

The similarity matrices for each of the three models were fitted to the empirical ones using multiple linear regression, providing us with one beta weight per predictor, per participants, and per ROI (or searchlight sphere). We then tested each predictor for significance across participants: in ROI-based RSA, we simply performed, for each ROI, a group-level one-sample *t*-test on the regression beta coefficients. Since the predictors were designed such that the similarities predicted by our theory should yield positive beta values, only positive betas were tested for significance and are reported here. In searchlight RSA, we attached each beta coefficient to the voxel on which the sphere was centered, thus obtaining a beta-map over the whole brain. We spatially smoothed the beta-map using an isotropic Gaussian filter of 6 mm FWHM, and then performed group-level one-sample *t*-tests for each voxel of the smoothed beta-maps to test which voxels exhibited a statistically significant association to the predictors The resulting statistical maps were thresholded using a false discovery rate (FDR) correction at *q* < 0.05 to control for multiple comparisons across the brain and identify regions showing reliable effects for each predictor.

### MEG Experiment

#### Tasks

During the MEG scanning session, participants were presented with a set of 15 mathematical stimuli comprising seven integers (zéro, un, deux, trois, quatre, cinq, six), three fractions (demi, tiers, quart), and five geometric shapes (segment, triangle, carré, pentagone, hexagone). Each stimulus was displayed in two formats: a written word format and a figure format, presented on a mid-gray background. Stimuli were presented for 200 ms, followed by a 300 ms blank inter-stimulus interval, resulting in a stimulus onset asynchrony (SOA) of 500 ms.

Participants completed 20 experimental runs (approximately 2 min per run), consisting of 10 runs with written word stimuli and 10 runs with figure stimuli. During each run, participants performed an intruder concept detection task in which an infrequent, non-mathematical intruder stimulus appeared at random intervals. The intruders were animal or object words in the word blocks, and letters in the figure blocks. Participants were instructed to press a response button as quickly as possible upon detecting intruder stimulus (see Figure S4 for all stimuli and intruders in both formats).

#### Participants

Thirty healthy adult participants (17 female) took part in the MEG experiment. All participants held at least a bachelor’s degree in a scientific discipline. Participants reported normal or corrected-to-normal vision and no history of neurological or psychiatric disorders. All provided written informed consent prior to participation, and the study was conducted in accordance with the Declaration of Helsinki and approved by the local ethics committee.

#### MEG parameters

Subjects’ brain activity was recorded at Neurospin, Gif-sur-Yvette, using a MEG system comprising 306 channels (204 planar gradiometers and 102 magnetometers) manufactured by Elekta-Neuromag Ltd. in Helsinki, Finland. The data were acquired at a sampling rate of 1000 Hz and underwent online band-pass filtering within the frequency range of 0.01–330 Hz. The participants were seated upright within a room that was shielded against magnetic interference (AK3B, Vakuumschmelze, Hanau, Germany). Prior to the recording session, the unique head shape of each participant was measured using a Polhemus FASTRAK 3D digitizer (Polhemus, Vermont, USA). This process involved acquiring three fiducial points (nasion and pre-auricular points), five head position indicator (HPI) coils (one on each of the left and right mastoids and three on the forehead), and approximately 100 more locations distributed across the paricipatns’ skull. At the start of each run, head positioning inside the MEG helmet was measured by inducing a non-invasive current through the HPI coils. Additionally, both horizontal and vertical electro-oculograms (EOGs) were recorded, along with an electrocardiogram (ECG), for later offline removal of artifacts related to eye movements and heart activity. For stimulus presentation and control, Expyriment (Krause & Lindemann, 2013) was employed. Each visual stimulus image was rear-projected onto a screen positioned at a distance of 120 cm from the participant’s eyes using a VPixx PROPixx projector.

#### MEG data preprocessing

The offline raw MEG data underwent visual inspection to identify and eliminate noisy channels. Subsequently, we employed the MaxMove function of the Elekta Maxfilter software to remove head motion and to denoise the data using Maxfilter Signal Space Separation (Taulu, Kajola, and Simola 2004). Following the maxfiltering, we proceeded with additional preprocessing using MNE-Python version 0.24.1 (Gramfort 2013). First, the data underwent band-pass filtering within the range of 0.05 to 330 Hz and bandstop filtering at 50 Hz and its harmonics for line noise removal. To eliminate artifacts caused by eye movement and heart rate, independent components of the MEG data were computed and correlated with the EOG and ECG signals. Components displaying significant correlations were subsequently removed through manual inspection. Subsequently, to enhance signal-to-noise ratio (SNR) and lower computational demands, we downsampled the data to 200 Hz (Grootswagers, Wardle, and Carlson 2017). Following these preprocessing steps, the data was epoched into 0.55-second trials, starting 50 ms before the stimulus onset. Each epoch was then normalized by subtracting the baseline period mean.

For all subsequent analyses, only planar gradiometer channels were used. Given that magnetometers and gradiometers have different measurement units, restricting the analysis to gradiometers ensures methodological consistency. This choice is consistent with prior MEG studies that exclusively analyzed gradiometer data (e.g. Karami, Castaldi, Eger, Hebart, et al. 2025; Ritchie, Tovar, and Carlson 2015). Moreover, planar gradiometers provide higher signal-to-noise ratio and improved sensitivity to superficial cortical sources compared to magnetometers, making them particularly suitable for multivariate sensor-space analyses (Hultén et al. 2021). All our codes and MEG data are available online at https://osf.io/vtnu2.

#### Statistical analyses

Time Resolved Multivariate Representational Similarity Analysis (RSA)

To examine how different stimulus properties were reflected in neural population responses over time, we employed representational similarity analysis (RSA; Kriegeskorte 2008) combined with multiple-regression modeling.

Neural representational dissimilarity matrices (RDMs) were constructed from the preprocessed MEG data by first organizing the signals into pattern vectors, each consisting of activity from all selected sensors for a given trial and time point. These vectors were then averaged across trials belonging to the same experimental condition, resulting in 15 condition-specific MEG patterns at each time point. This procedure was repeated for all time points spanning from 50 ms before to 450 ms after stimulus onset. For each time point, we computed the Pearson correlation between all pairs of the 15 condition-specific patterns and converted the resulting similarity matrix into a dissimilarity matrix by subtracting the correlation coefficients from 1, yielding a 15 × 15 neural RDM per time point. Additionally, we constructed cross-notation RDMs by correlating each condition-specific pattern from one notation with the patterns from the other notation, converting these cross-notation similarity matrices to dissimilarities (1−r). Because the resulting cross-notation matrices can be asymmetric, each was symmetrized by averaging the matrix with its transpose (equivalently, averaging the upper and lower triangles) before vectorization and inclusion in the multiple-regression RSA.

To assess the extent to which the dissimilarity structure of the MEG representations could be accounted for by different stimulus-related factors, we performed multiple-regression RSA. In this framework, the vectorized neural RDM at each time point served as the dependent variable, and multiple model (predictor) RDMs were entered simultaneously as regressors. Three classes of model RDMs were included in the regression analysis as in the fMRI experiment.

Regression coefficients obtained from the multiple-regression RSA were tracked over time to characterize when each class of stimulus information contributed to the neural representational geometry. The analysis was implemented in the `rsatoolbox` (Van Den Bosch et al. 2025) and custom-written code in Python (version 3.11).

Statistical significance was assessed using one-sample *t*-tests against zero across participants. To correct for multiple comparisons across time points, we employed threshold-free cluster enhancement (TFCE; Smith and Nichols 2009) combined with nonparametric permutation testing. Specifically, null distributions were generated using Monte Carlo simulations with 10,000 permutations. This procedure was implemented in MNE-Python. Resulting TFCE-corrected statistics were thresholded at *p* < 0.05 (one-tailed) to determine statistical significance.

MEG-fMRI Fusion with RSA

To link the temporal dynamics of neural representations measured with MEG to the spatially resolved representations obtained from fMRI, we performed RSA-based MEG–fMRI fusion (Cichy, Pantazis, and Oliva 2014). This approach assesses when time-resolved MEG representational structures resemble the representational geometry observed in specific brain regions with fMRI, indicating a shared representational format at a given moment in time.

For each participant and time point, MEG representational dissimilarity matrices (RDMs) were correlated with group-averaged fMRI RDMs derived from predefined regions of interest (ROIs). Pearson correlation coefficients were computed between the vectorized MEG RDMs and the corresponding fMRI RDMs, yielding a time course of MEG–fMRI representational correspondence for each ROI. The fMRI RDMs were obtained from our fMRI experiment, which used the same set of written stimuli as in the present MEG experiment but employed an explicit similarity-judgment task rather than passive viewing. The ROIs included early visual areas (V1–V3) and a math network comprising parietal, inferior temporal gyrus (ITG), and frontal regions.

We used group-averaged fMRI RDMs as a robust estimate of the underlying representational dissimilarity structure (Nili et al. 2014) for the MEG–fMRI fusion analysis. This procedure allowed us to track when MEG representations converged with region-specific fMRI representational structures, despite differences in task demands between the two experiments.

Multidimensional Scaling (MDS)

While RSA provides a hypothesis-driven assessment of how specific stimulus features explain variance in the neural data, we additionally examined the latent structure of the neural representations using a data-driven approach based on multidimensional scaling (MDS; Kruskal 1964). MDS was applied directly to the representational dissimilarity matrices (RDMs) to visualize the relational structure among stimuli without imposing prior assumptions about relevant dimensions. For visualization, we projected the neural dissimilarities onto a two-dimensional space defined by the first two dimensions of the MDS solution. In these plots, the distances between stimuli reflect the dissimilarity of the neural response patterns they elicited, such that stimuli positioned closer together evoke more similar neural representations (Nili et al. 2014). The MDS analysis was performed using the `rsatoolbox` on group-averaged neural RDMs. Group-level RDMs were obtained by averaging individual participants’ RDMs across all subjects, yielding a more stable and noise-reduced estimate of the underlying representational structure. MDS was applied to RDMs spanning multiple time points following stimulus onset, allowing us to visualize how the organization of neural representations evolved over time.

## Acknowledgements

We thank Antonio Moreno for sharing his stimuli for the localizer; Thomas Dighiero-Brecht, Antoine Grigis, Alexander Paunov, Manon Pietrantoni, and Bosco Taddei for sharing scripts and insights on data analysis; Minye Zhan for helping us set up the fMRI acquisition sequences; and the MRI technicians, the doctors and the nurses from NeuroSpin’s Biomedical Research Imaging Cell for their support with participant recruitment and fMRI & MEG acquisitions.

## Data and code accessibility

All the code related to this study is available through OSF: https://doi.org/10.17605/OSF.IO/VTNU2 The MEG data can be found in the same OSF repository. Due to French CEA regulations, the raw fMRI data could not be made public, but the second level statistical maps are available on NeuroVault: https://identifiers.org/neurovault.collection:22716

## Declaration of generative AI and AI-assisted technologies in the writing process

The authors used ChatGPT to assist with rephrasing certain sentences during the preparation of this work. They subsequently reviewed and edited the content as needed and took full responsibility for the final published article.

## Supplementary Materials

S2 - Table of peak coordinates for the math > non-math contrast (in the localizer)

S3 - Table of peak coordinates for the numbers > shapes contrast

S4 - Table of peak coordinates for the item length contrast

S5 - Table of peak coordinates for the integers > fractions contrast

S6 - List of HCP-MMP1.0 regions along bilateral IPS and ITG retained for subject specific ROI analyses

S7 - Beta plots for the arithmetic vs geometry analysis

S8 - Table of peak coordinates for the searchlight map of behavioral RDM

S9 - Table of peak coordinates for the searchlight map of log of difference of item lengths RDM

S10 - List of stimuli of MathLang task

S12 - Video of RDMs (Written Words)

S13 - Video of RDMs (Figures)

S14 - Video of RDMs (Cross-notation)

S15 - Video of MDS (Written Words)

S16 - Video of MDS (Figures)

## Supplementary Figures

**Figure S1.**
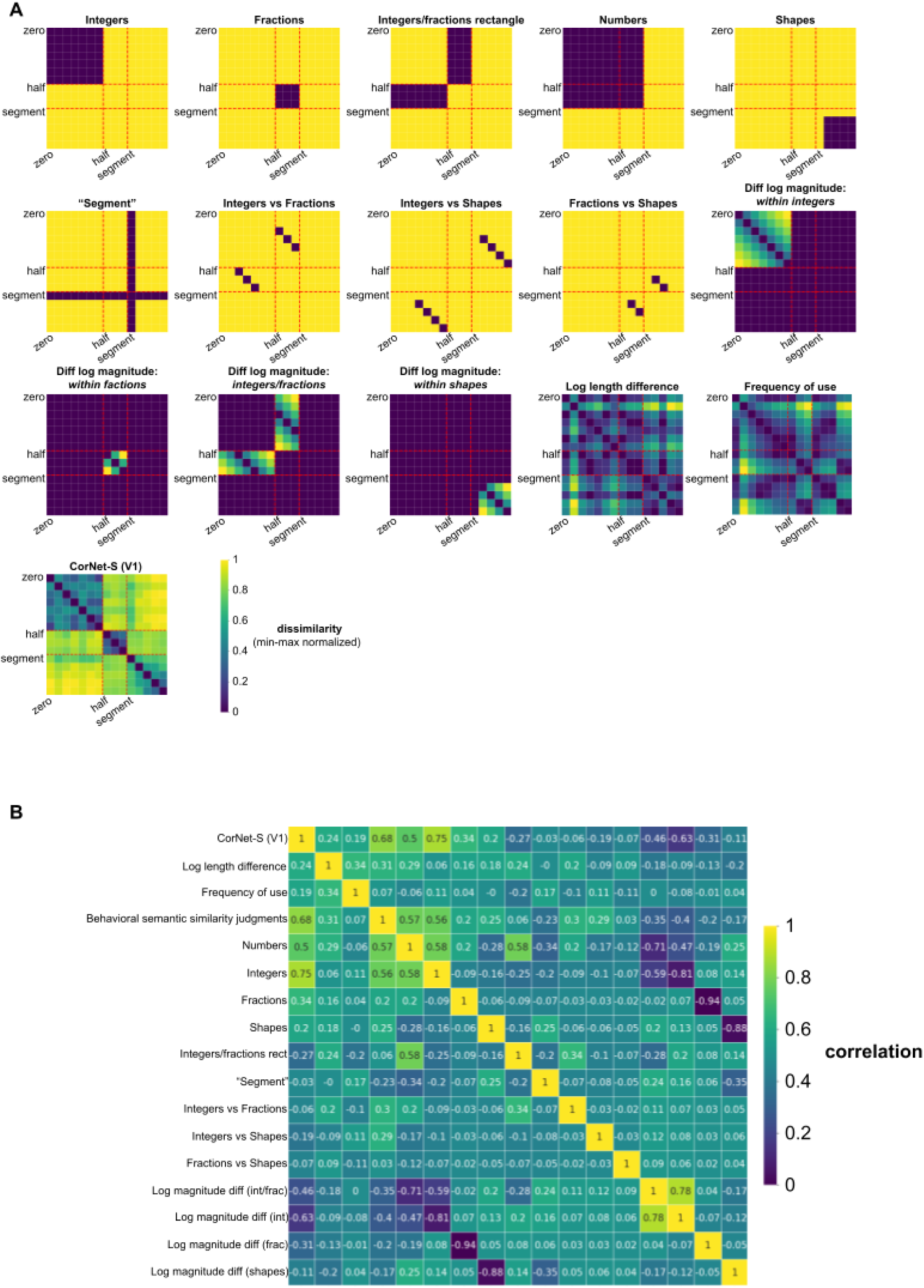
(fMRI and MEG theoretical RDMs)

**Figure S2.**
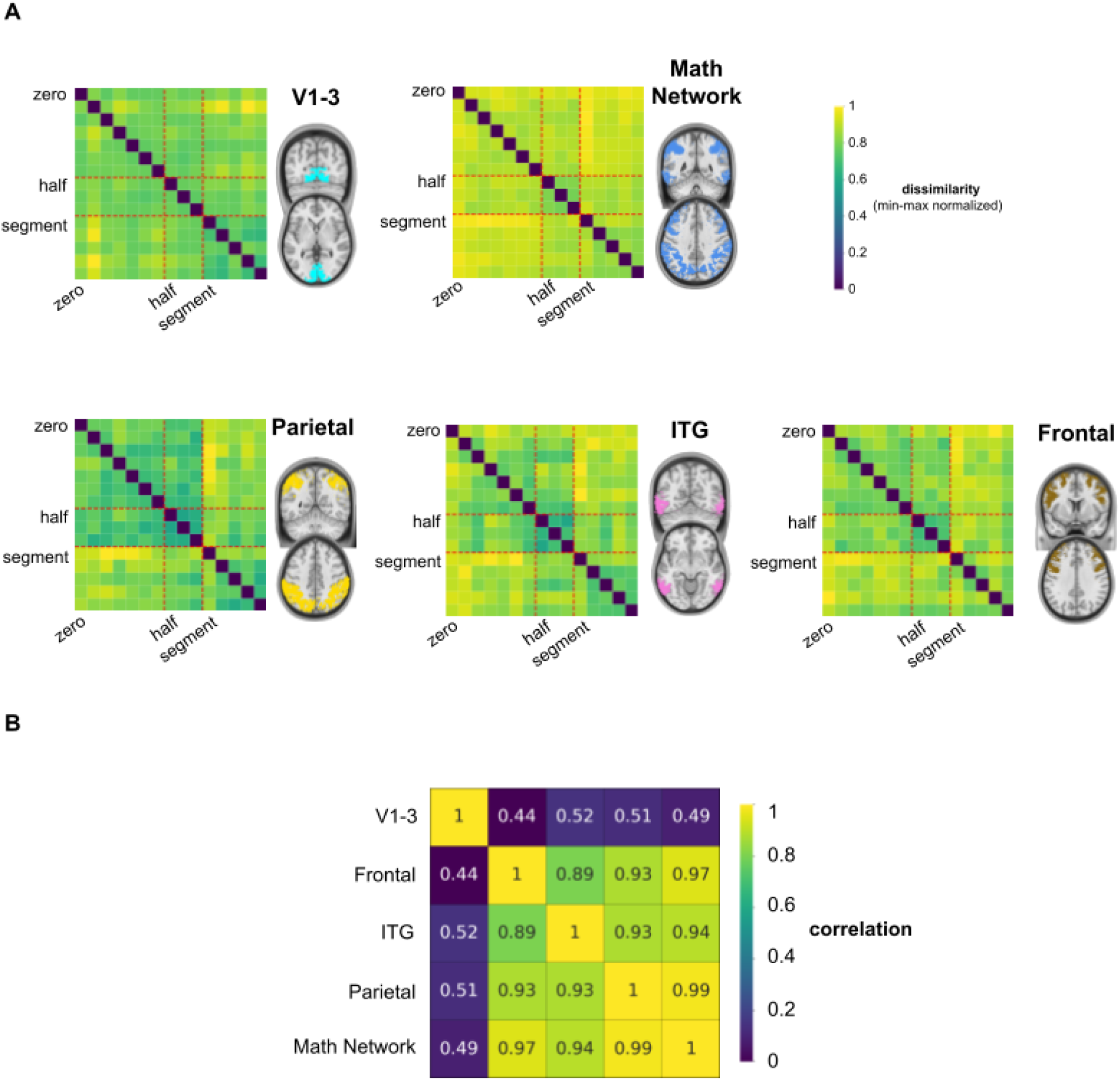
(Math network RDMs)

**Figure S3.**
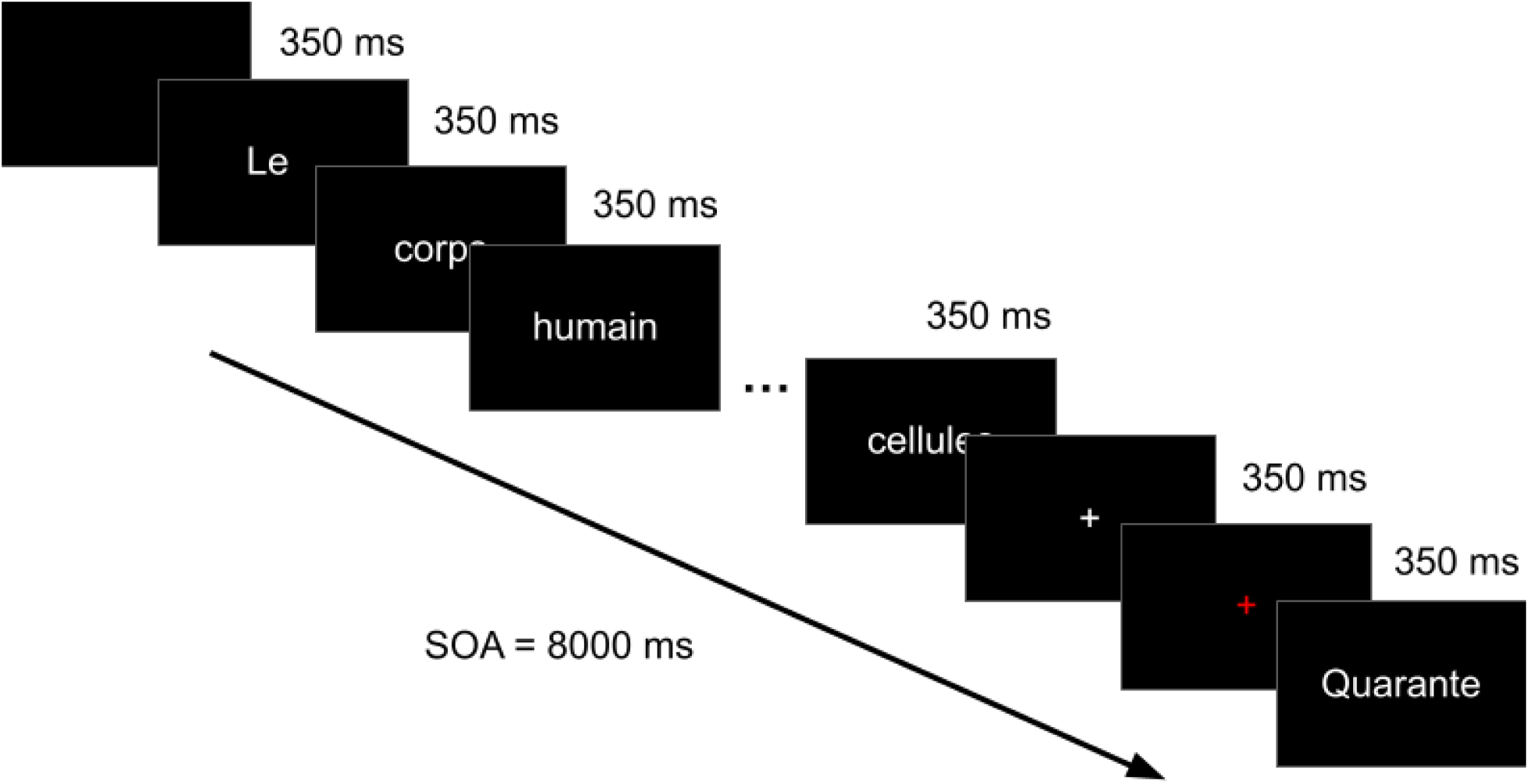
(MathLang design)

**Figure S4.**
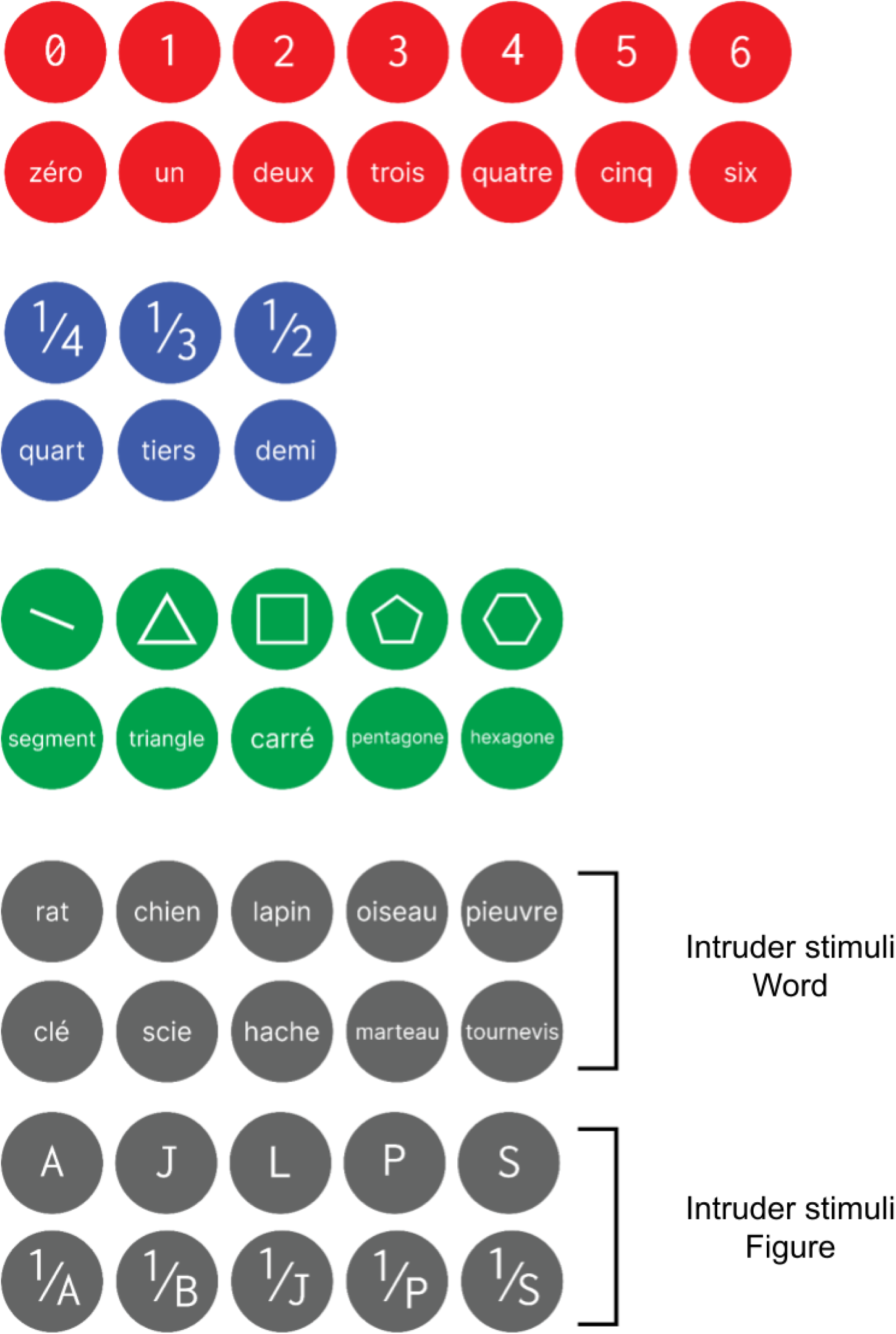
(List of Stimuli Used in fMRI and MEG Studies)

