## Supplementary material for "Cortical localization and dynamics of elementary mathematical concepts": S10

### List of stimuli of the localizer task

#### **Mathematical sentences^[1]^**

##### **Arithmetic facts**

###### **True statements**

1. Forty-five is the result of fifteen times three.
2. The result of twenty-seven minus three is the number twenty-four.
3. The result of seventy-nine minus five is seventy-four.
4. The result of multiplying six by ten is sixty.
5. The square root of thirty-six is greater than five.
6. The number twelve multiplied by ten equals one hundred twenty.
7. The sum of fifty-five and two gives fifty-seven.
8. Adding eighteen and three makes twenty-one.
9. Forty-two minus one gives the number forty-one.
10. The number twenty-seven divided by three gives nine as a result.

###### **False statements**

1. The number seventy-four is greater than one hundred.
2. The result of twenty-three times two is the number nine.
3. Forty divided by two equals ninety-eight.
4. Eighty-five is smaller than forty-four.
5. Twenty-two is the square of seventeen.
6. Seven times seven is greater than fifty-four.
7. The number nineteen is between one and seventeen.
8. Twenty-seven minus two minus four equals eighteen.
9. Thirty-seven is half of fifty-four.
10. Five plus sixteen is greater than twenty-eight.

##### **Geometric facts**

###### **True statements**

1. The diagonal of a square is longer than its side.
2. The diameter of a circle is twice its radius.
3. The area of a rectangle is the product of its sides.
4. The sides of an equilateral triangle have the same length.
5. An infinite number of lines can pass through a single point.
6. A square is a rectangle with equal sides.
7. All points on a circle are equidistant from the center.
8. An equilateral triangle can be divided into two right triangles.
9. The diagonals of a rhombus are its axes of symmetry.
10. The angles of a regular polygon are equal in size.

###### **False statements**

1. A triangle is a figure whose sides are all equal.
2. The sides of a rhombus can have different lengths.
3. Lines are perpendicular when they form an acute angle.
4. A hexagon is a figure with right angles.
5. A rectangle has only acute angles.
6. A circle has many straight sides.
7. A rectangle cut by a straight line produces equal squares.
8. A rectangle has only one axis of symmetry.
9. Perpendicular lines form an acute angle.
10. A right angle is greater than an obtuse angle.

#### **Non-mathematical sentences**

##### **General knowledge**

###### **True statements**

1. Photosynthesis occurs thanks to light energy.
2. The human body is made up of billions of cells.
3. Snowflakes have very diverse shapes.
4. Mitosis is the process by which cells divide.
5. The whale is a large mammal that lives in the sea.
6. Classical tragedy is based on unity of place.
7. Caterpillars are insects that transform into butterflies.
8. Water freezes at zero degrees Celsius.
9. The rose is the flower of a thorny plant.
10. The sun rises in the east every morning.

###### **False statements**

1. The young of a cow is called a foal.
2. The squirrel is the largest land animal in America.
3. Dinosaurs are mammals that disappeared.
4. Jupiter is the smallest planet in the solar system.
5. Green algae are aquatic plants.
6. A calf is the young of a goat.
7. The Earth rotates on itself in one month.
8. French sentences are read from right to left.
9. Winter is the hottest season of the year.
10. The Earth revolves around the planet Venus.

##### **Contextual knowledge**

###### **True statements**

1. In New York, buildings are tall modern structures.
2. In the Netherlands, windmills operate using wind.
3. In Beijing, pollution is a major problem.
4. In Ireland, having red hair is quite common.
5. According to the Bible, the universe was created in a few days.
6. In London, cars drive on the left side of the road.
7. In San Francisco, temperatures are mild.
8. In the sea, mammals have fins and a blowhole.
9. In the Caribbean, restaurants often serve fresh fish.
10. In London, buses have multiple levels.

###### **False statements**

1. In the Amazon, spiders are harmless animals.
2. In North Korea, people are free to express themselves.
3. In Kenya, lakes in the plains regularly freeze.
4. In Burundi, medical care is reimbursed by social security.
5. In India, cows are raised for meat.
6. In South Africa, diamonds are worthless stones.
7. In the United States, university education is free.
8. In Paris, the Milky Way is often visible at night.
9. In Brittany, temperatures often reach extreme highs.
10. On the planet Mars, the moon causes tides.

1. The original stimuli were presented in French, but have been translated into English in accordance with bioRxiv’s policy requiring all submitted materials to be in English.
