## Supplementary material for "Cortical localization and dynamics of elementary mathematical concepts": S7

Preference: arith > geom

### Region: Anterior_IntraParietal_Area_R (arith > geom)

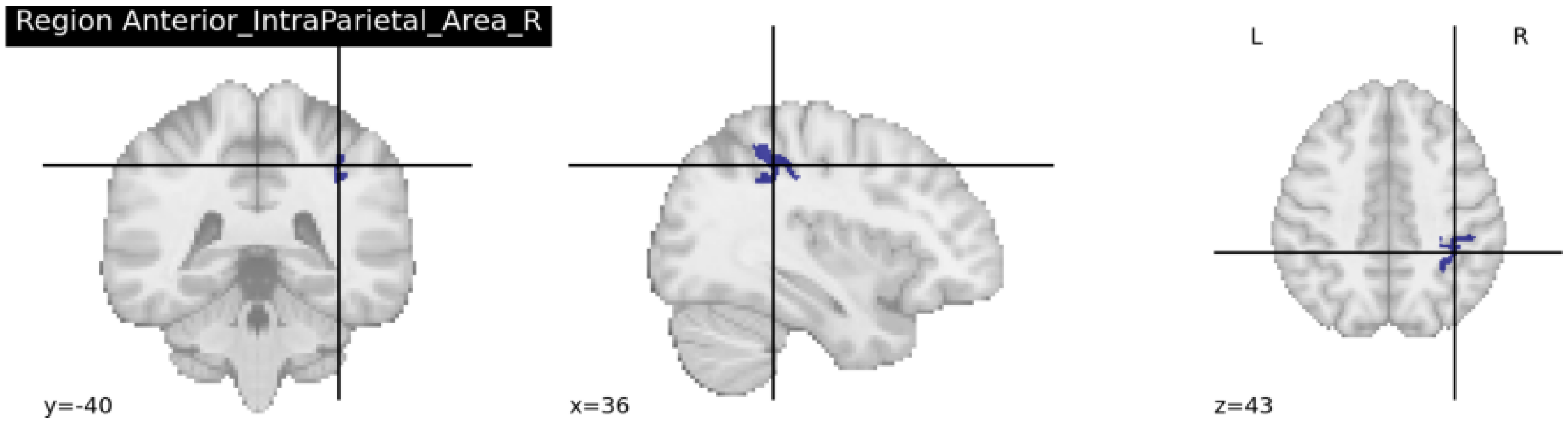


Average region size: 41 voxels (across 7 participants)
chi2-test between the two models: chi2(2) = 15.29, p = 4.313e-03 (FDR corrected)

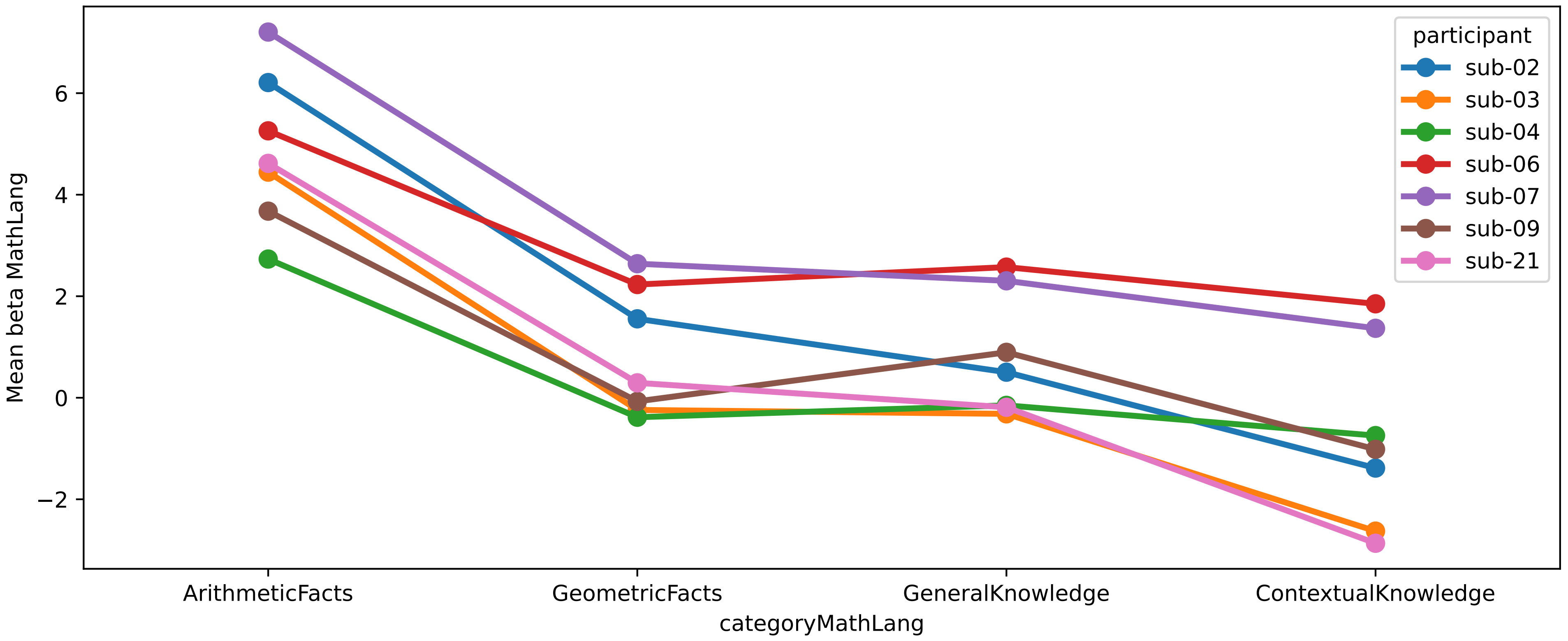


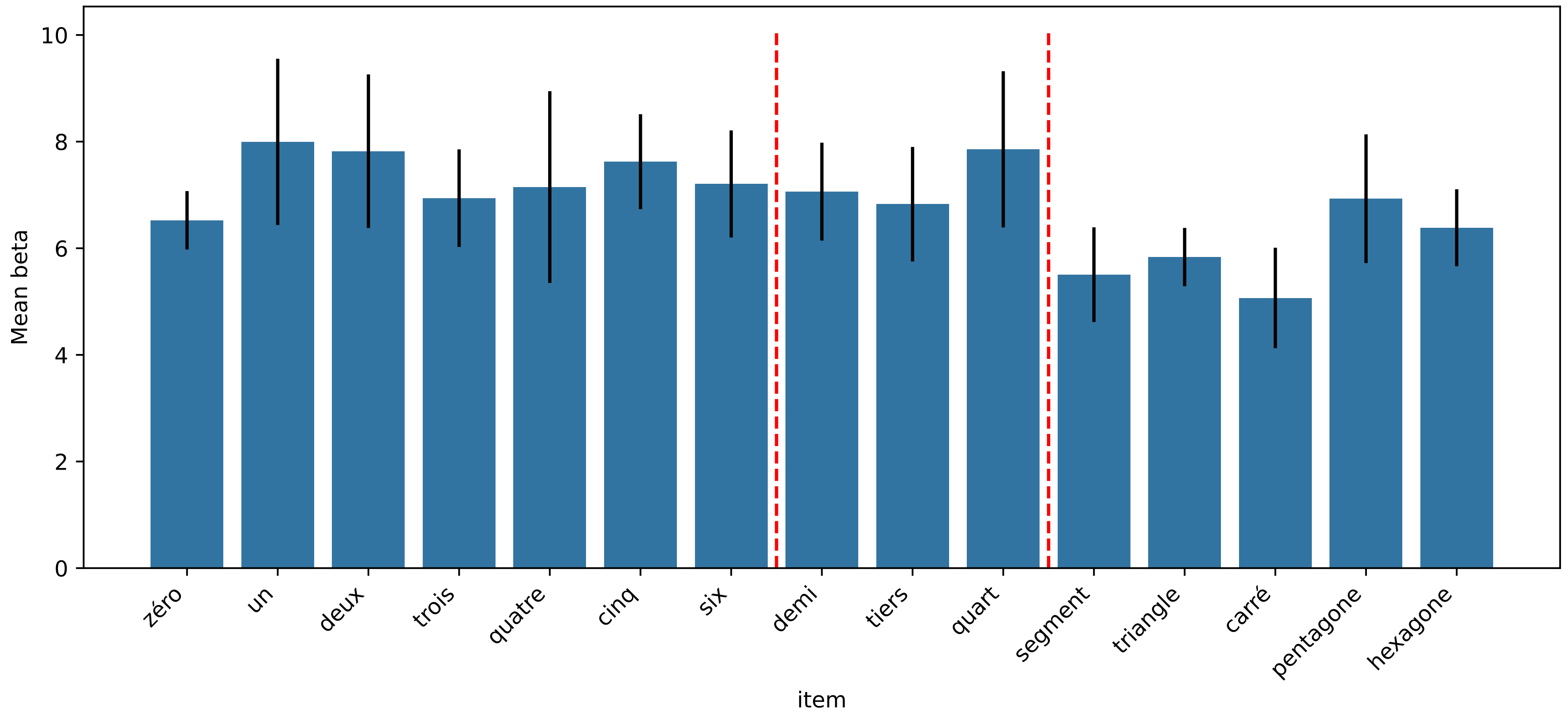

### Region: Area_TE1_posterior_L (arith > geom)

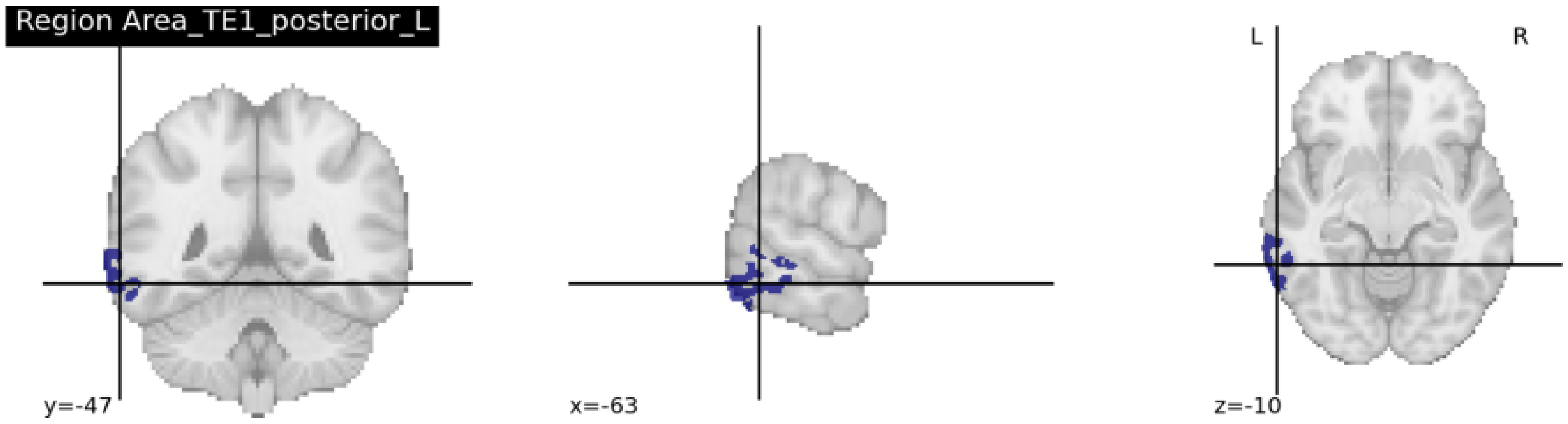


Average region size: 35 voxels (across 10 participants)
chi2-test between the two models: chi2(2) = 11.35, p = 2.055e-02 (FDR corrected)

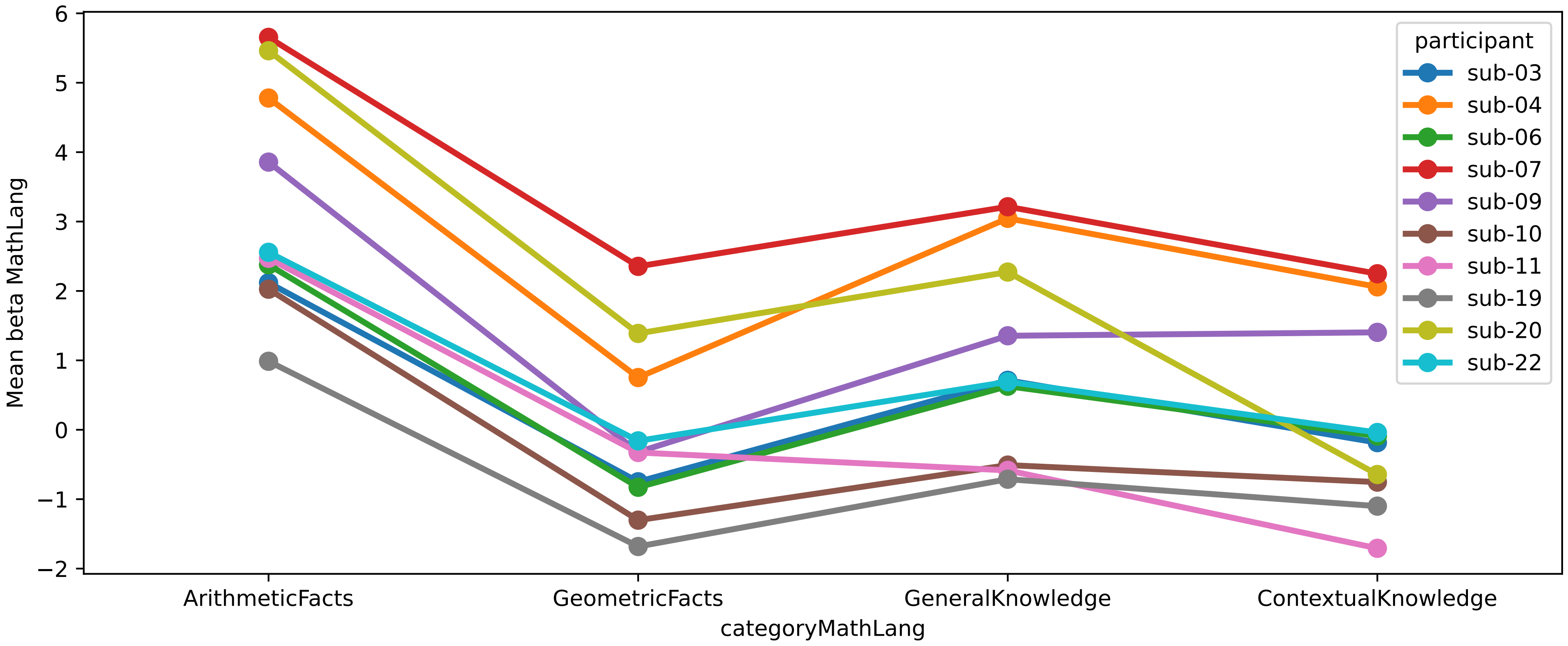


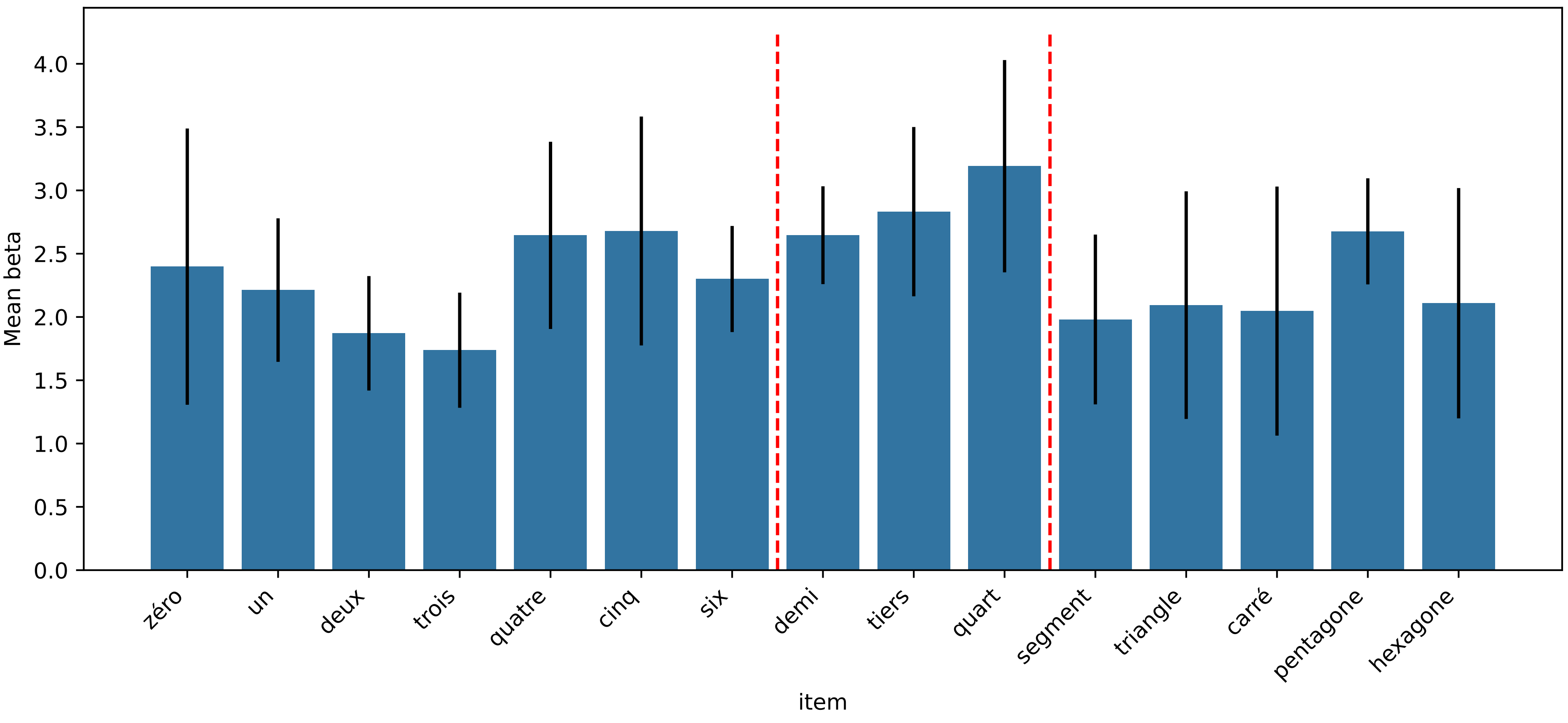

Preference: geom > arith

### Region: Area_IntraParietal_1_L (geom > arith)

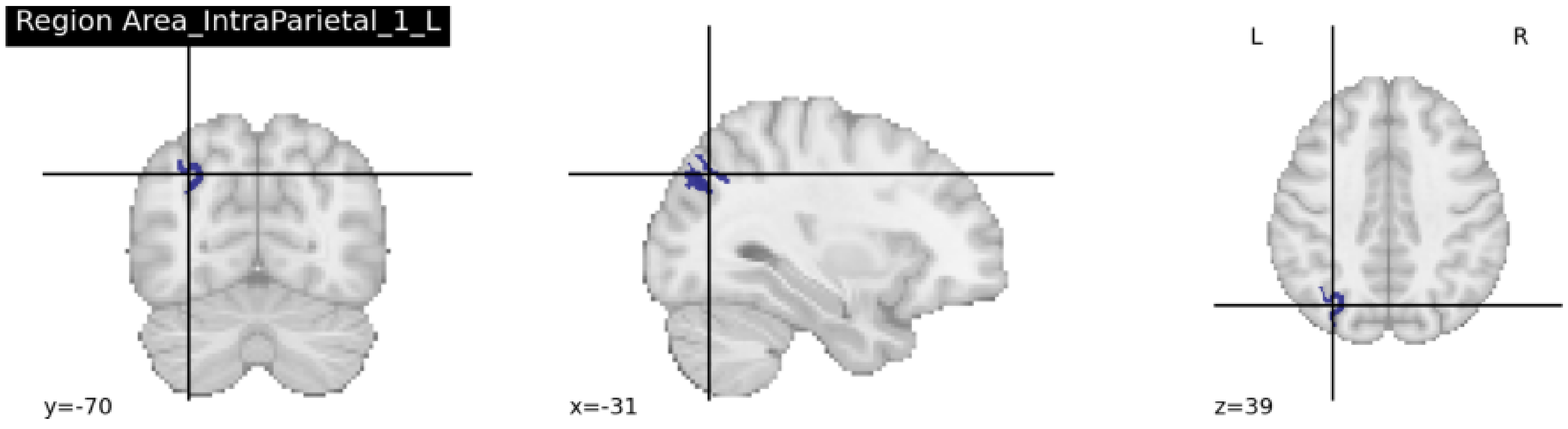


Average region size: 32 voxels (across 8 participants)
chi2-test between the two models: chi2(2) = 24.80, p = 4.437e-05 (FDR corrected)

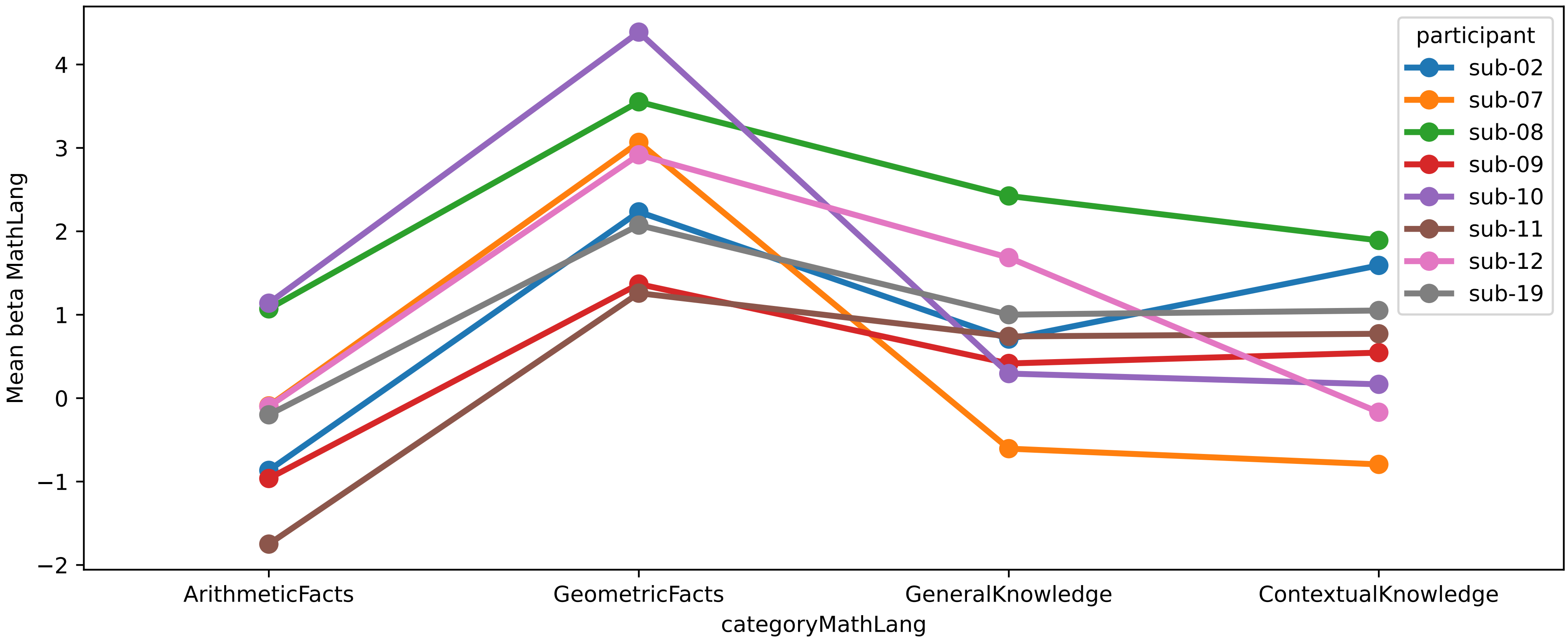


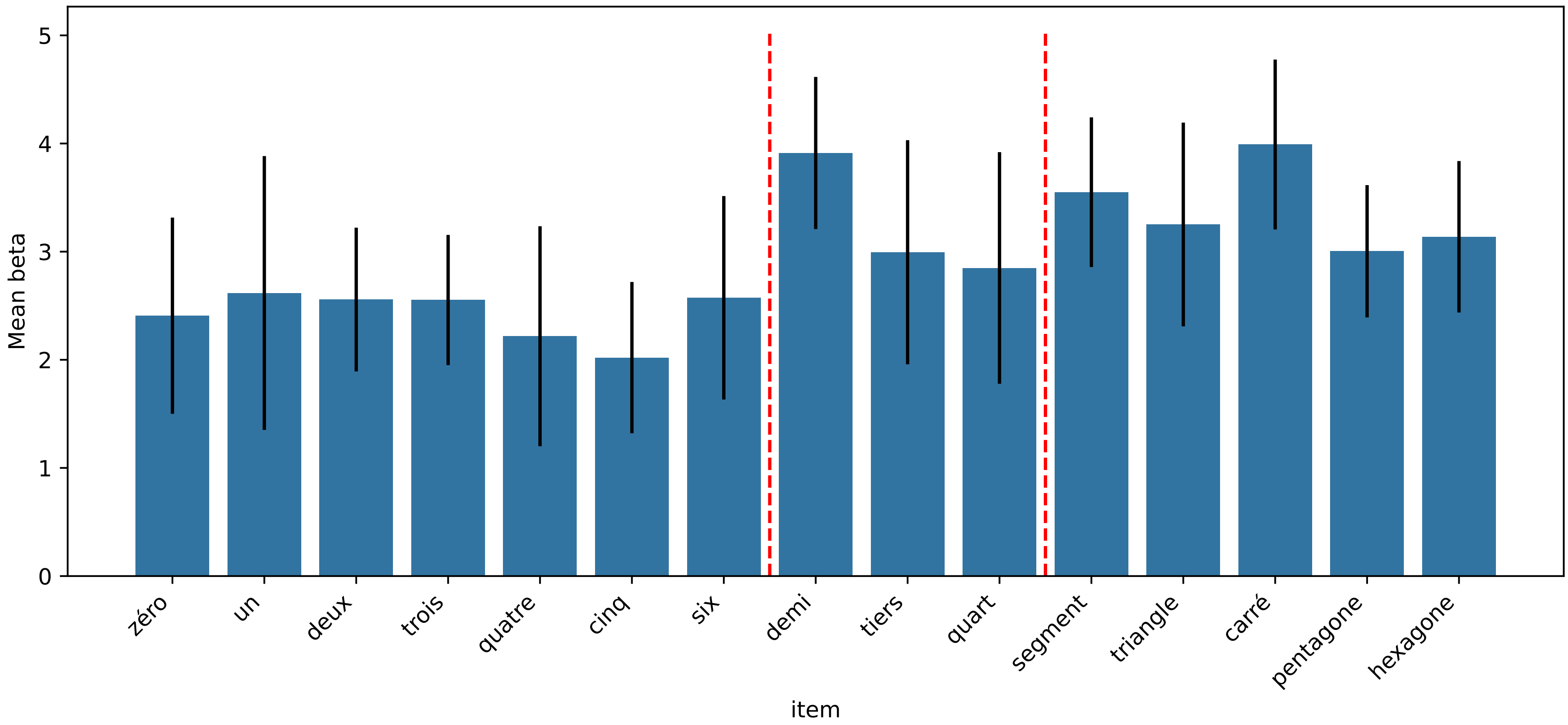

Region: Area_IntraParietal_1_R (geom > arith)

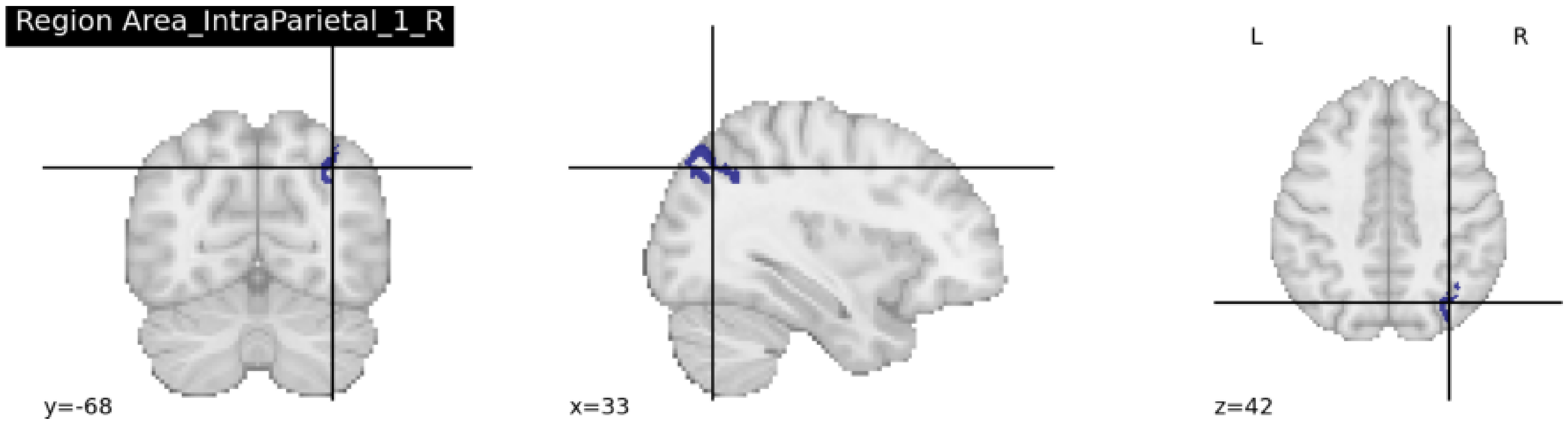


Average region size: 56 voxels (across 4 participants)
chi2-test between the two models: chi2(2) = 13.80, p = 7.792e-03 (FDR corrected)

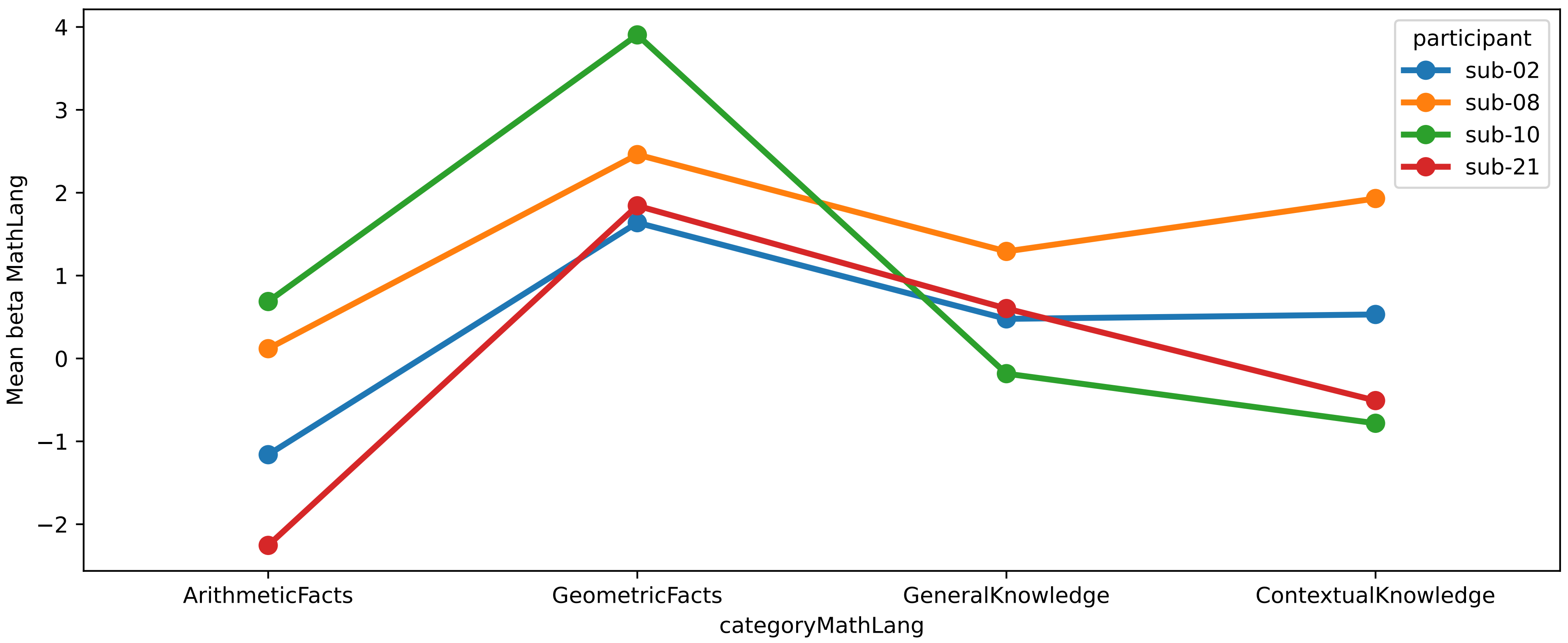


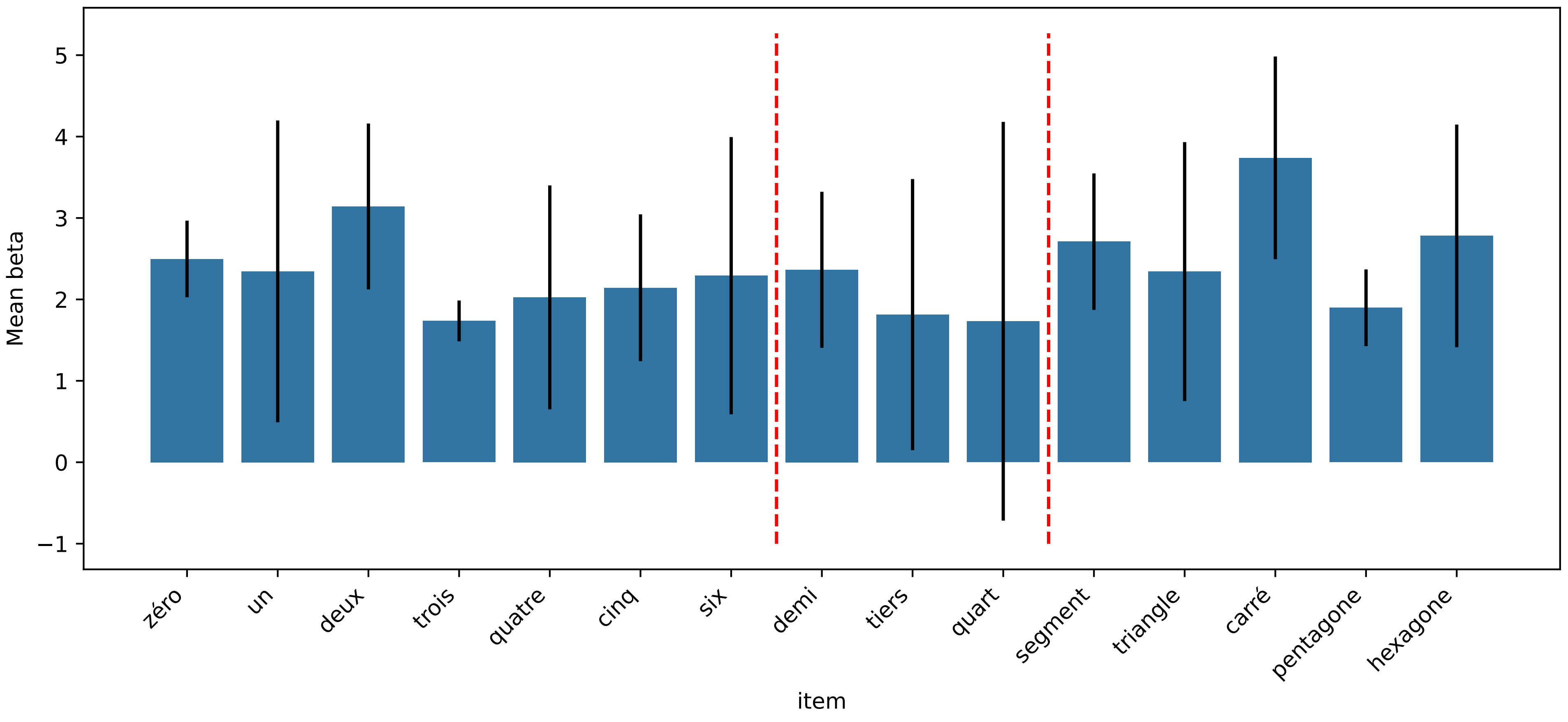

### Region: Area_PHT_L (geom > arith)

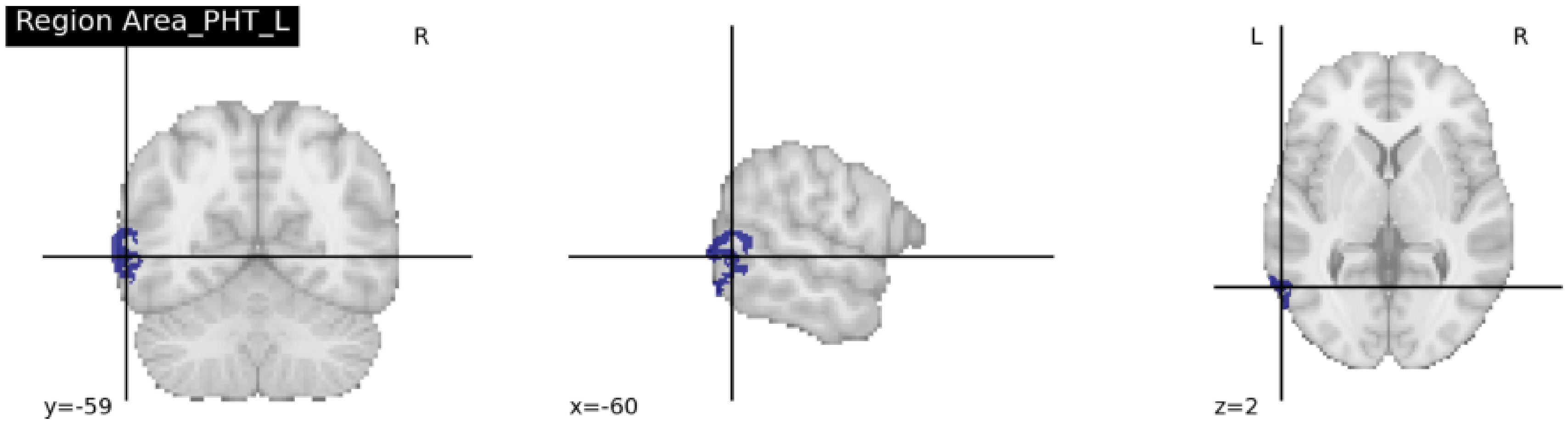


Average region size: 91 voxels (across 13 participants)
chi2-test between the two models: chi2(2) = 34.96, p = 4.613e-07 (FDR corrected)

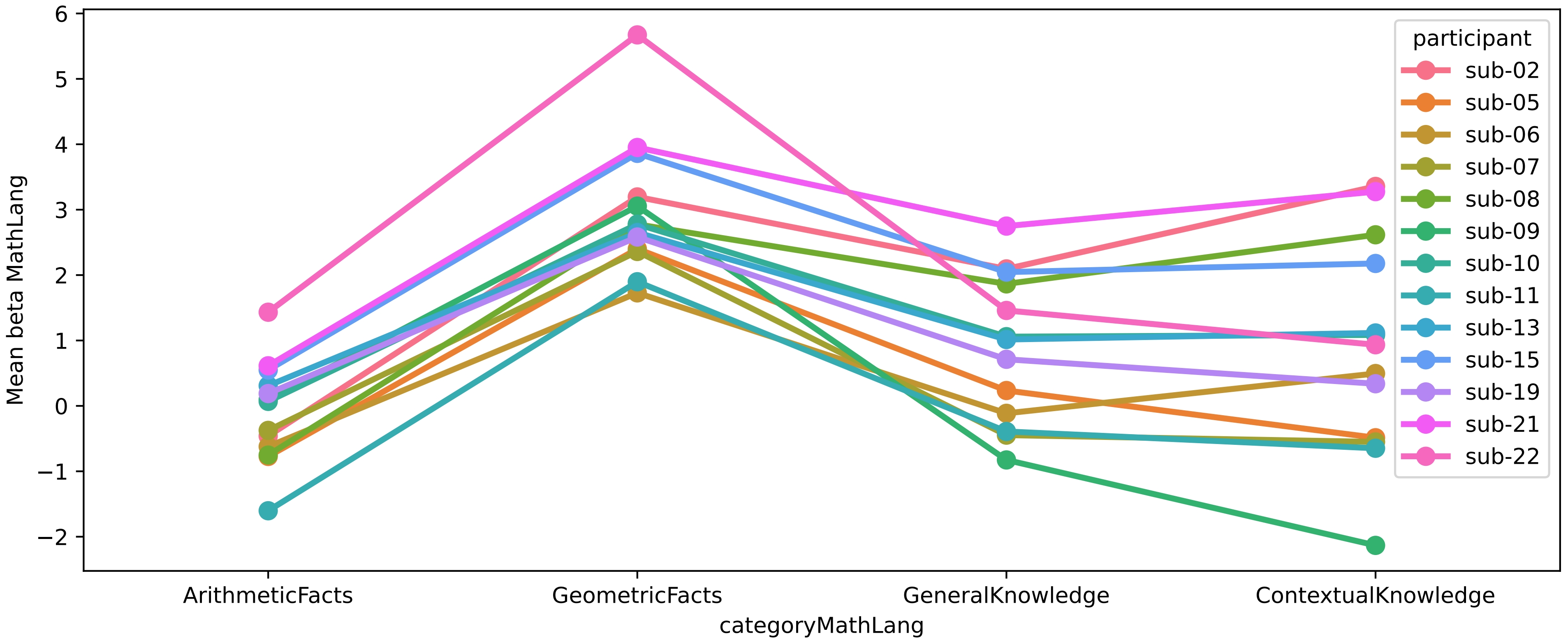


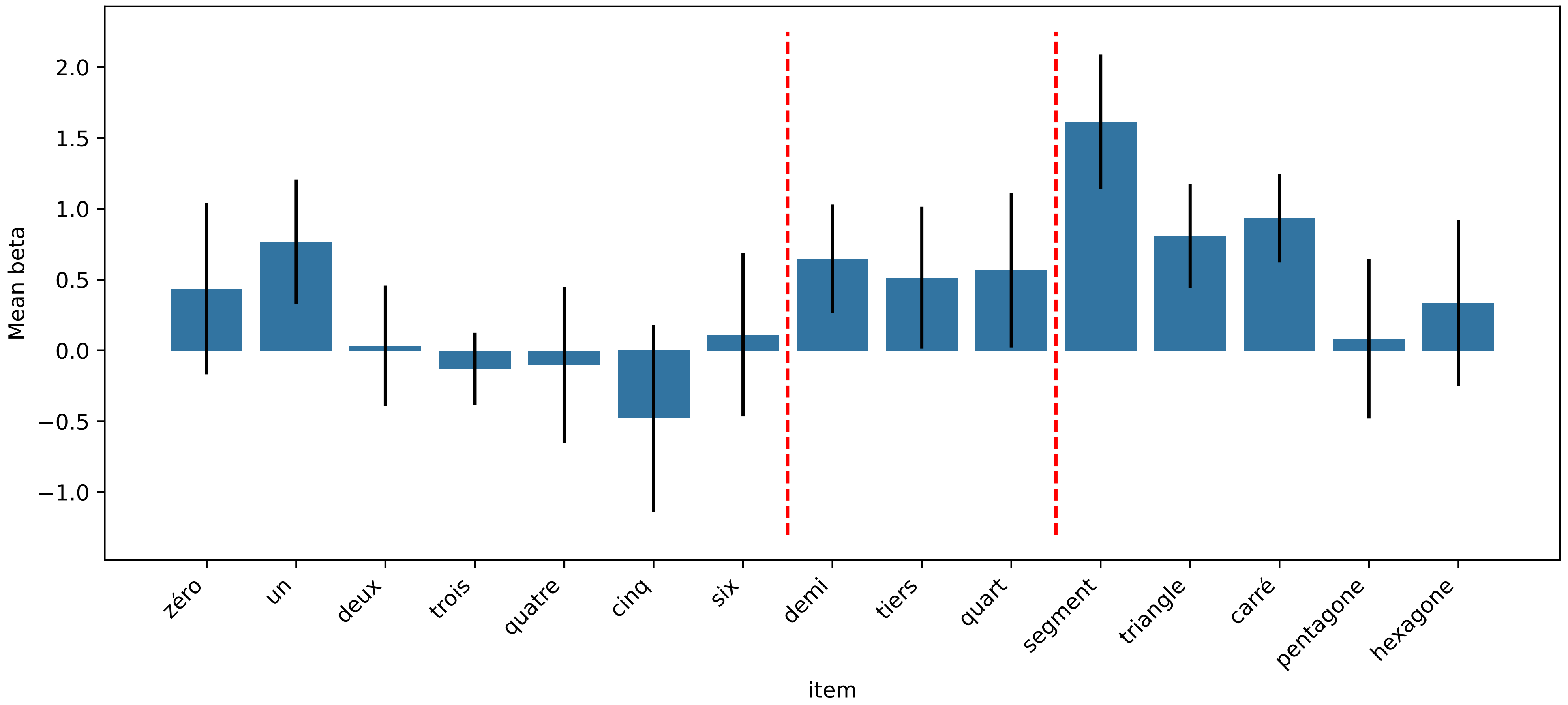

#### Region: Area_PH_L (geom > arith)

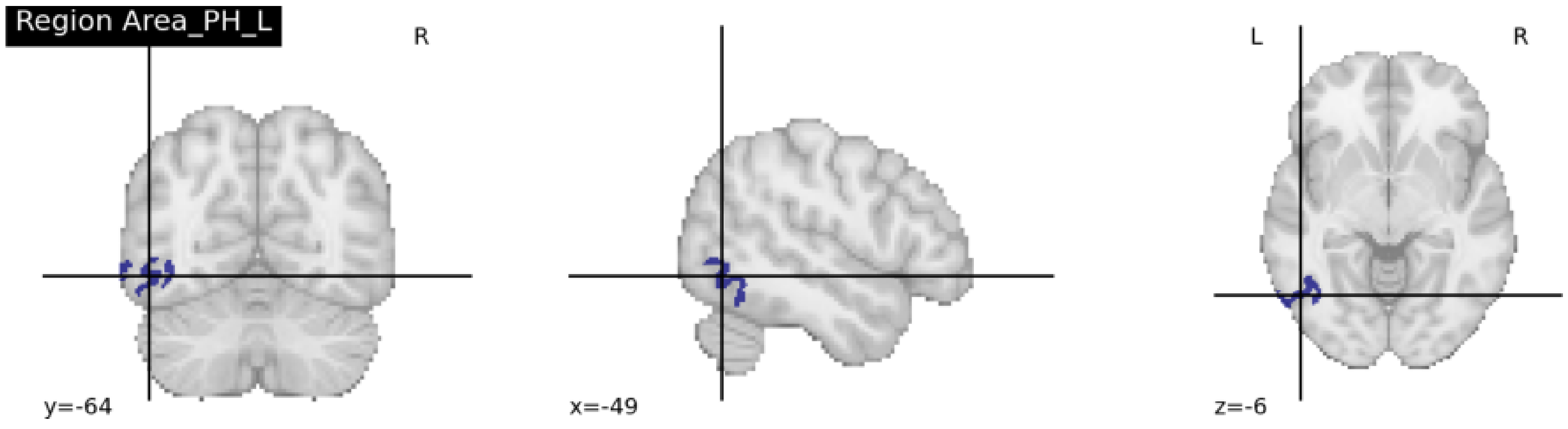


Average region size: 78 voxels (across 14 participants)
chi2-test between the two models: chi2(2) = 37.07, p = 2.414e-07 (FDR corrected)

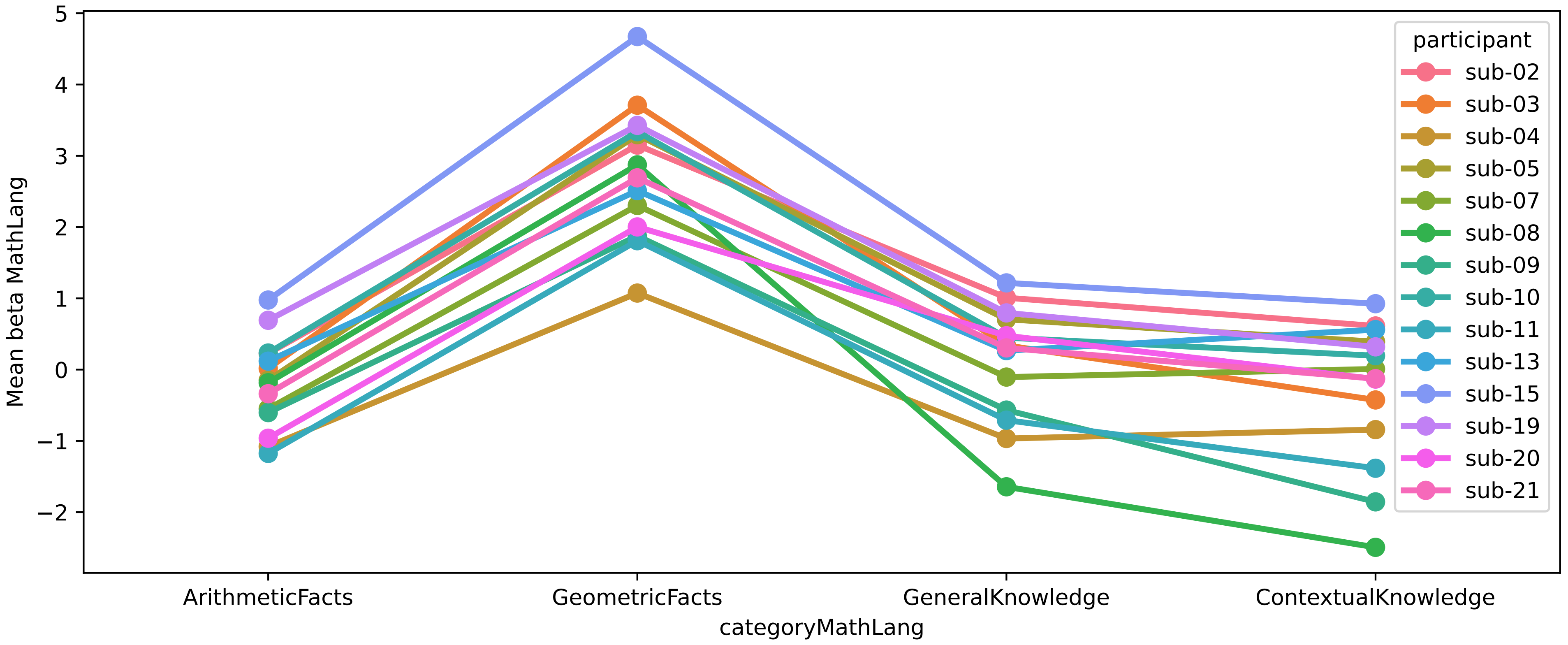


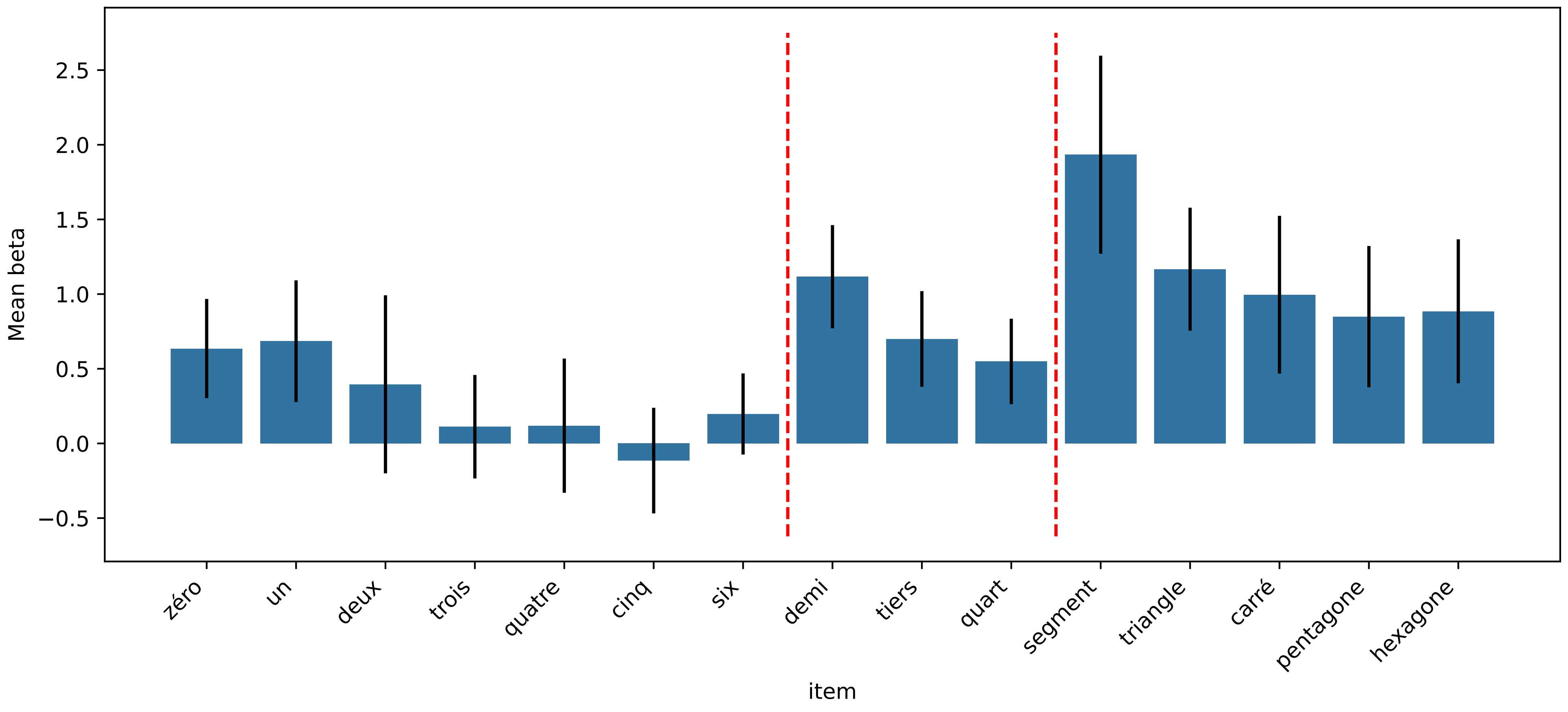

### Region: Area_TE1_posterior_L (geom > arith)

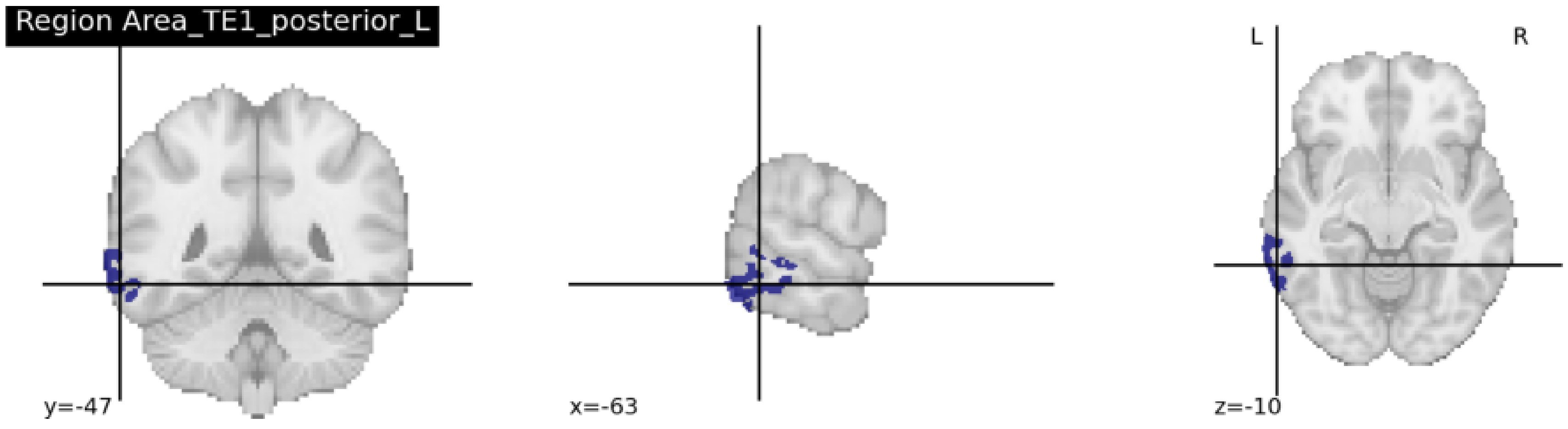


Average region size: 92 voxels (across 18 participants)
chi2-test between the two models: chi2(2) = 45.50, p = 7.110e-09 (FDR corrected)

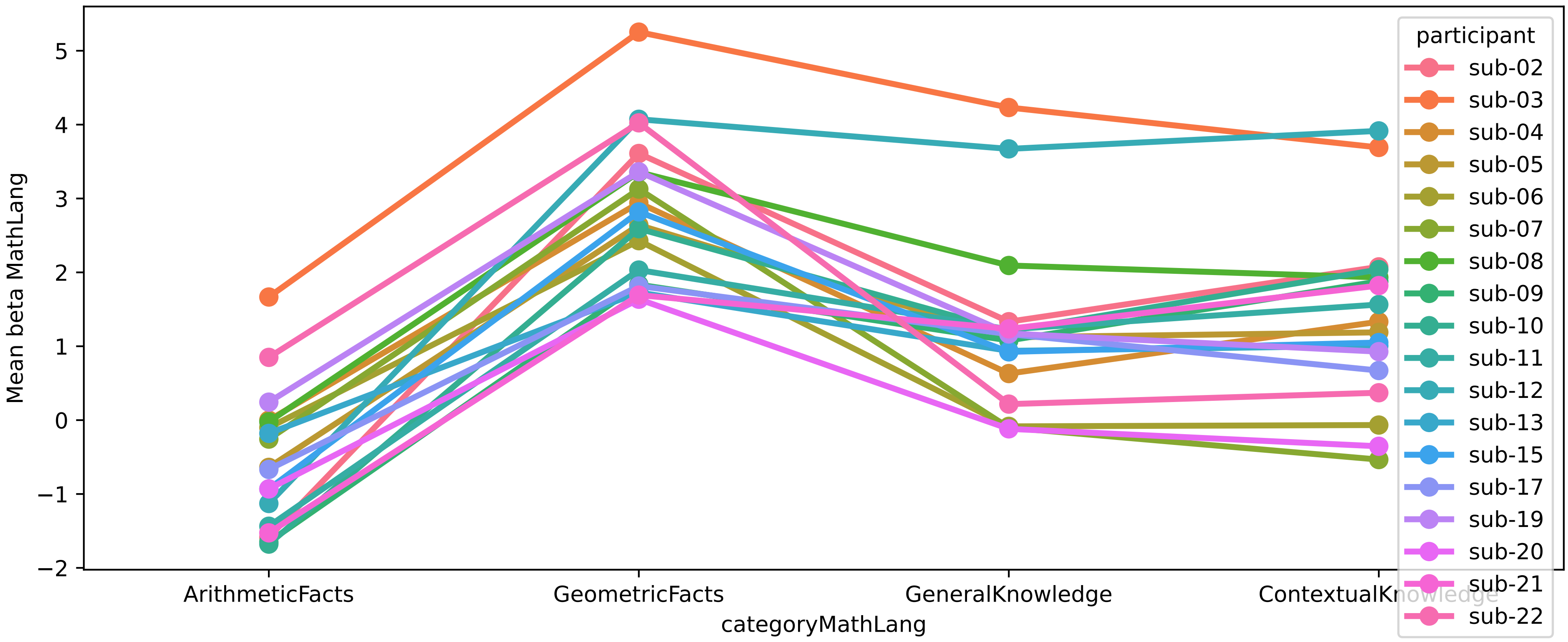


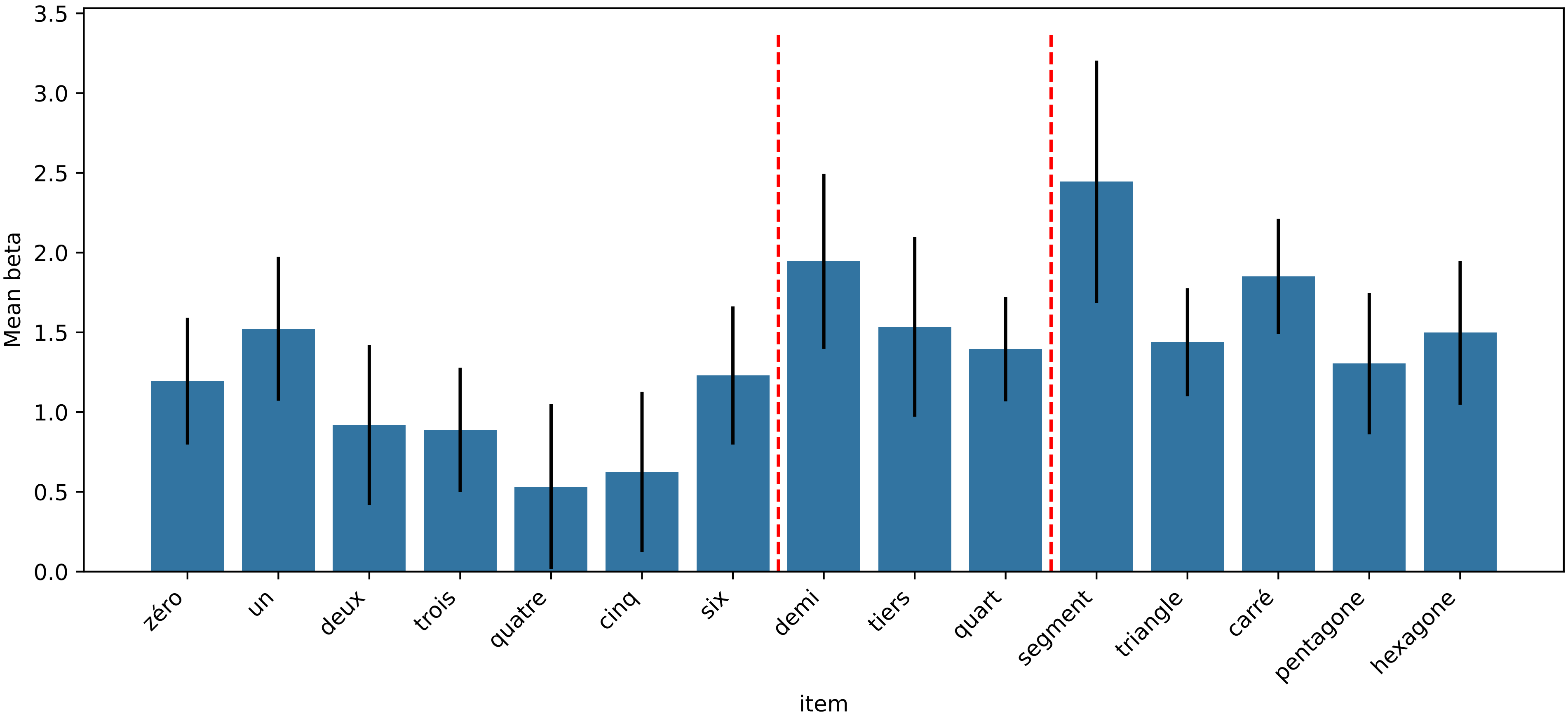

### Region: Area_TE2_posterior_L (geom > arith)

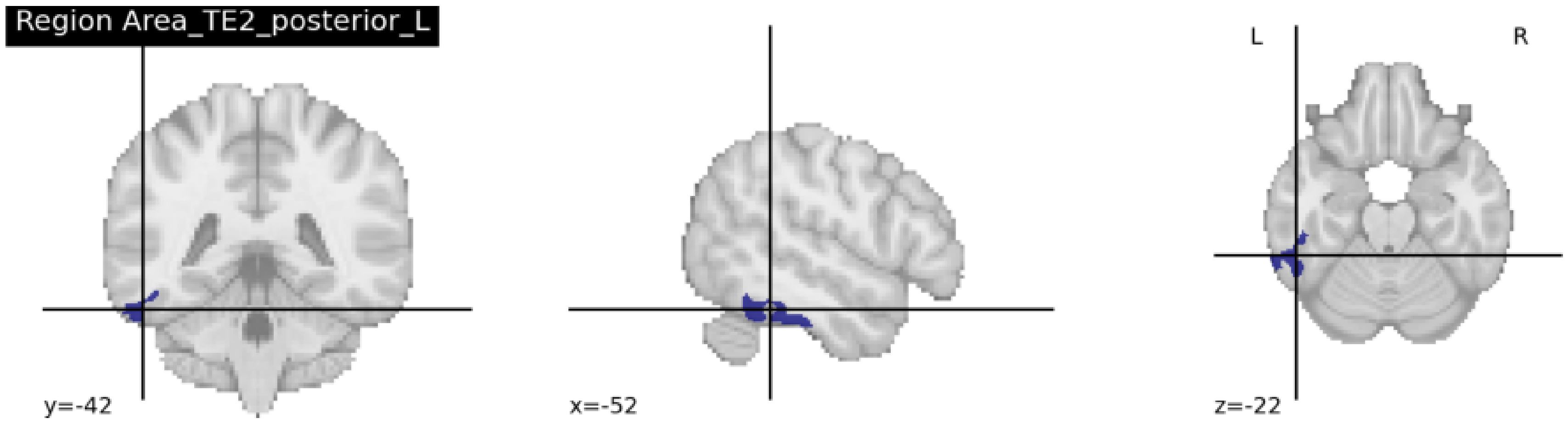


Average region size: 32 voxels (across 6 participants)
chi2-test between the two models: chi2(2) = 28.88, p = 7.221e-06 (FDR corrected)

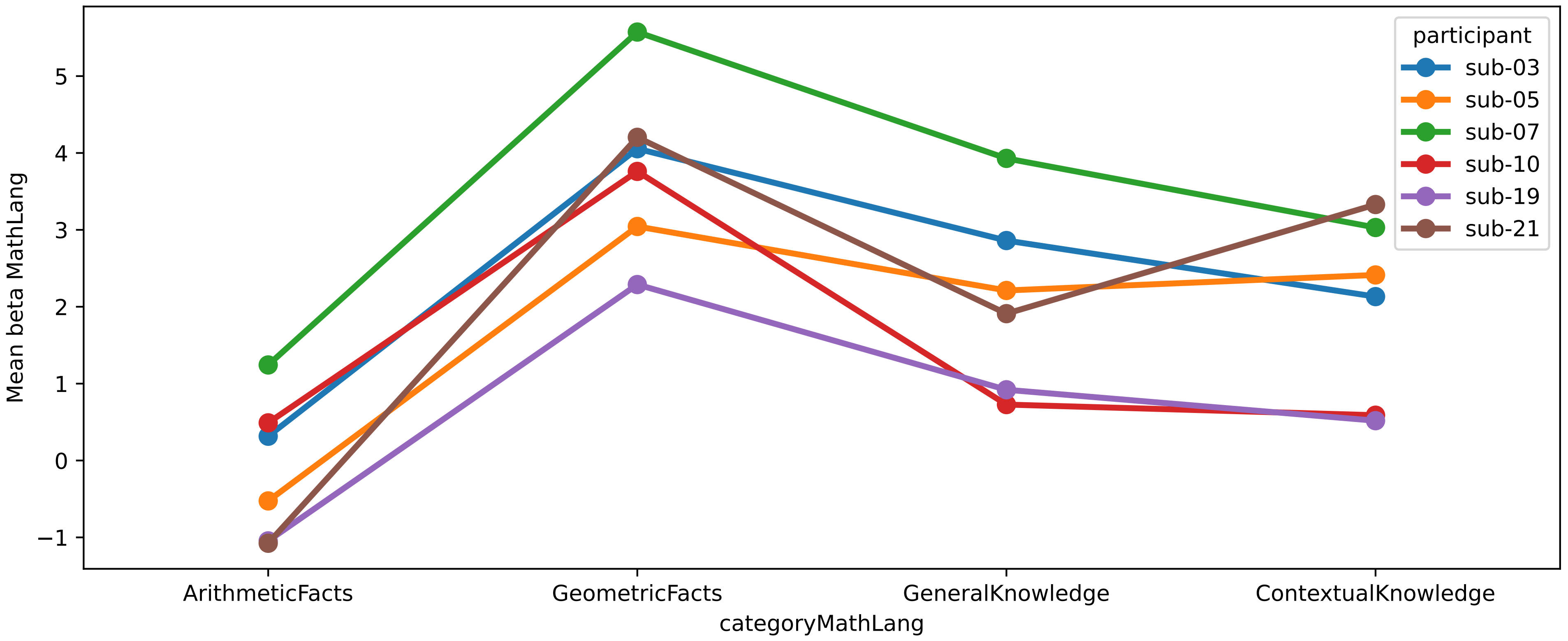


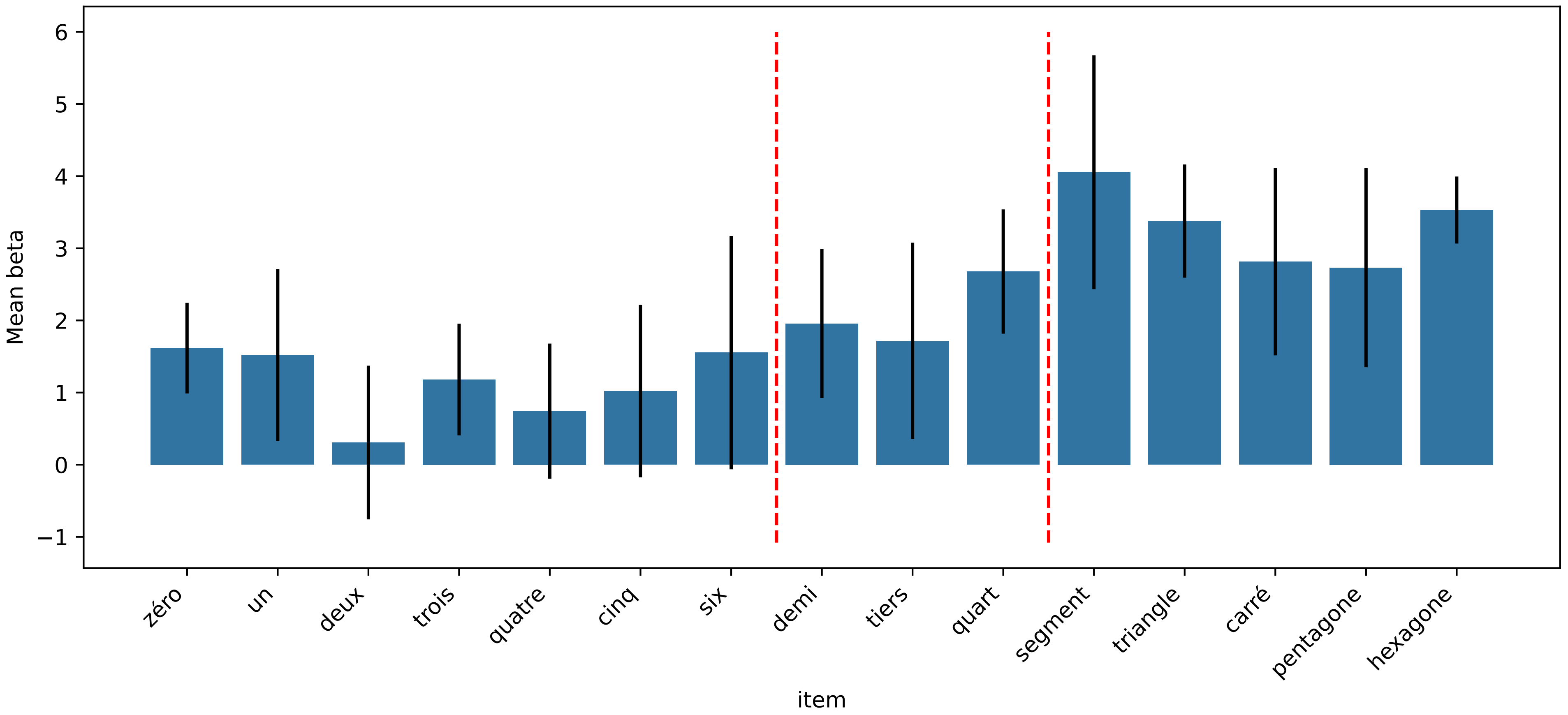

### Region: Fusiform_Face_Complex_L (geom > arith)

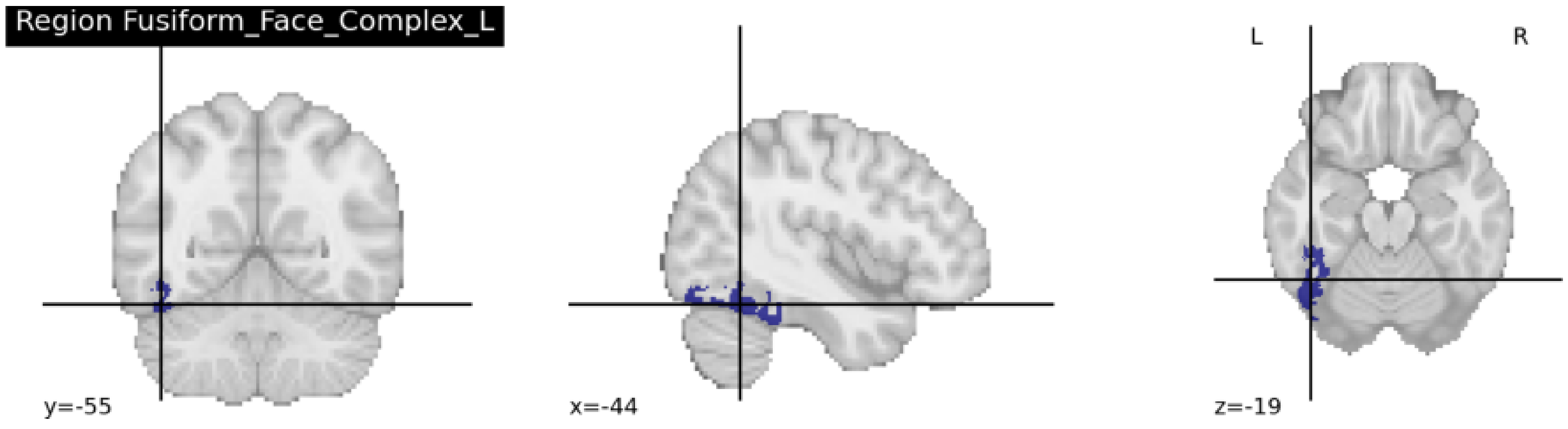


Average region size: 19 voxels (across 3 participants)
chi2-test between the two models: chi2(2) = 11.11, p = 2.085e-02 (FDR corrected)

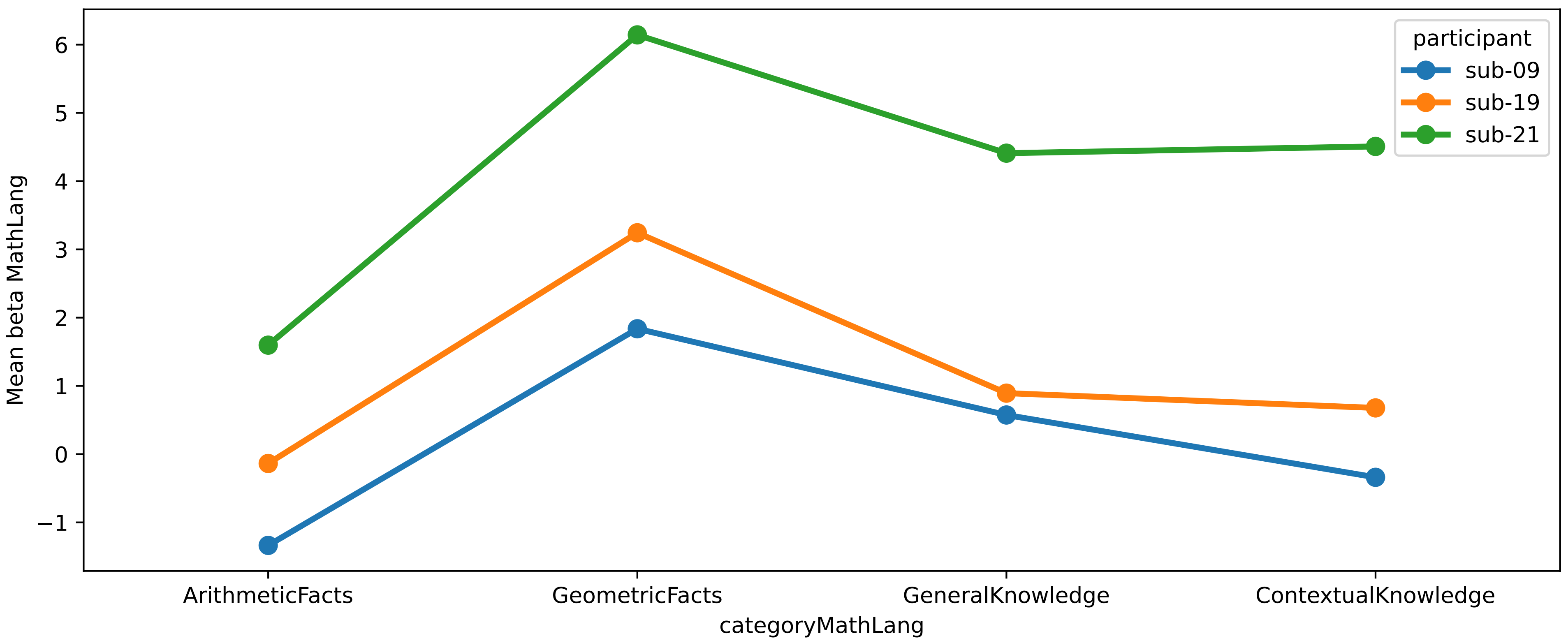


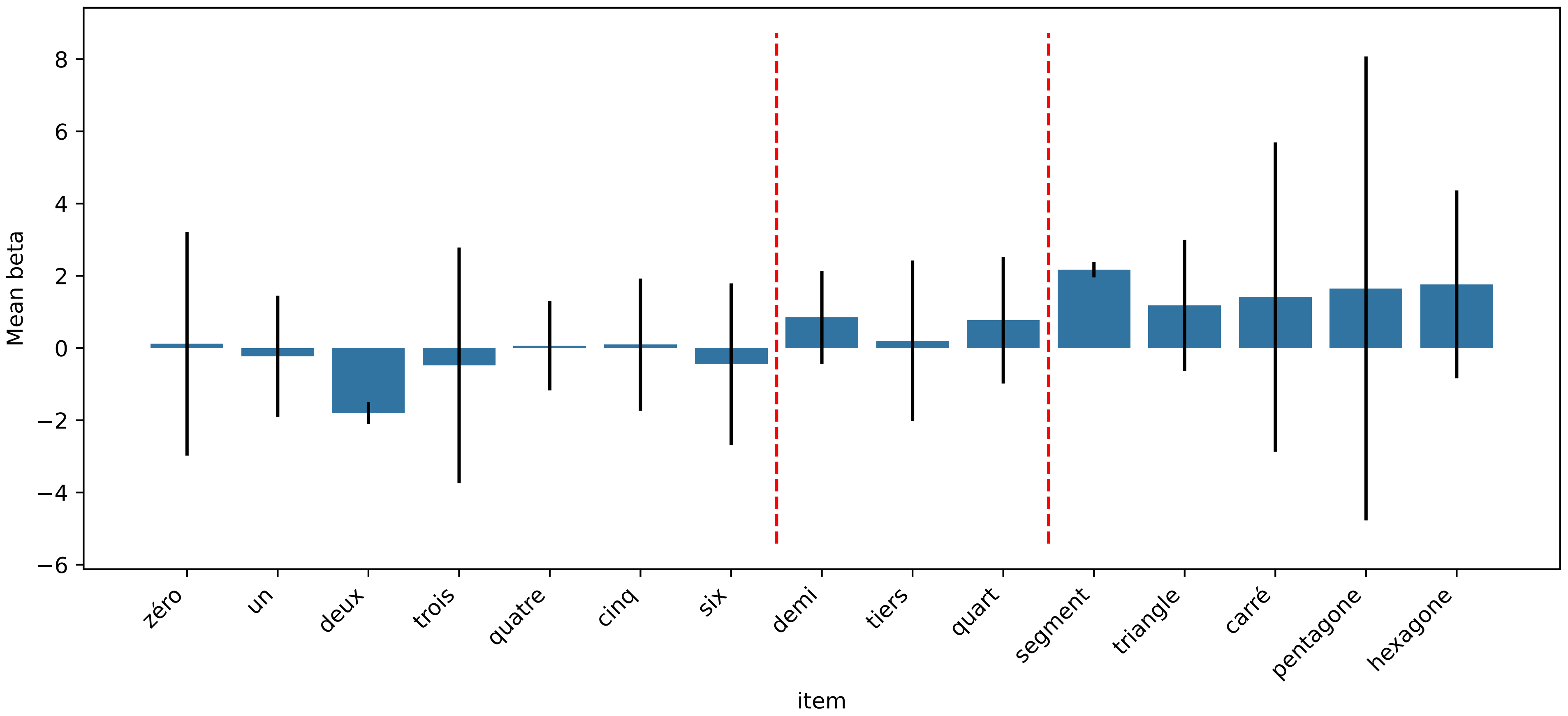

Region: Medial_IntraParietal_Area_L (geom > arith)

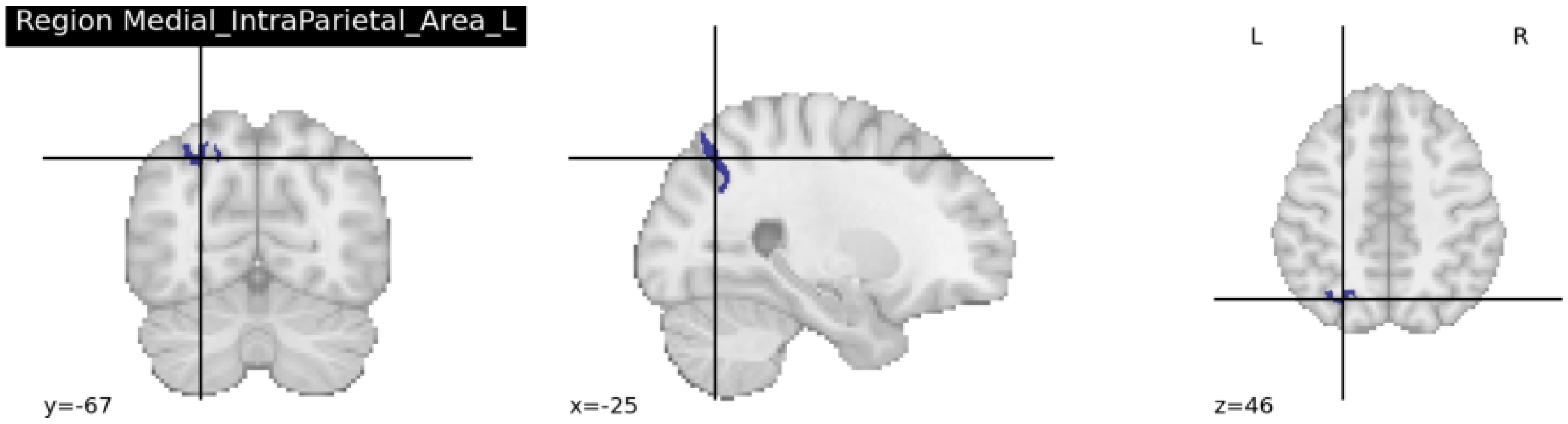


Average region size: 65 voxels (across 4 participants)
chi2-test between the two models: chi2(2) = 11.83, p = 1.824e-02 (FDR corrected)

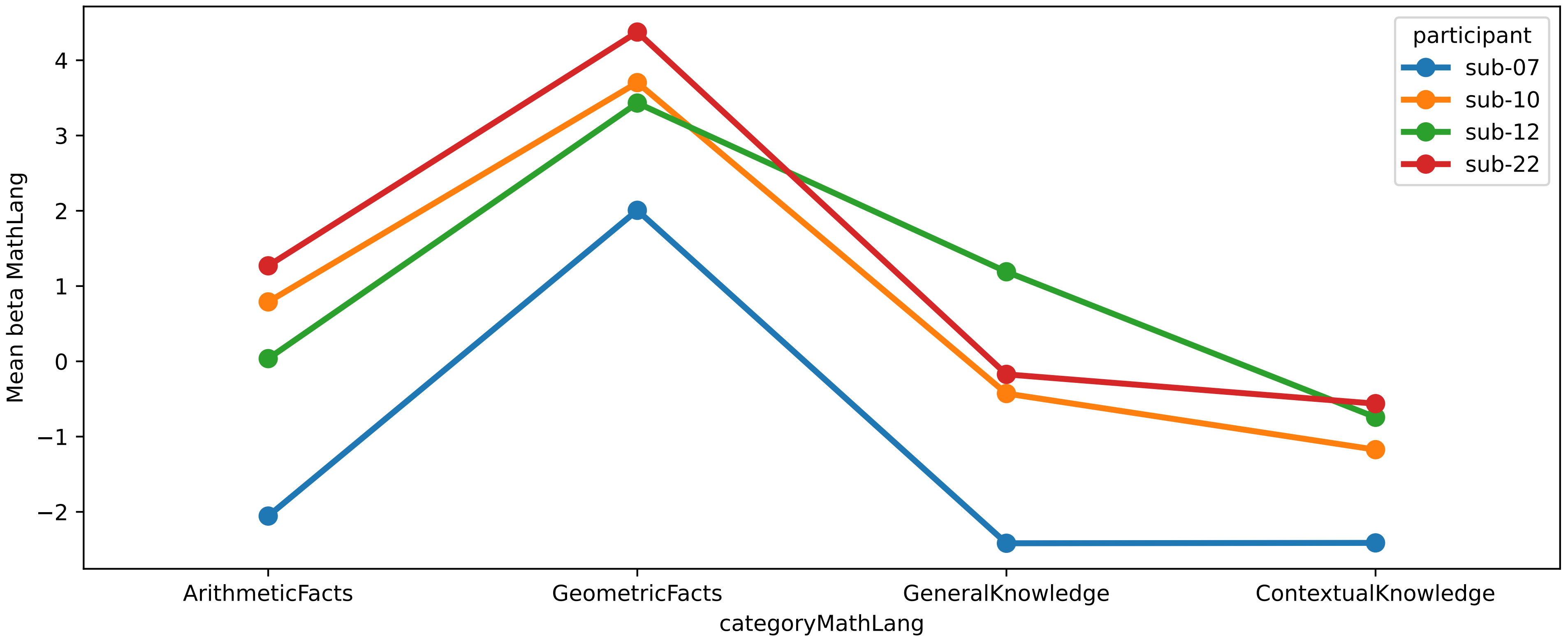


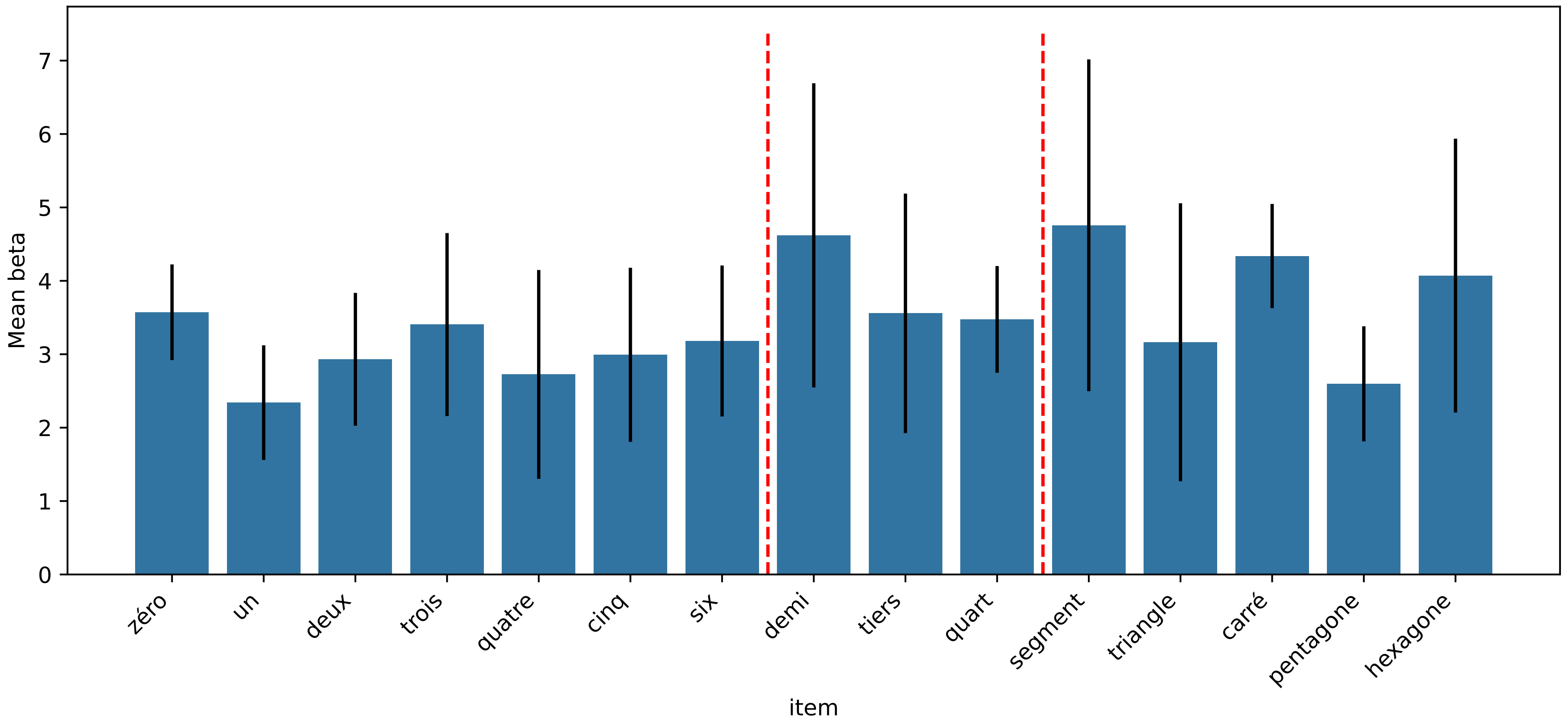
