## Supplementary material for "Cortical localization and dynamics of elementary mathematical concepts": S6

### List of HCP-MMP1.0 regions along bilateral IPS and ITG retained for subject specific ROI analyses

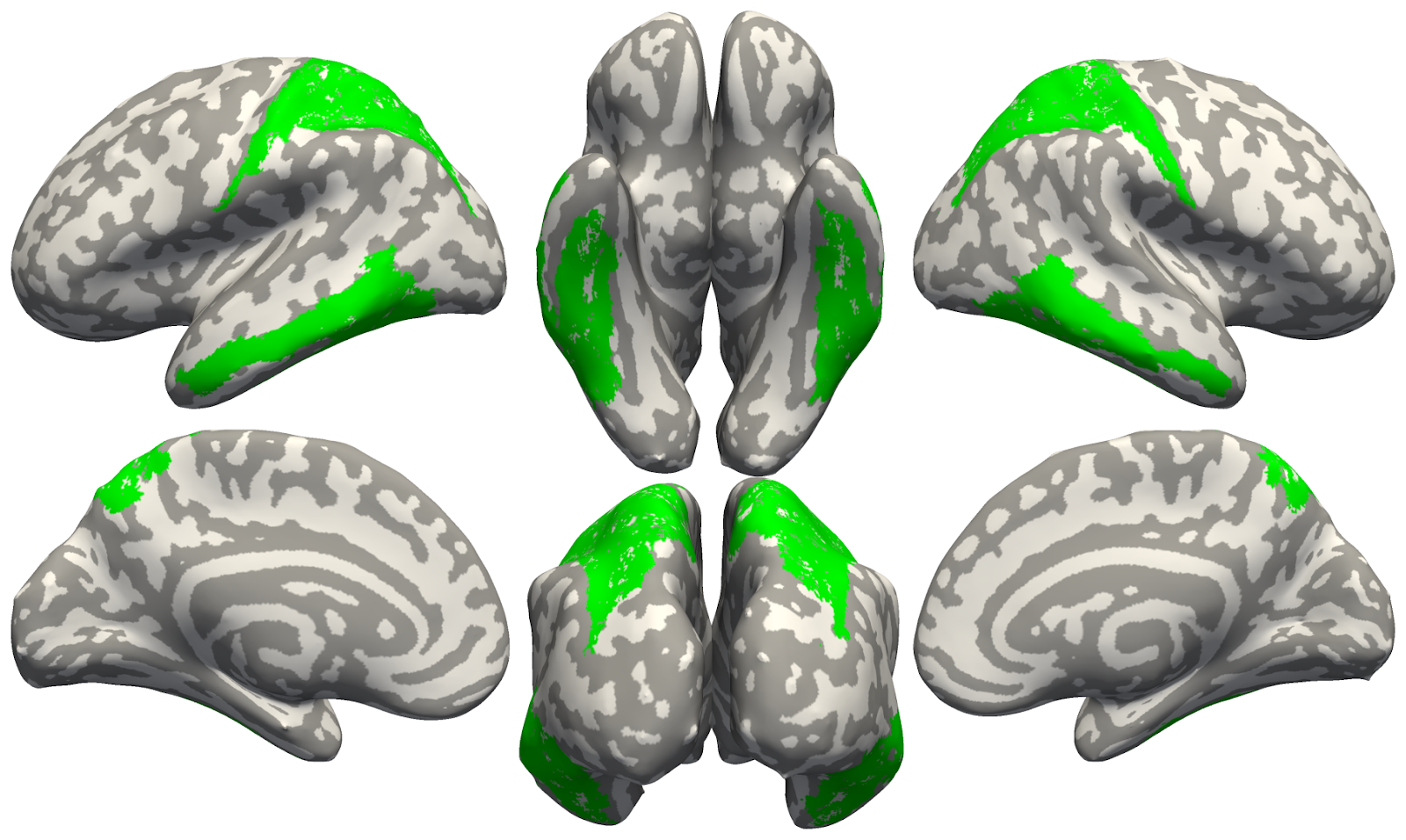

**Figure.** Surface view of the 48 atlas regions retained for the analyses.

#### List of regions

##### Bilateral IPS

| **region long name** | **region** | **lobe** | **cortex** |
| --- | --- | --- | --- |
| IntraParietal Sulcus Area 1 | IPS1 | Par | Dorsal Stream Visual |
| Medial Area 7P | 7Pm | Par | Superior Parietal |
| Lateral Area 7A | 7AL | Par | Superior Parietal |
| Medial Area 7A | 7Am | Par | Superior Parietal |
| Lateral Area 7P | 7Pl | Par | Superior Parietal |
| Area 7PC | 7PC | Par | Superior Parietal |
| Area Lateral IntraParietal ventral | LIPv | Par | Superior Parietal |
| Ventral IntraParietal Complex | VIP | Par | Superior Parietal |
| Medial IntraParietal Area | MIP | Par | Superior Parietal |
| Area 1 | 1 | Par | Somatosensory and Motor |
| Area 2 | 2 | Par | Somatosensory and Motor |
| Area Lateral IntraParietal dorsal | LIPd | Par | Superior Parietal |
| Anterior IntraParietal Area | AIP | Par | Superior Parietal |
| Area IntraParietal 2 | IP2 | Par | Inferior Parietal |
| Area IntraParietal 1 | IP1 | Par | Inferior Parietal |
| Area IntraParietal 0 | IP0 | Occ | Inferior Parietal |

##### Bilateral ITG

| **region long name** | **region** | **lobe** | **cortex** |
| --- | --- | --- | --- |
| Fusiform Face Complex | FFC | Temp | Ventral Stream Visual |
| Area TE1 anterior | TE1a | Temp | Lateral Temporal |
| Area TE1 posterior | TE1p | Temp | Lateral Temporal |
| Area TF | TF | Temp | Medial Temporal |
| Area TE2 posterior | TE2p | Temp | Lateral Temporal |
| Area PHT | PHT | Temp | Lateral Temporal |
| Area PH | PH | Temp | MT+ Complex and Neighboring Visual Areas |
| Area TE1 Middle | TE1m | Temp | Lateral Temporal |
